# TTF2 drives mitotic replisome disassembly and MiDAS by coupling the TRAIP ubiquitin ligase to Polε

**DOI:** 10.1101/2024.12.01.626218

**Authors:** Ryo Fujisawa, Karim P.M. Labib

## Abstract

Mammalian cells frequently enter mitosis before DNA replication has finished, necessitating the rapid processing of replication forks to facilitate chromosome segregation. The TRAIP ubiquitin ligase induces mitotic replisome disassembly, fork cleavage, and repair via ‘Mitotic DNA Synthesis’ (MiDAS). Until now, it was unclear how TRAIP is regulated in mitotic cells. Here we show that TRAIP phosphorylation mediates a complex with the TTF2 ATPase and DNA Polymerase ε (Polε). Whereas TTF2 ATPase activity removes RNA polymerase II from mitotic chromosomes, replisome disassembly involves an unanticipated mechanism. The TTF2 amino terminus couples TRAIP to Polε, via tandem Zinc fingers that recognise phosphorylated TRAIP, and a motif that binds to POLE2. Thereby, TTF2 and Polε cause TRAIP to ubiquitylate the CDC45-MCM-GINS (CMG) helicase, triggering replisome disassembly and MiDAS. These data identify TTF2 as a multifunctional regulator of chromatin transactions during mitosis.

---

When eukaryotic cells enter mitosis, any remaining regions of unreplicated DNA have the potential to impair the segregation of sister chromatids during anaphase, and thus represent a major threat to genome integrity. Cells duplicate the bulk of the genome during the S-phase of the cell cycle, by assembling replisomes at many origins of DNA replication, and the subsequent G2-phase was traditionally thought to separate the completion of replication from mitotic entry. In species with large genomes, however, mathematical modelling of origin spacing and replication errors indicated that the replication of a small subset of loci is likely to continue throughout G2-phase and even into mitosis (Al Mamun *et al*, 2016; Moreno *et al*, 2016). Correspondingly, mitotic entry before the completion of DNA replication is a feature of the early human embryo from the first cell cycle (Palmerola *et al*, 2022), and is especially prevalent in cancer cells with inherent DNA replication stress that results from oncogene activation (Bournaka *et al*, 2024). In such cases, cells still enter mitosis despite the persistence of a low number of DNA replication forks, which are not sufficient to activate cell cycle checkpoints. Therefore, eukaryotes with large genomes needed to evolve mechanisms to process sites of incomplete DNA replication very quickly during early mitosis, to support successful chromosome segregation during anaphase. At present, these pathways are poorly understood in all species.

One such mechanism involves the cleavage of DNA replication forks upon mitotic entry, followed by repair pathways such as MiDAS that help to produce two disentangled sister chromatids that are able to segregate during anaphase (Bhowmick *et al*, 2023). High-resolution mapping of MiDAS in human cells shows that it commonly occurs at very late replicating loci within large transcription units, corresponding to ‘common fragile sites’ that are frequently rearranged in human cancer (Ji *et al*, 2020; Macheret *et al*, 2020). A likely trigger for the cleavage of DNA replication forks in mitotic cells is disassembly of the CMG helicase, which forms the stable core of the replisome and encircles the DNA template for leading strand synthesis. Animal cells disassemble the CMG helicase at any remaining DNA replication forks upon entry into mitosis (Deng *et al*, 2019; Priego Moreno *et al*, 2019; Sonneville *et al*, 2019; Sonneville *et al*, 2017), thereby exposing the underlying parental DNA to nucleolytic attack. Work with *Xenopus laevis* egg extracts supports a model whereby mitotic CMG disassembly and leading strand cleavage at converging replisomes, followed by subsequent gap filling and Pol-theta dependent microhomology-mediated repair, would produce one daughter chromosome with a sister chromatid exchange and another with a small deletion (Deng *et al*., 2019). These features are typical of common fragile sites in human cells (Bhowmick *et al*., 2023; Bournaka *et al*., 2024).

In metazoa, the TRAIP ubiquitin ligase is responsible for CMG helicase ubiquitylation and disassembly upon mitotic entry, leading to replication fork cleavage and MiDAS (Deng *et al*., 2019; Priego Moreno *et al*., 2019; Sonneville *et al*., 2019). During S-phase, TRAIP associates with the replisome (Dewar *et al*, 2017; Villa *et al*, 2021) and specifically ubiquitylates factors in front of stalled replication forks, to initiate the repair of protein-DNA crosslinks (Larsen *et al*, 2019) or inter-strand DNA crosslinks (Wu *et al*, 2019), and to resolve transcription-replication conflicts (Scaramuzza *et al*, 2023). Upon entry into mitosis, however, TRAIP rapidly induces the ubiquitylation and disassembly of all remaining CMG helicase complexes (Deng *et al*., 2019; Priego Moreno *et al*., 2019; Sonneville *et al*., 2019; Villa *et al*., 2021). This does not require the convergence of two replisomes and is not impaired by replication fork DNA, unlike CMG ubiquitylation by the ubiquitin ligase CUL2^LRR1^, which is inhibited by the lagging strand DNA template at replication forks until the termination of DNA synthesis (Deegan *et al*, 2020; Jenkyn-Bedford *et al*, 2021; Low *et al*, 2020).

Until now, the mechanism by which TRAIP is activated to induce CMG helicase disassembly during mitosis was unknown. Here we show that mammalian TRAIP is regulated by the TTF2 ATPase in mitotic cells. TTF2 couples TRAIP to Polε, which binds to the CMG helicase at DNA replication forks and synthesises the nascent leading strand (Daigaku *et al*, 2015; Jenkyn-Bedford *et al*., 2021; Langston *et al*, 2014; Nick McElhinny *et al*, 2008; Pursell *et al*, 2007; Sengupta *et al*, 2013). These findings indicate that TTF2 and Polε position TRAIP to promote CMG helicase ubiquitylation and disassembly in mitotic cells, thereby initiating the rapid processing of sites of incomplete DNA replication.

## Results

### CDK phosphorylation of TRAIP is required for CMG helicase disassembly during mitosis

To study how TRAIP is regulated upon entry into mitosis, we used mouse embryonic stem cells (ES cells) that provide a tractable system for studying the assembly and disassembly of the mammalian replisome (Evrin *et al*, 2023; Villa *et al*., 2021). When mouse ES cells were synchronised in mitosis, by inhibition of the Eg5 kinesin with S-trityl-L-cysteine (STLC), the mobility of TRAIP was shifted in polyacrylamide gels (Figure 1A, compare lanes 4-5) that contained the phosphate-binding molecule known as ‘Phos-tag’ (Kinoshita *et al*, 2006). The migration of TRAIP in Phos-tag gels was further retarded when mouse ES cells were driven prematurely into mitosis by treatment with Calyculin A (Gotoh *et al*, 1995), which inhibits the protein phosphatases PP2A and PP1 that counteract mitotic protein kinases (Figure 1A, lane 6; Figure S1). These findings suggested that TRAIP is regulated by phosphorylation during mitosis.

**Figure 1.**
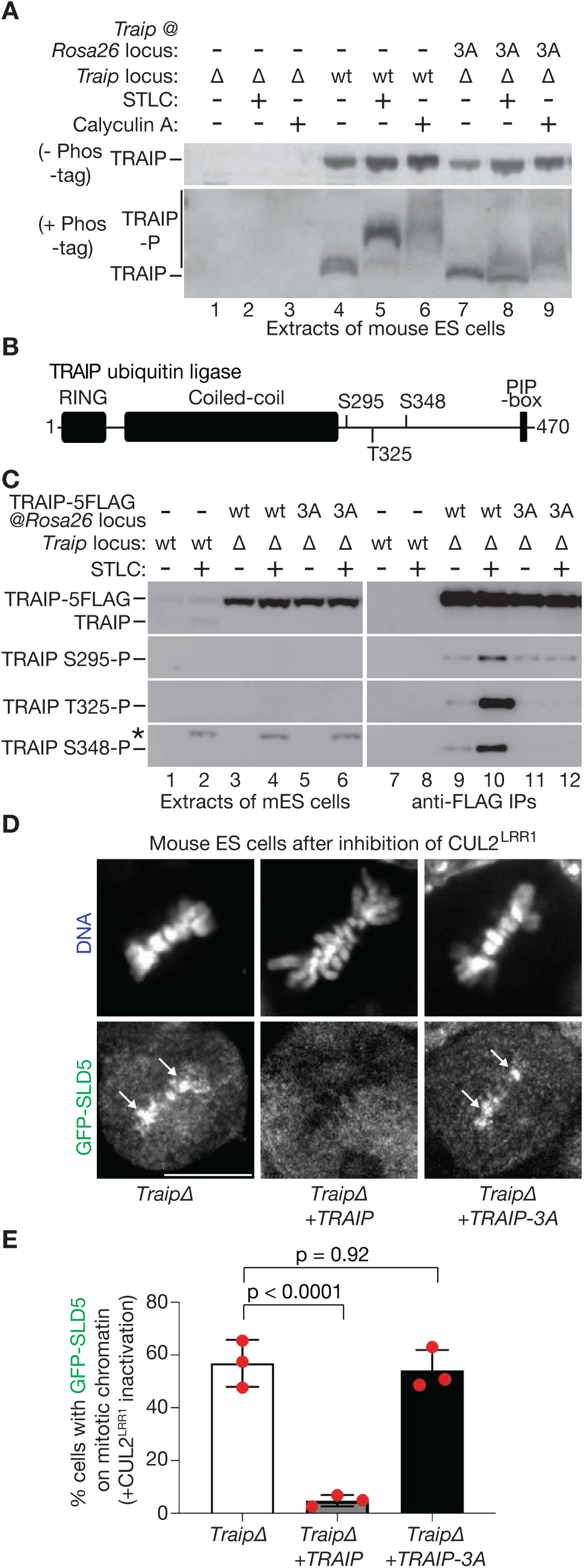
CDK phosphorylation of TRAIP is required for mitotic CMG helicase disassembly. (A) Mouse ES cells of the indicated genotypes were treated as shown with 5 µM S-trityl-L-cysteine (STLC) or 100 nM Calyculin A. The phosphorylation status of TRAIP was monitored via immunoblotting, following SDS-PAGE gel in the absence or presence of Phos-tag. (B) Domain structure of mouse TRAIP, including the location of three consensus phosphorylation sites for Cyclin-Dependent Kinase (CDK) within the disordered carboxy-terminal half of the protein (S295, T325 and S348). (C) Mouse ES cells of the indicated genotypes were treated as shown with STLC. Extracts were incubated with magnetic beads coated with anti-FLAG antibodies, to immunoprecipitate TRAIP-5FLAG (anti-FLAG IPs), before immunoblotting with anti-TRAIP antibodies or with phospho-specific antibodies corresponding to the three consensus CDK sites (S295-P, T325-P and S348-P). The asterisk indicates a non-specific band in the immunoblot for TRAIP-S348-P. (D) CUL2^LRR1^ activity was inhibited in mouse ES cells of the indicated genotypes, as described in Materials and Methods. The presence of the CMG helicase on mitotic chromosomes (white arrows) was monitored by spinning disk confocal microscopy, via GFP-SLD5 and Hoechst staining of DNA. The scale bar corresponds to 10 µm. (E) Quantification of the data from (D). The mean values for three independent experiments are shown as red spots, whereas the histograms show the average of the mean values, with associated standard deviations. P-values were calculated using a one-way ANOVA followed by Dunnett’s multiple-comparison test.

Mouse TRAIP contains three consensus sites for cyclin-dependent kinases (Figure 1B), which are broadly conserved in TRAIP orthologues from other vertebrate species (Figure S2A). Mutation of all three sites reduced the phosphorylation of TRAIP in mitotic cells (Figure 1A, TRAIP-3A), suggesting that cyclin-dependent kinases are a major source of mitotic TRAIP phosphorylation. Moreover, phospho-specific antibodies confirmed that all three of TRAIP’s CDK-consensus sites are phosphorylated during mitosis (Figure 1C, compare lanes 9-10 with lanes 11-12). To test whether phosphorylation of TRAIP on CDK-consensus sites is required for mitotic CMG helicase disassembly, we inhibited CUL2^LRR1^ to block CMG helicase disassembly during DNA replication termination (Dewar *et al*., 2017; Sonneville *et al*., 2019; Villa *et al*., 2021), and then used spinning disk confocal microscopy to monitor the association of a GFP-tagged CMG subunit (GFP-SLD5) with condensed mitotic chromosomes. GFP-SLD5 persisted on metaphase chromosomes in *TraipΔ* cells (Figure 1D-E, *TraipΔ*), as seen previously (Villa *et al*., 2021), and this could be rescued by expression of wild type TRAIP from the *Rosa26* locus (Figure 1D-E, *TraipΔ* + TRAIP). In contrast, expression of the TRAIP-3A mutant was unable to rescue the mitotic CMG disassembly defect of *TraipΔ* cells (Figure 1D-E, *TraipΔ* + TRAIP-3A), though TRAIP-3A retained other aspects of TRAIP function, since it rescued the sensitivity of *TraipΔ* cells to the DNA-damaging agent mitomycin-C and the PARP inhibitor Olaparib (Figure S2B). These data indicate that mitotic phosphorylation of TRAIP, likely by Cyclin-Dependent Kinase, is essential for mitotic disassembly of the CMG helicase.

### Mitotic phosphorylation of TRAIP induces complex assembly with the TTF2 ATPase and DNA polymerase ε

Alphafold (Abramson *et al*, 2024; Evans *et al*, 2022; Mirdita *et al*, 2022) predicts with high confidence that TRAIP forms an extended dimer via a long region of coiled coil, preceded at the amino terminus by the RING domain that couples TRAIP to ubiquitin conjugating enzymes, and followed at the carboxy-terminal side by an extensive disordered tail that comprises about 40% of the protein (Figure S3). The three CDK-consensus sites are located within the disordered tail (Figure 2A), suggesting that they might mediate the interaction of dimeric TRAIP with other factors.

**Figure 2.**
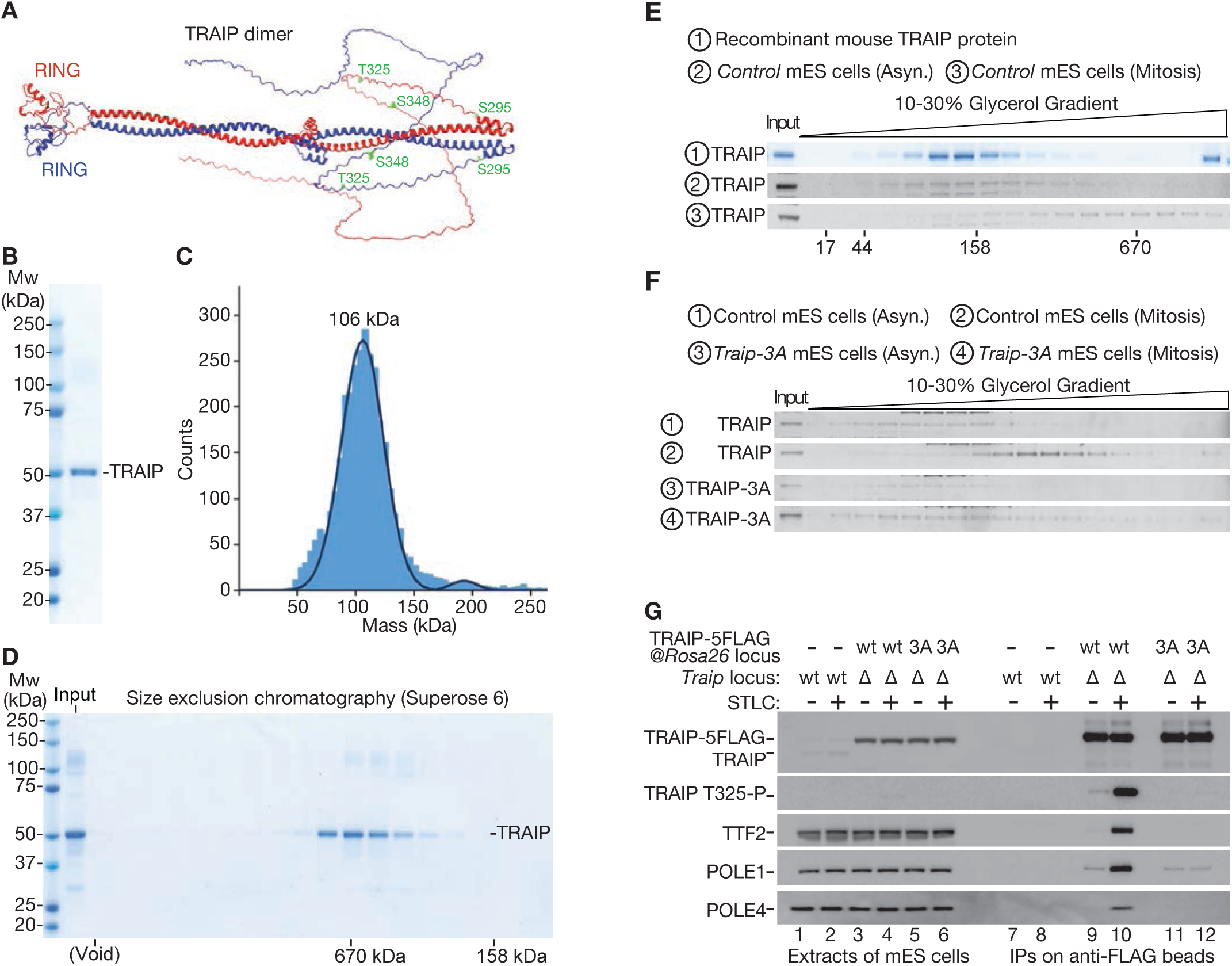
Mitotic phosphorylation of TRAIP drives complex formation with DNA polymerase epsilon and the TTF2 ATPase. (**A**) Alphafold3 prediction of dimeric TRAIP (also see Figure S3), indicating the location of the three mitotic CDK phosphorylation sites within the disordered C-terminal tail. (**B**) Recombinant mouse TRAIP was purified from *E. coli* and analysed by SDS-PAGE, before staining with Coomassie Brilliant blue. (**C**) Mass photometry analysis of recombinant mouse TRAIP (Mw = 53 kDa), indicating dimer formation (peak of 106 kDa). (**D**) Gel filtration analysis of recombinant mouse TRAIP, together with the indicated marker proteins (Thyroglobulin = 670 kDa, γ-globulin = 158 kDa). Fractions from the column were analysed by SDS-PAGE, before staining with Coomassie Brilliant blue. (**E**) Glycerol gradient fractionation of purified TRAIP (1) or extracts of mouse ES cells grown asynchronously (2; Asyn. = Asynchronous) or arrested in mitosis with 5 µM STLC (3). Fractions were analysed by SDS-PAGE and monitored by staining with Coomassie Brilliant blue (1) or by immunoblotting (2-3). Peak positions from equivalent gradients are indicated for a range of marker proteins (Thyroglobulin = 670 kDa, γ-globulin = 158 kDa, Ovalbumin = 44 kDa and Myoglobulin = 17 kDa). (**F**) Equivalent analysis comparing extracts of control mouse ES cells or *Traip-3A* cells grown asynchronously (1 + 3) or arrested in mitosis (2 + 4). (**G**) Cells of the indicated genotypes were treated as shown with 5 µM STLC. Extracts were then incubated with magnetic beads coated with anti-FLAG antibodies, and the proteins associated with TRAIP-5FLAG were analysed by immunoblotting with the indicated antibodies.

We expressed a recombinant form of mouse TRAIP in *E. coli* and used mass photometry to show that the purified protein (Figure 2B) had a mass of about 106 kDa (Figure 2C), confirming dimerization as predicted by AlphaFold (Mw TRAIP monomer = 53.1 kDa). Moreover, recombinant TRAIP migrated more slowly than expected in a size exclusion column (Figure 2D), but not in a glycerol gradient (Figure 2E, sample 1), consistent with the elongated nature of the predicted TRAIP dimer.

When an extract of asynchronous mouse ES cells was fractionated in a glycerol gradient, the migration of TRAIP was very similar to the recombinant TRAIP dimer (Figure 2E, sample 2). However, fractionation of mitotic cell extracts revealed a dramatic shift of TRAIP to an apparently larger protein complex (Figure 2E, sample 3), formation of which was impaired by mutation of the consensus CDK sites (Figure 2F, compare samples 2 and 4). These findings suggested that mitotic phosphorylation causes the TRAIP ubiquitin ligase to associate with other factors that might control its ability to ubiquitylate the CMG helicase at DNA replication forks.

To identify mitotic partners of the TRAIP ubiquitin ligase, we generated mouse ES cells in which the only form of TRAIP was modified by addition of five copies of the FLAG epitope, either at the carboxyl terminus of the protein, or between two poorly conserved residues in the disordered tail (Figure S4A-B; Figure S2A). Both tagged forms of TRAIP were able to rescue the sensitivity of *TraipΔ* cells to mitomycin C and Olaparib (Figure S4C), as well as supporting mitotic CMG helicase disassembly (Figure S4D-E). Subsequently, therefore, we isolated FLAG-tagged TRAIP from extracts of asynchronous or mitotic mouse ES cells, and used mass spectrometry to analyse co-purifying proteins. In three independent experiments (Tables S1-S3), FLAG-tagged TRAIP co-purified from asynchronous and mitotic samples with the DNA repair factor RAP80, consistent with previous work (Soo Lee *et al*, 2016). RAP80 is part of the BRCA1-A complex and we also found that FLAG-tagged TRAIP co-purified with additional BRCA1-A components, namely ABRAXAS1, BABAM2, BABAM1 and BRCC36 (Tables S1-S3).

Strikingly, two factors selectively co-purified with TRAIP during mitosis. The first was an ATPase known as TTF2 (Transcription Termination Factor 2), which was originally identified as a factor in *Drosophila melanogaster* and Humans that releases RNA Polymerase II and nascent transcripts from template DNA *in vitro* (Liu *et al*, 1998; Xie & Price, 1997; Xie & Price, 1996). Subsequent work showed that human TTF2 and its *Drosophila* orthologue known as ‘lodestar’ are predominantly cytoplasmic proteins, which associate with chromosomes during mitosis and are required to displace RNA polymerase II and nascent transcripts from chromatin (Carmo *et al*, 2023; Girdham & Glover, 1991; Jiang *et al*, 2004), by a mechanism that remains to be characterised. Live cell video microscopy indicated that mouse TTF2 is principally a cytoplasmic protein during interphase (Figure S5A), before associating with chromosomes throughout mitosis (Figure S5B).

The second factor that co-purified selectively with TRAIP during mitosis was Polε. The preferential association of TRAIP with both TTF2 and Polε during mitosis was confirmed by immunoblotting, using pulldowns of either TRAIP-5FLAG (Figure 2G, compare lanes 9-10) or 5FLAG-POLE1 (Figure S6, lane 12). Moreover, selective association of TRAIP with TTF2 and Polε in mitotic cells was dependent upon TRAIP’s CDK-consensus sites (Figure 2G, lanes 11-12; note that TRAIP-3A still retained a residual association with Polε). These findings indicate that phosphorylation of TRAIP causes the assembly of a stable complex with TTF2 and Polε, during mitosis in mammalian cells.

### Tandem Zinc finger domains at the TTF2 amino terminus are required for mitotic CMG disassembly but are dispensable for displacement of RNA polymerase II from mitotic chromatin

TTF2 contains a SWI2/SNF2 ATPase domain in the carboxy-terminal half of the protein (Figure 3A, Figure S7), which is thought to mediate the release of RNA polymerase and nascent transcripts from mitotic chromosomes (Carmo *et al*., 2023; Jiang *et al*., 2004). TTF2 also has a pair of zinc-fingers of unknown function at the amino terminus, connected by a long disordered region to the ATPase domain (Figure 3A, Figure S7). To explore whether TTF2 is important for mitotic CMG helicase disassembly, and then elucidate the underlying mechanism, we used CRISPR-Cas9 to generate a set of tools for studying TTF2 function in mouse ES cells.

**Figure 3.**
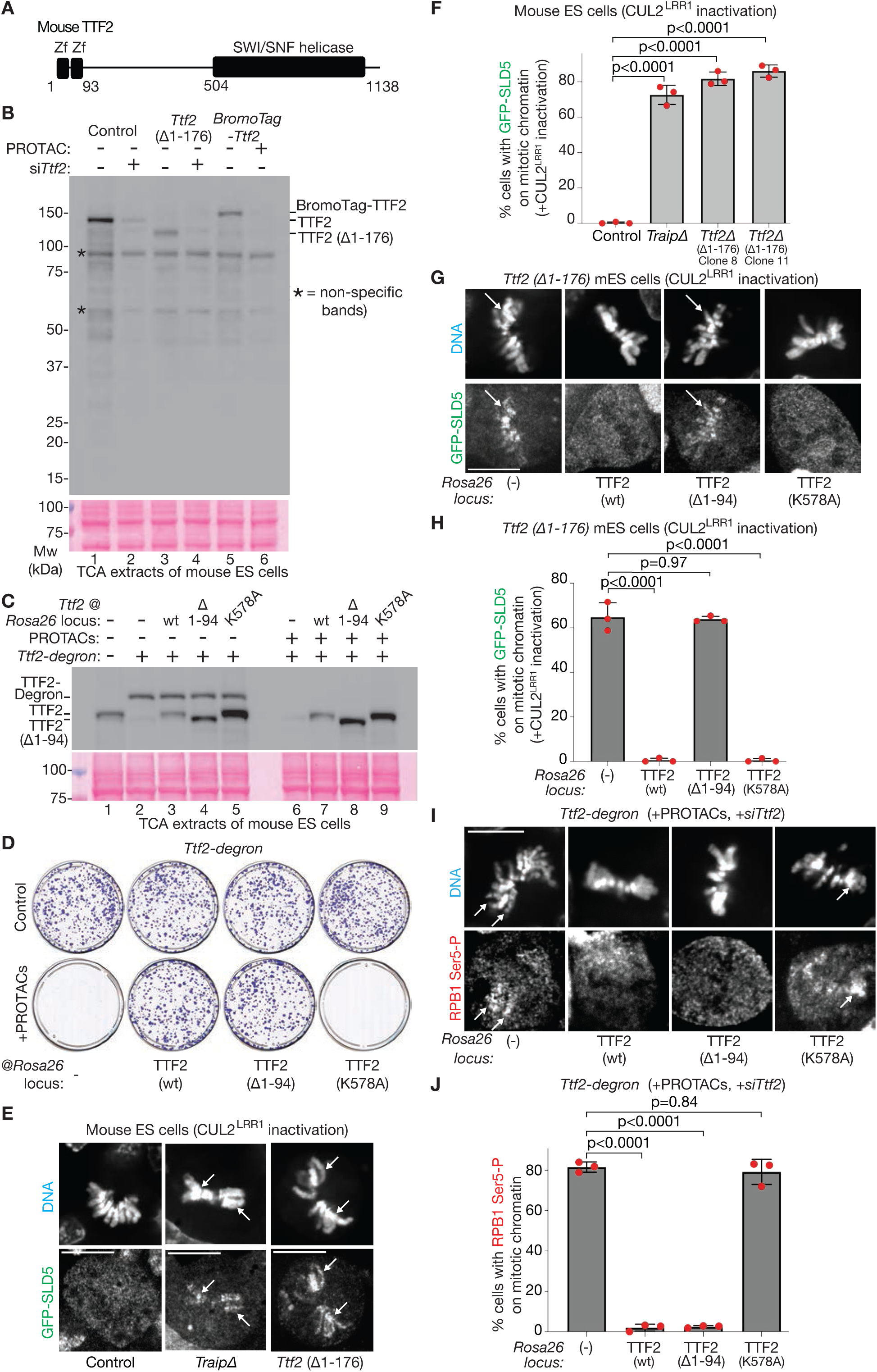
Tandem Zinc finger domains at the amino terminus of TTF2 are required for mitotic CMG disassembly but are dispensable for displacement of RNA polymerase II from mitotic chromatin. (A) Domain structure of mouse TTF2. Zf = Zinc finger. (B) Mouse ES cells of the indicated genotypes were treated as shown with 250 nM AGB1 (PROTAC) for two hours, or with siRNA specific to *Ttf2* (si*Ttf2*). Extracts were analysed by SDS-PAGE followed by immunoblotting (upper panel) or Ponceau S (lower panel). Also see Figures S8-S9. (C) Cells of the indicated genotypes (*Ttf2-degron* = *Ttf2-dTAG-BromoTag* – see Figure S9) were treated as shown with a combination of 250 nM AGB1 and 250 nM dTAG-13 (= ‘PROTACs’) for two hours. Extracts were analysed as above. (D) Colony formation assay with Ttf2-degron cells (*Ttf2-dTAG-BromoTag*) with the indicated TTF2 transgene expressed from the *Rosa26* locus. ‘PROTACs’ = 250 nM AGB1 + 250 nM dTAG-13. (E) CUL2^LRR1^ activity was inhibited in cells of the indicated genotypes, before detection of CMG on mitotic chromosomes (white arrows) as in Figure 1D. Scale bar = 10 µm. (F) Quantification of the data from (E), as in Figure 1E. (G) Cells of the indicated genotypes were analysed as in (E). Scale bar = 10 µm. (H) Quantification of the data from (G), as in Figure 1E. (I) Ttf2-degron cells (*Ttf2-dTAG-BromoTag*) were treated with *Ttf2* siRNA (si*Ttf2*), combined with addition of PROTACs (250 nM AGB1 + 250 nM dTAG-13) for the last 16 hours. The association of RPB1 with mitotic chromosomes was monitored by spinning disk confocal microscopy, using antibodies specific to phosphorylated Ser5 with the RPB1 C-terminal tail. Scale bar = 10 µm. (J) Quantification of the data from (I), as in Figure 1E.

Firstly, we used the repair of dual Cas9 nicking sites to generate small deletions within Exon 2 of the *Ttf2* gene, close to the start of the coding sequence, or in Exon 11 within the region encoding the ATPase domain (Figure S8A-C). Cell viability was reduced dramatically when the *Ttf2* coding sequence was cut within Exon 11 (Figure S8D), consistent with data from large-scale CRISPR screens in human cancer cell lines that indicate that TTF2 is essential for cell proliferation (Tsherniak *et al*, 2017). However, targeting Cas9 to Exon 2 of *Ttf2* did not compromise viability (Figure S8D), despite causing frame-shift deletions after the first 10-12 amino acids of the *Ttf2* coding sequence (Figure S8E), which prevented expression of full-length protein (Figure S8F). Such deletions within Exon 2 of *Ttf2* were associated with expression of a truncated protein that lacked the amino terminus (Figure S8F-G) and likely started from Met177, which represents the next in-frame methionine after the initiator ATG (Figure 3B shows that the truncated TTF2 protein was sensitive to *Ttf2* siRNAs, confirming specificity of the observed band). These data indicated that cells are able to proliferate in the absence of the amino-terminal pair of Zinc fingers of TTF2, and suggested that the ATPase domain might be essential for cell viability.

Secondly, we modified the *Ttf2* locus in mouse ES cells to introduce degrons that allow the rapid and conditional depletion of TTF2 protein. Initially, we used BromoTag that is ubiquitylated by the CUL2^VHL^ ubiquitin ligase (Bond *et al*, 2021), when cells are treated with the PROTAC (Proteolysis Targeting Chimera) known as AGB1 (Figure S9A-B, Figure 3A lanes 5-6). Despite depletion of the bulk of BromoTag-TTF2 in the presence of AGB1, cell proliferation was impaired but not completely blocked, likely due to residual protein that still provides some measure of TTF2 function. Therefore, we generated cells in which TTF2 was fused not only to BromoTag but also to dTAG (Nabet *et al*, 2018), which is ubiquitylated by CRL4^CRBN^ in response to the PROTAC dTag-13. Depletion of TTF2-dTAG-BromoTag in the presence of either AGB1 or dTag-13 (Figure S9B) partially reduced the growth of *Ttf2-dTAG-BromoTag* cells (Figure S9C), whereas combination of both PROTACs blocked cell proliferation more efficiently without affecting growth of control cells (Figure S9C).

The growth defect caused by efficient depletion of TTF2 could be rescued by expression of wild type TTF2 from the *Rosa26* locus (Figure 3C-D, wt TTF2). Similarly, expression of truncated TTF2 lacking the amino-terminal Zinc fingers also rescued depletion of degron-tagged TTF2 (Figure 3C-D, TTF2-Δ1-94). In contrast, TTF2 with a mutated Walker A motif (Figure 3C-D, TTF2-K578A) lacked ATPase activity (Figure S10) and was unable to support cell proliferation. These data confirmed that cells are able to proliferate in the absence of the TTF2 Zinc fingers and indicated that the ATPase activity of TTF2 is essential for cell viability.

Using these tools, we tested whether the TTF2 Zinc fingers and ATPase activity are required for mitotic CMG disassembly, using spinning disk confocal microscopy as above. Strikingly, cells lacking the amino terminus of TTF2 were just as defective in mitotic CMG disassembly as *TraipΔ* cells (Figure 3E-F). This defect could be rescued by expression of wild type TTF2 from the *Rosa26* locus (Figure 3G-H, TTF2 1-1138), but not by expression of TTF2 lacking the two Zinc fingers (Figure 3G-H, TTF2 95-1138). Interestingly, the TTF2-K578A allele that lacks ATPase activity (Figure S10) supported mitotic CMG disassembly as efficiently as wild type TTF2 (Figure 3G-H, TTF2-K578A). These data indicated that the TTF2 Zinc fingers are essential for mitotic CMG disassembly, whereas TTF2 ATPase activity is dispensable. Correspondingly, the amino terminal half of TTF2 that precedes the ATPase domain is sufficient to mediate complex assembly with TRAIP and Polε in mitotic cells (Figure S11, compare lanes 6 and 10). Similarly, the amino terminal half of TRAIP mediates mitotic CMG disassembly (Figure S11B), dependent upon the TTF2 Zinc fingers (Figure S11A-B).

These data imply that TTF2 contributes to mitotic CMG disassembly by a mechanism that is different from how TTF2 displaces RNA polymerase from mitotic chromosomes. To test whether the TTF2 Zinc fingers and ATPase activity are required to unload RNA polymerase from mitotic chromosomes, we monitored active RNA polymerase II in mouse ES cells during mitosis by spinning disk confocal microscopy, using antibodies that recognise phosphorylated Ser5 within the carboxy-terminal repeats of the RPB1 catalytic subunit (Figure S11C). Consistent with previous studies of human cells (Jiang *et al*., 2004; Parsons & Spencer, 1997), RNA polymerase II was excluded from chromosomes during mitosis in mouse ES cells (Figure S11C-D, Control). Similarly, RNA polymerase II was excluded from mitotic chromosomes in cells lacking either TRAIP or the amino terminus of TTF2 (Figure S11C-D, *TraipΔ* and *Ttf2(Δ1-176)*). In contrast, efficient depletion of degron-tagged TTF2 led to the accumulation of RNA polymerase II on mitotic chromosomes (Figure S11C-D and Figure 3G-H). This phenotype could be rescued by expression at the *Rosa26* locus of wild type TTF2, or TTF2 lacking the Zinc finger domains, but not by TTF2-K578A that lacks ATPase activity (Figure 3G-H). These findings indicate that displacement of RNA polymerase II from mitotic chromosomes requires the ATPase activity of TTF2, consistent with *in vitro* studies of human TTF2 and *Drosophila* lodestar (Liu *et al*., 1998), but is independent of the TTF2 Zinc fingers and does not require the TRAIP ubiquitin ligase.

### The TTF2 Zinc fingers are a receptor for CDK-phosphorylated T325 of TRAIP

Although Zinc finger domains have not previously been found to recognise phosphorylated proteins, the above data suggested that the TTF2 Zinc fingers might function as a receptor for phosphorylated TRAIP or Polε during mitosis. To explore this idea further, we first tested the impact on mitotic CMG disassembly of individually mutating the three consensus CDK sites in TRAIP. Whereas CMG was still unloaded from chromosomes in mitotic cells lacking either TRAIP-S295 or TRAIP-S348, mutation of TRAIP-T325 blocked mitotic CMG disassembly (Figure 4A). Correspondingly, mutation of TRAIP-T325 blocked the association of TRAIP with TTF2 and Polε in mitotic cells (Figure 4B, compare lanes 10 and 14).

**Figure 4.**
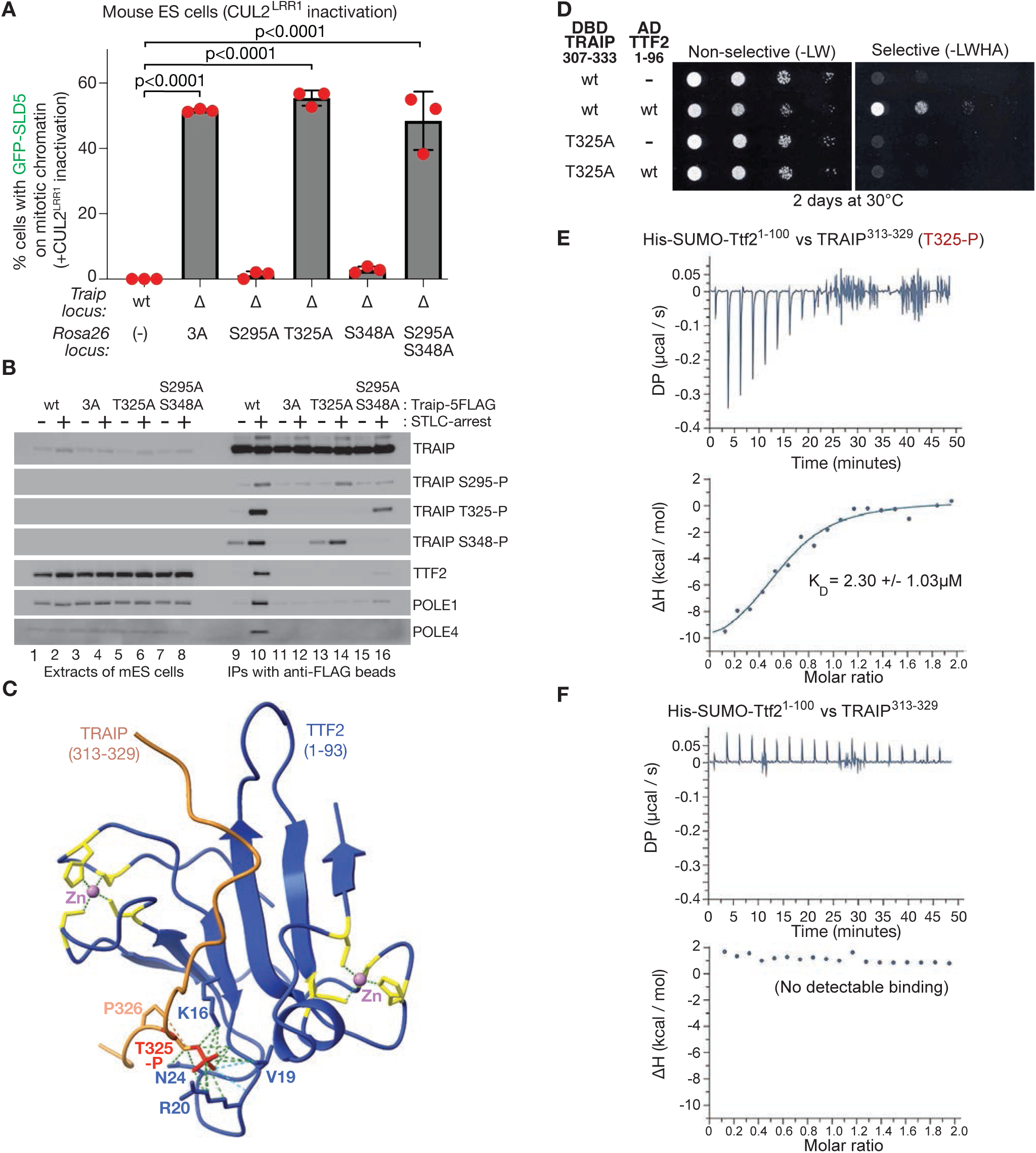
The TTF2 Zinc fingers are a receptor for CDK-phosphorylated TRAIP-T325. (**A**) The association of CMG with mitotic chromatin was analysed and quantified as in Figures 1+3. (**B**) *TraipΔ* cells expressing the indicated versions of TRAIP-5FLAG from the *Rosa26* locus were treated as shown with 5 µM STLC. FLAG-tagged TRAIP was isolated by immunoprecipitation from cell extracts, before immunoblotting of the indicated factors or phosphorylated forms of TRAIP. (**C**) AlphaFold 3 prediction of TTF2 1-93 bound to TRAIP 313-329 (containing phosphorylated TRAIP-T325). Contacts between phosphorylated TRAIP-T325 and TTF2 are shown as dashed lines in green, with hydrogen bonds shown in cyan. ‘Zn’ = Zinc ion. Also see Figure S13. (**D**) Yeast two-hybrid assay involving budding yeast cells expressing the indicated versions of TRAIP 307-333 fused to the Gal4 DNA-binding domain (DBD-TRAIP 307-333) and TTF2 1-96 fused to the Gal4 activation domain (AD-TTF2 1-96). Cells were grown on non-selective medium lacking leucine and tryptophan (-LW) or selective medium additionally lacking histidine and adenine (-LWHA). (**E**) The interaction of His-SUMO-tagged TTF2 with a peptide corresponding to TRAIP 313-329 (with phosphorylated T325) was monitored by isothermal titration calorimetry. The top panel shows the power differential (DP) during the experiment, whereas the bottom panel plots the enthalpy (ΔH) against the molar ratio of ligand and binding partner. K_D_ = dissociation constant. (**F**) Equivalent analysis for the combination of His-SUMO-tagged TTF2 with unphosphorylated peptide corresponding to TRAIP 313-329.

Interestingly, simultaneous mutation of TRAIP-S295 and TRAIP-S348 also blocked mitotic CMG disassembly (Figure 4A), impaired complex formation with TTF2 and Polε, and reduced the phosphorylation of TRAIP-T325 about five-fold (Figure 4B, lane 16). These findings indicated that phosphorylation of TRAIP-T325 is essential for mitotic CMG disassembly, suggesting that the region around T325 is a potential binding site for TTF2 or Polε. The data further suggest that phosphorylation of TRAIP-S295 and TRAIP-S348 might contribute indirectly to mitotic CMG disassembly, by promoting the efficient phosphorylation of TRAIP-T325.

The first of the TTF2 Zinc fingers resembles the ‘GRF’ Zinc fingers of DNA repair factors such as APE2 and NEIL3, which bind to single-strand DNA (Rodriguez *et al*, 2020; Wallace *et al*, 2017; Wu *et al*., 2019). However, we were unable to detect binding of the amino-terminal Zinc fingers of TTF2 to either single-strand or double-strand DNA in vitro, although the remainder of TTF2 did bind DNA under the same experimental conditions (Figure S12). This suggested that the the TTF2 Zinc fingers might have evolved to mediate protein-protein interactions instead of binding to DNA.

Importantly, AlphaFold 3 (Abramson *et al*., 2024) predicted that the two TTF2 Zinc fingers bind to residues 315-328 within the disordered tail of TRAIP in the presence of phosphorylated TRAIP-T325, with the phosphate being coordinated by conserved basic residues in the first of the TTF2 Zinc Fingers (Figure 4C; Figure S13 shows that the interaction is predicted with high confidence in all five AlphaFold models). To test the predicted interaction, we initially utilised the yeast two-hybrid assay, in which TRAIP 307-333 was found to interact with TTF2 1-96, dependent upon TRAIP-T325 (Figure 4D). Subsequently, we used Isothermal Titration Calorimetry to show that a recombinant form of TTF2 1-100 (Figure S12A) bound directly to a peptide corresponding to TRAIP 313-329, dependent upon phosphorylation of TRAIP-T325 (Figure 4E-F). These findings demonstrate that the TTF2 Zinc fingers comprise a new class of phospho-binding domain that functions as a receptor during mitosis for TRAIP, upon phosphorylation of TRAIP-T325.

### An amino terminal fragment of TTF2, comprising the Zinc fingers and a binding site for Polε, is sufficient for mitotic CMG unloading

When cells enter mitosis, TRAIP associates not only with TTF2 but also with Polε (Figure 2G), dependent upon the TTF2 Zinc fingers (Figure S11A, lanes 6+8 and lanes 10+12) and also upon phosphorylation of TRAIP-T325 (Figure 4B). This suggests that TTF2 bridges TRAIP to Polε in mitotic cells. Since the amino-terminal half of TTF2 is sufficient to support the TRAIP-TTF2-Polε complex and mitotic CMG disassembly (Figure S11A-B), we used AlphaFold to screen for interactions between TTF2 1-504 and the four subunits of Polε. This indicated that a short peptide downstream of the TTF2 Zinc fingers binds to the POLE2 subunit of Polε (Figure 5A, TTF2 124-129; Figure S14 shows that the interaction is predicted with high confidence in all five AlphaFold models). The predicted interaction involves highly conserved residues in TTF2 (Figure 5A-B, TTF2 124-129) and does not clash with binding of POLE2 to the POLE1 catalytic subunit of Polε, or impede association of Polε with the replisome (Figure 5C).

**Figure 5.**
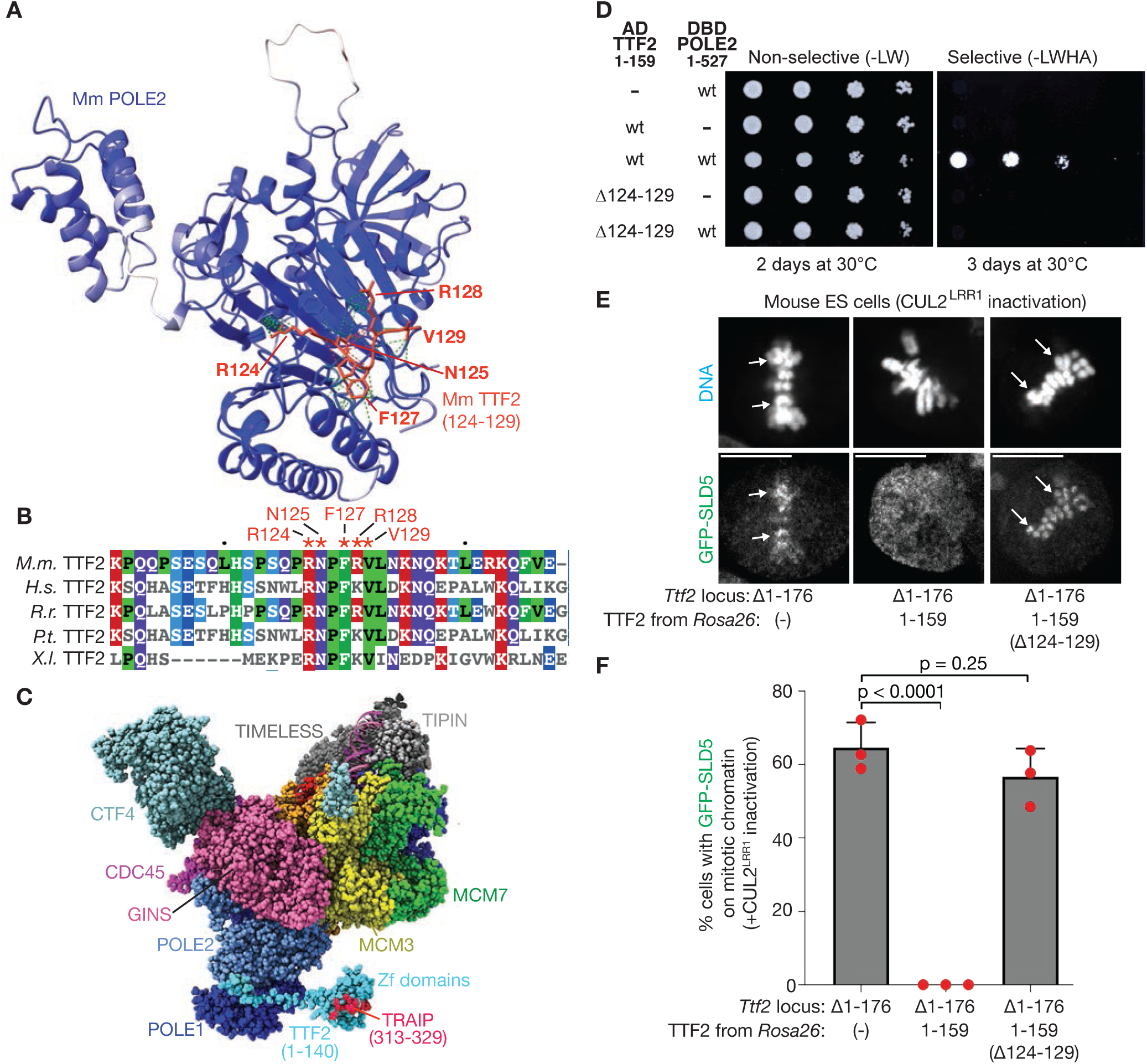
An amino terminal fragment of TTF2, comprising the tandem Zinc fingers and a binding site for POLE2, is sufficient for mitotic CMG disassembly. (A) AlphaFold 3 prediction of the interaction between mouse POLE2 and TTF2 124-129, with contacts depicted as green dashed lines. (B) Multiple sequence alignment of the TTF2 region containing the POLE2-binding motif. M.m. = *Mus musculus*, H.s. = *Homo sapiens*, R.r. = *Rattus rattus*, P.t. = *Pan troglodytes*, X.I. = *Xenopus laevis*. (C) Docking of mouse TTF2 124-140 onto the cryo-electron microscopy structure of the human replisome (PDB 7PLO), together with TTF2 1-123 in complex with TRAIP 313-329 containing phosphorylated TRAIP-T325. Structures of the TTF2 and TRAIP fragments were predicted by AlphaFold Multimer and AlphaFold 3. ‘Zf domains’ = Zinc finger domains. (**D**) Yeast two-hybrid assay with budding yeast cells expressing the indicated versions of TTF2 1-159 fused to the Gal4 activation domain (AD-TTF2 1-159) and POLE2 fused to the Gal4 DNA-binding domain (DBD-POLE2 1-527). (**E**) CUL2^LRR1^ activity was inhibited in cells of the indicated genotypes, before detection of CMG on mitotic chromosomes (white arrows) as in Figure 1D. Scale bar = 10 µm. (F) Quantification of the data from (E), as in Figure 1E.

The association of TTF2 with POLE2 was confirmed via the yeast two-hybrid assay, which provides a convenient tool for the validation of AlphaFold predictions (Xia *et al*, 2023). Thus, a fragment comprising the first 159 amino acids of mouse TTF2 interacted in the two-hybrid assay with mouse POLE2, dependent upon the conserved residues of TTF2 that are predicted to contact POLE2 (Figure 5D). Moreover, TTF2 1-159 was sufficient to support CMG helicase disassembly during mitosis in mouse ES cells (Figure 5E-F, TTF2 1-159), again dependent upon the interaction site with POLE2 (Figure 5E-F, TTF2 1-159 (Δ124-129)). These data indicated that the amino terminus of TTF2 supports mitotic CMG disassembly by bridging phosphorylated TRAIP to the POLE2 subunit of Polε (Figure 5C).

### Phospho-TRAIP, the TTF2 Zinc fingers and Polε drive mitotic ubiquitylation of the MCM3 and MCM7 subunits of CMG

The above data support a model whereby the TRAIP-TTF2-Polε complex positions TRAIP on the replisome during mitosis to induce CMG helicase ubiquitylation. Direct monitoring of mitotic CMG ubiquitylation in cells is challenging, as the vast majority of the genome is normally replicated before mitosis, with the associated replisomes being disassembled during DNA replication termination. Therefore, we took advantage of the fact that Calyculin A drives mouse ES cells prematurely from S-phase into mitosis (Figure S1), to explore the regulation of mitotic CMG ubiquitylation by TRAIP, TTF2 and Polε, under conditions where cells still contain many DNA replication forks. Treatment of asynchronous mouse ES cells for one hour with Calyculin A was sufficient to induce phosphorylation of TRAIP-T325, and complex formation with TTF2 and Polε, analogous to when cells were arrested in mitosis by treatment with STLC (Figure 6A).

**Figure 6.**
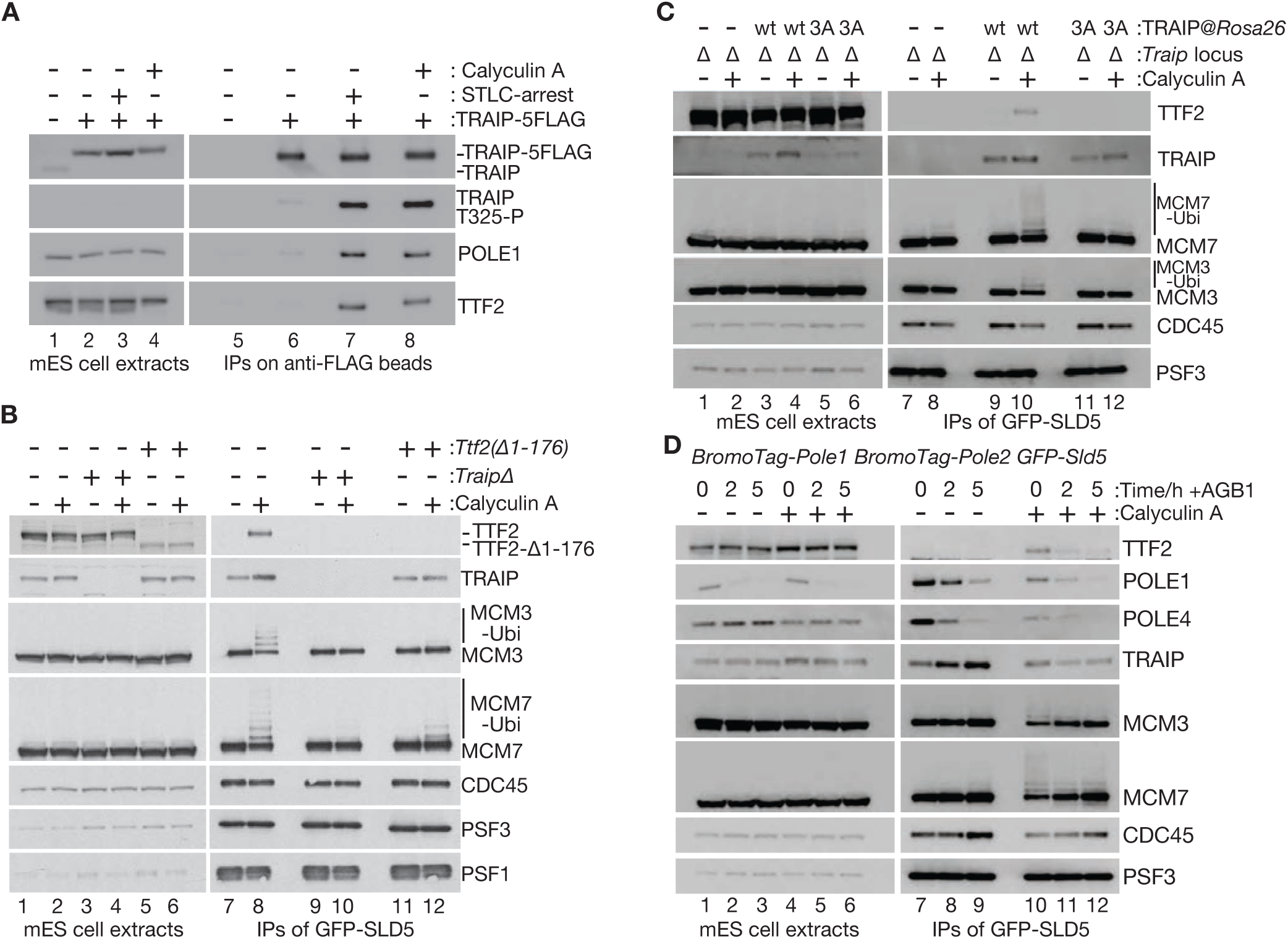
Phospho-TRAIP, the TTF2 Zinc fingers and Polε drive mitotic ubiquitylation of the MCM3 and MCM7 subunits of CMG. (A) Cells of the indicated genotypes were treated as shown with 100 nM Calyculin A or 5 µM STLC. Extracts were incubated with magnetic beads coated with anti-FLAG antibodies, and the indicated factors were monitored by immunoblotting, including phosphorylation of TRAIP-T325 (TRAIP-T325-P). (B) Cells were treated as shown with 100 nM Calyculin A. The CMG helicase was then isolated from cell extracts by immunoprecipitation of GFP-tagged SLD5. The indicated factors were monitored by immunoblotting. (C) Analogous experiment with *TraipΔ* cells expressing the indicated forms of TRAIP from the *Rosa26* locus. (D) Cells with the BromoTag degron fused to both POLE1 and POLE2 (see Figure S16) were treated as shown with 100 nM calyculin A for one hour, before addition of AGB1 for the indicated times. CMG was isolated from cell extracts by immunoprecipitation of GFP-SLD5, before immunoblotting of the indicated factors.

To monitor CMG ubiquitylation when cells entered mitosis prematurely from S-phase, we isolated CMG and associated factors from cell extracts, by immunoprecipitation of the SLD5 subunit of the helicase. In untreated cells, CMG co-purified with TRAIP as observed previously (Villa *et al*., 2021), likely reflecting the S-phase association of TRAIP with the replisome to mediate DNA repair reactions in front of replication forks (Figure 6B, lane 7). After treatment with Calyculin A, TTF2 was recruited to CMG (Figure 6B, lane 8), dependent upon TRAIP (Figure 6B, lane 10). Correspondingly, a ladder of modified bands was detected for the MCM3 and MCM7 components of CMG in the presence of Calyculin A (Figure 6B, lane 8), uniquely amongst the 11 subunits of the helicase (Figure S15A). Modification of MCM3 and MCM7 corresponded to ubiquitylation, since the USP2 deubiquitylase could remove the modified bands (Figure S15B), in a reaction that was sensitive to propargylated ubiquitin that inhibits deubiquitylase activity (Ekkebus *et al*, 2013). Ubiquitylation of MCM3 and MCM7 was dependent upon mitotic kinase activity (Figure S15C) and the TRAIP ubiquitin ligase (Figure 6B, lane 10), but was not affected by inactivation of CUL2^LRR1^ (Figure S15D). CMG ubiquitylation in response to Calyculin A did not cause large-scale helicase disassembly, probably due to most complexes having ubiquitin chains that were too short for efficient recruitment of p97 (Fujisawa *et al*, 2022), reflecting a general reduction in bulk ubiquitylation in cells treated with Calyculin A (Figure S15E).

Mitotic ubiquitylation of the MCM3 and MCM7 subunits of CMG required the amino terminal region of TTF2, which contains the tandem Zinc fingers and the POLE2 binding site (Figure 6B, lane 12), and was also dependent on the CDK consensus sites of TRAIP (Figure 6C, lane 12). To test directly whether Polε was required for mitotic CMG ubiquitylation, we used CRISPR-Cas9 to introduce the BromoTag degron into the *Pole1* and *Pole2* loci in mouse ES cells (Figure S16A-B). Depletion of BromoTag-POLE1 or BromoTag-POLE2 (Figure S16C) was sufficient to block cell proliferation (Figure S16D) but simultaneous depletion of both factors improved the efficiency with which residual Polε and associated factors like CTF18-RFC were displaced from CMG (Figure S16E). When *BromoTag-Pole1 BromoTag-Pole2* cells were treated with AGB1 for 2 hours or 5 hours, with Calyculin A added for the final hour in both cases, displacement of Polε from CMG led to a concomitant displacement of TTF2 and a corresponding reduction in the ubiquitylation of the MCM3 and MCM7 helicase subunits (Figure 6D, lanes 10-12). These findings indicate that Polε is required for mitotic CMG ubiquitylation, together the TTF2 amino terminal region and the TRAIP ubiquitin ligase. Interestingly, recruitment of TTF2 to the replisome in mitotic cells was not only dependent upon Polε, but also upon phosphorylated TRAIP (Figure 6B, compare lanes 8 and 10; Figure 6C, compare lanes 10 and 12). This suggests that TRAIP makes additional contacts with replisome components that contribute to stable engagement of TTF2 with DNA replication forks during mitosis, via the TRAIP-TTF2-Polε complex.

### Both TRAIP and the TTF2 amino terminus are required for MiDAS in mouse ES cells

In response to mitotic protein kinase activation in *Xenopus* egg extracts, TRAIP-dependent ubiquitylation and disassembly of the CMG helicase leads to replication fork cleavage and repair (Deng *et al*., 2019). Similarly, depletion of TRAIP or inhibition of TRAIP ubiquitin ligase activity in human cells inhibits MiDAS (Sonneville *et al*., 2019). Using similar protocols to those developed for human cells, MiDAS can also be detected when mouse ES cells enter mitosis following a period of DNA replication stress (Figure 7A-C, *Control*), corresponding to increased foci of the nucleoside precursor ethynyl-2’-deoxyuridine (EdU). Under such conditions, MiDAS is impaired though not abolished entirely by deletion of TRAIP (Figure 7C, *TraipΔ*). Strikingly, MiDAS is similarly defective in cells that lack the amino terminus of TTF2, containing the Zinc fingers and the POLE2 binding site (Figure 7C, *Ttf2(Δ1-176)*). These data indicate that the ability of TTF2 to form a complex with phosphorylated TRAIP and Polε is important to process sites of incomplete DNA replication when mammalian cells enter mitosis.

**Figure 7.**
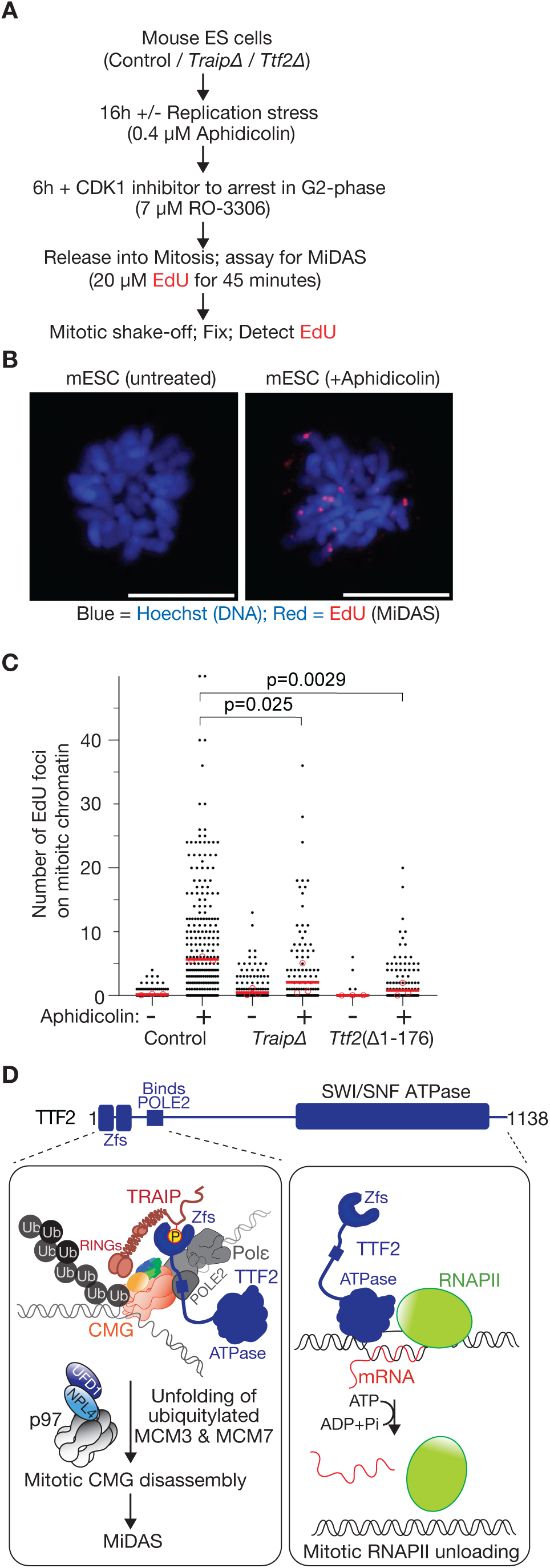
Both TRAIP and the TTF2 amino terminus are required for MiDAS in mouse ES cells. (A) Experimental scheme. MiDAS = Mitotic DNA Synthesis. EdU = ethynyl-2’-deoxyuridine. (B) Example of MiDAS in mouse ES cell. A z-stack of five images with 1 µm spacing was gathered and the images show a 2-dimensional projection of the 3-dimensional data. The scale bars represent 10 µm. (C) Quantification of the number of EdU foci on chromatin per mitotic cell. Individual datapoints from three independent experiments are shown as black spots. The mean values from each experiment are denoted by red circles, with a red line showing the average of the three means. P-values were calculated using a one-way ANOVA followed by Dunnett’s multiple-comparison test. (D) Model for how TTF2 contributes to mitotic CMG disassembly (left panel) and mitotic unloading of RNA polymerase II (RNAPII) from chromosomes (right panel). In the left panel, the three sub-assemblies of the CMG helicase are shown in different colours: the hexameric MCM2-7 ATPases as a ring of red dumbells, the GINS complex in green and blue, and CDC45 in orange. ‘Zfs’ = tandem Zinc finger domains of TTF2. The TTF2 motif that binds to the POLE2 subunit of Polε is shown as a blue rectangle. ‘RINGs’ = two RING domains of the TRAIP dimer, which recruit ubiquitin conjugating enzymes. ‘P’ = phosphorylation of TRAIP-T325.

## Discussion

Work with *Xenopus laevis* egg extracts indicates that TRAIP associates with the leading edge of the vertebrate replisome during S-phase, to mediate DNA repair reactions in front of DNA replication forks, without causing ubiquitylation and disassembly of the CMG helicase *in cis* (Wu *et al*., 2019). At present, a structural understanding for how TRAIP associates with the replisome during S-phase is lacking for any species. TRAIP forms an elongated dimer via its extended region of coiled coil (Figure 2, Figure S3) and it is likely that the amino-terminal RING finger domains of the TRAIP dimer are orientated to the front of the replication fork and away from the replisome itself. The association of TRAIP with the replisome during S-phase does not require TTF2, which itself is only recruited to the replisome during mitosis (Figure 6B, compare lanes 7-8).

In response to the activation of mitotic protein kinases, TRAIP, TTF2 and Polε form a complex (Figure 2), within which the TTF2 amino terminal region plays a central role by bridging phosphorylated TRAIP to Polε (Figure S11A). The tandem Zinc fingers of TTF2 bind to a region of the disordered tail of TRAIP, dependent upon phosphorylation of TRAIP-T325 (Figure 4), and represent a new class of phospho-receptor domain. Simultaneously, a motif downstream of the TTF2 tandem Zinc fingers binds to the POLE2 subunit of Polε (Figure 5). In this way, we propose that TTF2 and Polε reorientate the TRAIP ubiquitin ligase on the replisome during mitosis (Figure 7D, Figure 5C), allowing the RING domains of the TRAIP dimer to direct the ubiquitylation of MCM3 and MCM7 (Figure 6), which are adjacent to each other within the structure of the CMG helicase (Costa *et al*, 2011; Davey *et al*, 2003; Yuan *et al*, 2016). The observed selectivity for the MCM3 and MCM7 subunits of CMG might reflect flexible tethering of TRAIP on TTF2 and Polε, adjacent to MCM3-MCM7, combined with shielding of other CMG subunits that are located underneath other replisome components (Figure 5C). Alternatively, TRAIP might make additional replisome contacts after docking onto TTF2 and Polε, thereby orientating the RING domains more specifically towards MCM3 and MCM7. Distinguising between these possibilities will be aided by biochemical reconstitution of mitotic CMG ubiquitylation and further structural analysis in future studies.

Ubiquitylation of MCM3 and MCM7 would lead to their unfolding by the p97 ATPase (Bodnar & Rapoport, 2017; Deegan *et al*., 2020; Fujisawa *et al*., 2022; Mukherjee & Labib, 2019), in turn disrupting the integrity of the CMG helicase. In this way, CMG disassembly would expose replication fork DNA to nucleolytic attack, leading to the initiation of MiDAS at sites of incomplete DNA replication (Figure 7D), and subsequently facilitating sister chromatid segregation during anaphase.

It is striking that TTF2 was previously found to remove RNA polymerase II from chromosomes during mitosis (Jiang *et al*., 2004), apparently analogous to its role in removing the CMG helicase from mitotic chromosomes. However, our data indicate that the two mechanisms are completely different, with the unloading of RNA polymerase II requiring TTF2 ATPase activity and CMG helicase disassembly involving assembly of the TRAIP-TTF2-Polε complex by the TTF2 Zinc fingers and POLE2-binding motif. It is interesting to consider why TTF2, rather than some other factor, should have been selected during the course of evolution to mediate mitotic CMG disassembly in animal cells. One possibility is that the recruitment of TTF2 to mitotic chromosomes, where it removes RNA polymerase II and nascent transcripts to help shutdown mitotic transcription (Carmo *et al*., 2023; Jiang *et al*., 2004), might help to concentrate the TRAIP-TTF2-Polε complex in the vicinity of mitotic chromosomes, thereby promoting rapid CMG disassembly.

Mammalian TTF2 and *Drosophila* lodestar are both important for the segregation of sister chromatids during mitosis (Girdham & Glover, 1991; Jiang *et al*., 2004). A recent study indicated that the eviction of nascent transcripts from mitotic chromosomes is not the only reason why Lodestar is important for chromosome segregation (Carmo *et al*., 2023). In the future, it will be interesting to explore whether TTF2 displaces other factors from chromatin during mitosis, in addition to RNA polymerase II and the CMG helicase, either via TTF2 ATPase activity or via recognition of additional phospho-proteins by the TTF2 Zinc fingers.

## Materials and Methods

Reagents and resources used in this study are listed in Table S4.

## Plasmid DNA construction

PCRs were performed with Phusion® High-Fidelity DNA Polymerase (New England Biolabs, M0530), and the amplified DNA fragments were cloned with the Gibson Assembly Cloning Kit (New England Biolabs, E2611), according to the manufacturer’s protocol. All new constructs were verified by Sanger sequencing.

Coding sequences of mouse TRAIP (Uniprot identifier Q8VIG6-1) and TTF2 (Uniprot identifier Q5NC05) were amplified from XpressRef Universal Total mouse RNA (QIAGEN, 338112) by RT-PCR (TaKaRa, RR014), and then cloned into the pSC-B-Amp/Kan vector that is part of the StrataClone Blunt PCR Cloning Kit (Stratagene, 240207).

To express recombinant proteins in *E. coli*, genes were cloned into the ‘K27SUMO’ vector (Stein *et al*, 2014), in which 14His-tagged Smt3 (budding yeast SUMO) was fused to the amino terminus of the protein of interest.

To design pairs of guide RNAs (gRNAs) to target a specific site in the mouse genome, ‘The CRISPR Finder’ provided by the Welcome Sanger Institute was used (https://wge.stemcell.sanger.ac.uk//find_crisprs). Annealed oligonucleotides containing the homology region were phosphorylated with T4 polynucleotide kinase (New England Biolabs, M0201) and then ligated into the BbsI site of the vectors pX335 and pKN7 (Table S4) with T4 DNA ligase (New England Biolabs, M0202), as previously described (Pyzocha *et al*, 2014).

Donor vectors for genome editing of the *Pole1, Pole2*, and *Ttf2* loci in mouse ES cells were assembled by golden gate cloning, using the previously reported ‘mammalian toolkit’ (Evrin *et al*., 2023).

To express genes from the *Rosa26* locus, the coding sequence of the gene of interest was cloned into pKN18 (no selective marker) or pKL3223 (Hygromycin selection marker), which drive expression from the CAG or EF1 promoters respectively.

## Culture of mouse ES cells

Mouse ES cells were grown as described previously (Villa *et al*., 2021). E14tg2a cells (Table S4) were cultured in a humidified atmosphere of 5 % CO_2_, 95 % air at 37 °C under feeder-free conditions in serum-containing medium with Leukemia Inhibitory Factor (LIF; MRC PPU Reagents and Services DU1715). All culturing dishes were precoated with PBS containing 0.1 % (w/v) gelatin (Sigma-Aldrich, G1890). The medium was based on Dulbecco’s Modified Eagle Medium (DMEM; ThermoFisher Scientific, 11960044), supplemented with 10 % Fetal Bovine Serum (FBS, FCS-SA/500, Labtech), 5 % Knockout serum replacement (ThermoFisher Scientific, 10828028), 2 mM L-Glutamine (ThermoFisher Scientific, 25030081), 100 U/ml Penicillin-Streptomycin (ThermoFisher Scientific, 15140122), 1 mM Sodium Pyruvate (ThermoFisher Scientific, 11360070), a mixture of seven non-essential amino acids (ThermoFisher Scientific, 11140050), 0.05 mM β-mercaptoethanol (Sigma-Aldrich, M6250) and 0.1 µg / ml LIF. For passaging, cells were released from dishes using 0.05 % Trypsin-EDTA (ThermoFisher Scientific, 25300054).

Mouse ES cells were grown 1.0 x 10^7^ cells were plated on a 15 cm dish and grown for 24 hours at 37 °C. Subsequently, 5 µM STLC (S-Trityl-L-cysteine, Santa Cruz, sc-202799A) was added to the medium before incubation for a further eight hours.

To drive cells into mitosis prematurely, 100 nM Calyculin A (Santa Cruz, sc-202799A) was added for one hour.

To induce ubiquitylation and degradation of proteins fused to degrons, cells were treated with 250 nM AGB1 (TOCRIS, 7686) for BromoTag fusions, or 250 nM dTAG-13 (MRC PPU Reagents and Services) for dTAG fusions.

## Cell survival assay for mouse ES cells

For the experiments in Figure S2 and S4, 0.5 x 10^5^ cells were plated into individual wells of a 24-well plate. After six hours, the medium was changed for fresh medium containing the indicated concentrations of Mitomycin C or Olaparib and incubated for 24 hours. The drug-containing medium was then replaced with fresh medium lacking Mitomycin C or Olaparib, and incubation continued for another four days.

For the experiments with degron-tagged proteins in Figure 3D, Figure S9C and Figure S16D, 500 - 1,000 cells were spread on a 10 cm plate. After six hours, the indicated PROTAC (AGB1 for BromoTag, dTAG-13 for dTAG) was added to the medium at 250 nM and incubation continued for a further 7-10 days.

The surviving cells were washed with PBS and then fixed with methanol, before staining with 0.5 % crystal violet solution (Sigma Aldrich, HT90132). The plates were then imaged with a scanner.

## siRNA transfection of mouse ES cells

Lipofectamine™ RNAiMAX Transfection Reagent (ThermoFisher Scientific, 13778075) was used to transfect siRNA into mouse ES cells, according to the manufacturer’s instructions. *Lrr1* siRNA (Horizon Discovery, J-057816-10) and *Ttf2* siRNA (Horizon Discovery, L-048681-01) were transfected at a final concentration of 50 nM. After 24 hours, the transfection was repeated and incubation continued for a further 24 hours. To inactivate CUL2^LRR1^, cells were subjected to two rounds of transfection with *Lrr1* siRNA as above, and then additionally treated with 5 µM MLN4924 (Activebiochem, A1139) during the final 5 hours (Figure S15) or 12 hours (Figures 1, 3, 4, 5, S4, S11) of transfection, as described previously (Villa *et al*., 2021).

## Transfection of plasmids into mouse ES cells and selection of clones

A mixture of 1 µg of each plasmid DNA (including one plasmid expressing Puromycin resistance), 100 µl of OPTI-MEM (ThermoFisher Scientific, 31985062) and 15 µl of 1 mg/ml PEI (PEI; Polysciences, Inc 24765-2) was vortexed and incubated at room temperature for 15 minutes. Subsequently, 1.0 × 10^6^ cells resuspended in 200 µl OPTI-MEM and 115 µl of DNA-PEI mix were gently mixed and incubated at room temperature for 30 minutes in a 1.5 ml microfuge tube. Cells were then transferred to a single well of a 6-well plate that contained 2 ml of complete DMEM medium. Cells were incubated for 24 hours after transfection, followed by two 24-hour rounds of selection with fresh medium containing 2 µg / ml puromycin (ThermoFisher Scientific, A1113802). The surviving cells were then released from the wells with 0.05 % Trypsin / EDTA, diluted with complete DMEM medium and plated on a 10 cm dish. To select cells expressing the marker resistance genes from donor vectors, 7.5 µg / ml Blastcidin S (ThermoFisher Scientific, A1113902) or 200 µg / ml Hygromycin B (ThermoFisher Scientific, 10687010) was included in the medium during colony formation. Subsequently, colonies were expanded and monitored as appropriate by immunoblotting, PCR and DNA sequencing of the target locus.

## CRISPR-Cas9 mediated genome editing of mouse ES cells

Pairs of guide RNAs (gRNAs) to target specific sites in the mouse genome were selected using ‘The CRISPR Finder’ provided by the Welcome Sanger Institute (https://wge.stemcell.sanger.ac.uk//find_crisprs). Annealed oligonucleotides containing the homology region were phosphorylated with T4 polynucleotide kinase (New England Biolabs, M0201) and then ligated into the BbsI site of the vectors pX335 and pKN7 (Table S4) with T4 DNA ligase (New England Biolabs, M0202), as previously described (Pyzocha *et al*., 2014).

To create a small deletion at a specific locus, mouse ES cells were transfected with two plasmids expressing a corresponding pair of gRNAs together with the Cas9-D10A ‘nickase’ mutant and the puromycin resistance gene. To introduce tags into specific genomic loci, cells were transfected with an appropriate donor vector containing silent mutations to prevent recutting of the edited locus by Cas9-D10A (Table S4), in addition to the plasmids expressing Cas9-D10A, a pair of gRNAs and puromycin resistance. To express derivatives of TRAIP or TTF2 at the *Rosa26* locus, a corresponding donor vector containing silent mutations to prevent recutting of the edited locus by Cas9-D10A (Table S4) was transfected into cells together with pKN88 expressing Cas9-D10A, a single gRNA for the *Rosa26* locus, and puromycin resistance.

## Genotyping of mouse ES cells by PCR and DNA sequencing

Cells from a single well of a 6-well plate were resuspended in 100 µl of 50 mM NaOH and the sample was then heated at 95 °C for 15 minutes. Subsequently, 11 µl of 1 M Tris-HCl (pH 6.8) was added to neutralise the pH. A 0.5 µl aliquot of the resulting genomic DNA solution was used as the template for genotyping purposes in 25 µl PCR reactions with Ex Taq DNA polymerase (TaKaRa, RR001). PCR products were subcloned with the TOPO TA cloning Kit (Invitrogen, K457540) and sequenced with T3 primer (5′-AATTAACCCTCACTAAAGGG-3′).

## Expression of recombinant proteins in bacterial cells

Plasmids expressing 14His-SUMO-tagged proteins were transformed into the *E. coli* strain Rosetta (DE3) pLysS (Novagen, 70956). Transformed cells were inoculated into a 100 ml culture of LB medium supplemented with kanamycin (50 µg / ml) and chloramphenicol (35 µg / ml), before growth overnight at 37°C with shaking at 200 rpm. The following morning, the culture was diluted five-fold with 900 ml of LB medium supplemented with kanamycin (50 µg / ml) and chloramphenicol (35 µg / ml) and then left to grow at 37°C until an OD600 of 0.8 was reached. Protein expression was then induced overnight at 18°C. To express mouse TRAIP, 50 µM IPTG and 100 µM ZnOAc were added to the bacterial culture. For expression of mouse TTF2, 20 µM IPTG was added to the bacterial culture. Subsequently, cells were harvested by centrifugation for 15 minutes in a JLA-9.1000 rotor at 5,180 x g and the pellets were stored at –20 °C.

## Protease inhibitors

One tablet of ‘Roche cOmplete EDTA-free protease inhibitor Cocktail’ (Roche, 11873580001) was either added to 20 ml of protein purification buffer or else dissolved in 1 ml of water to make a 20X stock.

## Buffers

TRAIP-Lysis buffer:

## 50 mM Tris-HCl (pH 7.5), 0.5 M NaCl, 5 mM Mg(OAc)_2_ 10 %(v/v) Glycerol, 0.01 % (v/v) IGEPAL CA-630 and 0.5 mM TCEP

TRAIP-Gel filtration buffer:

## 50 mM BisTris-HCl (pH 6.0), 0.5 M NaCl, 5 mM Mg(OAc)_2_, 0.3 M sorbitol, 0.01 % (v/v) IGEPAL CA-630 and 0.5 mM TCEP

TRAIP-Ion exchange buffer A:

## 50 mM Tris-HCl (pH 7.0), 5 mM Mg(OAc)_2_, 10%(v/v) Glycerol, 0.01 % (v/v) IGEPAL CA-630 and 0.5 mM TCEP

TRAIP-Ion exchange buffer B:

## 50 mM BisTris-HCl (pH 6.0), 1 M NaCl, 5 mM Mg(OAc)_2_, 10 % (v/v) Glycerol, 0.0 1 %(v/v) IGEPAL CA-630 and 0.5 mM TCEP

Lysis buffer:

## 50 mM Tris-HCl (pH 8.0), 0.5 M NaCl, 5 mM Mg(OAc)_2_ and 0.5 mM TCEP

Gel filtration buffer:

## 20 mM Hepes-KOH (pH 7.8), 0.15 M NaCl, 5 mM Mg(OAc)_2_, 0.3 M sorbitol and 0.5 mM TCEP

Ion exchange buffer A:

50 mM Tris-HCl (pH 8.0), 5 mM Mg(OAc)_2_, 10 % (v/v) Glycerol and 0.5 mM TCEP. <u>Ion exchange buffer B:</u> 50 mM Tris-HCl (pH 8.0), 1 M NaCl, 5 mM Mg(OAc)_2_ 10 % (v/v) Glycerol and 0.5 mM TCEP.

EMSA buffer:

## 25 mM Hepes-KOH (pH 7.9), 100 mM KOAc, 10 mM Mg(OAc)_2_, 0.02 % (v/v) IGEPAL CA-630, 0.1 mg/ml BSA, 10 % Glycerol, 1 mM DTT

Wash buffer:

## 100 mM Hepes-KOH (pH 7.9), 100 mM KOAc, 10 mM Mg(OAc)_2_, 2 mM EDTA, 0.1 % (v/v) IGEPAL CA-630

Mouse ES cell lysis buffer:

100 mM Hepes-KOH (pH 7.9), 100 mM KOAc, 10 mM Mg(OAc)_2_, 2 mM EDTA, 10 % (v/v) Glycerol, 0.1 % (v/v) Triton X-100. <u>High pH SDS buffer:</u> 31.5 mM Tris-HCl (pH 6.8), 10 % (v/v) glycerol, 1 % (w/v) SDS, 0.005 % (w/v) Bromophenol Blue, 150 mM Trizma base

## Purification of mouse TRAIP

Bacterial pellets expressing 14His-SUMO-TRAIP (∼20 g from 4 L cultures) were resuspended in 40 ml TRAIP-Lysis buffer supplemented with 30 mM imidazole, 500 U Pierce™ Universal Nuclease and 1X Roche cOmplete EDTA-free protease inhibitor Cocktail. Samples were lysed by incubation with 1 mg / ml lysozyme on ice for 30 minutes, followed by sonication for 90 seconds (15 seconds on, 30 seconds off) at 40% on a Branson Digital Sonifier, then clarified by centrifugation at 10,000 x g for 30 minutes in an SS-34 rotor (Sorvall).

The supernatant was mixed with 1 ml resin volume of Ni-NTA beads (Qiagen, 30210) which had been equilibrated with TRAIP-Lysis buffer supplemented with 30 mM imidazole. After 1 hour incubation at 4°C, beads were recovered in a disposable gravity flow column and washed with 150 ml of TRAIP-Lysis buffer containing 30 mM imidazole. 14His-SUMO-TRAIP was eluted with 5 ml of Lysis buffer containing 400 mM imidazole. Yeast Ulp1 SUMO protease (10 µg / ml) was added to the eluate and incubated at 4°C for 2 hours, to cleave 14His-Smt3 from TRAIP. The sample was then loaded onto a 1 ml HiTrapQ column that had been equilibrated with a mixture of 85% TRAIP-Ion exchange buffer A and 15% TRAIP-Ion exchange buffer B. Flow-through fractions were collected, diluted with 1 volume of TRAIP-Ion exchange buffer A and loaded onto a 1 ml HiTrap Heparin that had been equilibrated with 85% TRAIP-Ion exchange buffer A and 15% TRAIP-Ion exchange buffer B. The bound proteins were eluted by a 25 ml gradient of TRAIP-Ion exchange buffer A with 25-60% TRAIP-Ion exchange buffer B. TRAIP-containing fractions were pooled and diluted with 0.5 volume of TRAIP-Ion exchange buffer A, loaded onto a 1 ml HiTrap Heparin that had been equilibrated in 75% TRAIP-Ion exchange buffer A and 25% TRAIP-Ion exchange buffer B. The bound proteins were eluted by an 8 ml gradient of TRAIP-Ion exchange buffer A with 25-60% TRAIP-Ion exchange buffer B. Fractions containing TRAIP were pooled, concentrated with an Amicon Ultra Centrifugal Filter Unit (Merck, UFC803024), loaded onto a 24 ml Superose 6 column equilibrated in TRAIP-Gel filtration buffer. TRAIP eluted from the column around ∼13 ml, similar to the ∼670 kDa thyroglobulin marker protein (Figure 2D). Purified TRAIP was concentrated with an Amicon Ultra Centrifugal Filter Unit, aliquoted and snap-frozen with liquid nitrogen.

## Purification of mouse TTF2

Bacterial pellets from cultures expressing 14His-SUMO-TTF2 (1 L for wt TTF2 TTF2-Δ1-100 and TTF2 1-100, 2 L for TTF2-K578A) were resuspended in 10 ml (20 ml for TTF2-K578A) of Lysis buffer containing 30 mM imidazole and 1X Roche cOmplete EDTA-free protease inhibitor Cocktail. The samples were lysed by incubation with 1 mg / ml lysozyme on ice for 30 minutes, followed by sonication for 90 seconds (15 seconds on, 30 seconds off) at 40% on a Branson Digital Sonifier, before clarification by centrifugation at 10,000 x g for 30 minutes in an SS-34 rotor.

The supernatant was mixed with 1 ml resin volume of Ni-NTA beads that had been equilibrated with Lysis buffer containing 30 mM imidazole. After 1 hour at 4°C, beads were recovered in a disposable gravity flow column and washed with 150 ml of Lysis buffer containing 30 mM imidazole. Recombinant TTF2 proteins were then eluted with 5 ml of Lysis buffer containing 400 mM imidazole. For wt TTF2, TTF2-Δ1-100 and TTF2-K578A, samples were diluted with 1 volume of Ion exchange buffer A and then loaded onto a 1 ml HiTrapQ column that had been equilibrated in Ion exchange buffer A with 25% Ion exchange buffer B. Flow-through fractions were then collected, loaded onto a 1 ml HiTrap Heparin that had been equilibrated in Ion exchange buffer A with 25% Ion exchange buffer B. The bound proteins were eluted by a 10 ml gradient of Ion exchange buffer A with 25-100% Ion exchange buffer B. Fractions containing TTF2 were pooled and concentrated with an Amicon Ultra-4 Centrifugal Filter Unit (Merck, UFC803024). This ion exchange chromatography step was not required for TTF2 1-100. Finally, all TTF2 samples were loaded onto a 24 ml Superdex 200 column that had been equilibrated in Gel filtration buffer. Purified TTF2 eluted around ∼10 ml and was concentrated with an Amicon Ultra-4 Centrifugal Filter Unit, aliquoted and snap-frozen with liquid nitrogen.

## Mass photometry analysis of purified TRAIP

The mass of mouse TRAIP was measured by with a mass photometer (Refeyn OneMP, Refeyn). Mass photometry buffer (25 mM Hepes-KOH pH 7.9, 150 mM NaCl, 10 mM MgOAc and 1 mM DTT) was freshly prepared and filtered with a 0.22 µm filter before use. Samples were loaded into a silicon gasket well (Grace Bio-Labs reusable CultureWell™ gaskets, Merck, GBL103250) on a coverslip (Precision cover glasses thickness No. 1.5H, Marienfeld, 0107222) mounted onto an oil-immersion lens. Protein standards (NativeMark™ Unstained Protein Standard, Invitrogen, LC0725) were used for calibration and to produce a standard curve. A 12 μl aliquot of diluted TRAIP (6.4 ng / µl (∼50 nM of TRAIP dimer)) was loaded into the chamber and movies of 60 s duration were recorded. Data acquisition was performed using Acquire MP (Refeyn Ltd, v1.1.3) and analysed using Discover MP software.

## *In vitro* ATPase assay

For the experiment in Figure S10, recombinant mouse TTF2 (60 nM) was incubated at room temperature in 50 μl reaction buffer, comprising 50 mM Tris-HCl pH7.5, 50 mM NaCl, 1 mM MgCl_2_, 1 mM ATP (Pi-free) and 3 µM ‘Phosphate sensor’ (Thermo Scientific #PV4407). The latter is a recombinant *E. coli* phosphate-binding protein, labelled with the fluorophore MDCC (7-Diethylamino-3-[N-(2-maleimidoethyl)carbamoyl]coumarin), which exhibits fluorescence in the presence of free phosphate generated by ATP hydrolysis. Pi-free ATP was prepared by incubating 10 mM ATP (Thermo Scientific #1030430), 50 mM NaCl, 150 mM Tris pH7.5, 1 mM MgCl2, 20 μM 7-Methylguanosine (Sigma #M0627) and 4 U / mL Nucleoside Phosphorylase (Sigma #N8264) for 30 minutes at 30°C.

Phosphate release derived from ATP hydrolysis was measured as fluorescence intensity, using a Varian Cary Eclipse spectrometer (Gemini Lab), with excitation and emission wavelengths of 430 and 460 nm respectively. During the experiment, plasmid DNA (pRF096, 8.2 kb) was added at the indicated times to a final concentration of 20 ng / µl.

## Monitoring the DNA binding activity of TTF2 by electrophoretic mobility shift assay

A single-stranded DNA substrate was prepared by synthesising and PAGE purifying an 80-nucleotide oligo (YX011), labelled at the 5’ end with the far-red fluorescent dye Cyanine 5 (5’-Cy5-CAGTGCCAAGCTTGCATGCCTGCAGGTCGACTCTAGAGGATCCCCGGGTACCGAGCTCGAATTCGTAATCATGGTCATAG-3’).

To prepare a double-stranded DNA substrate, the complementary 80-nucleotide sequence to the above (YX012) was synthesised and PAGE purified (5’-CTATGACCATGATTACGAATTCGAGCTCGGTACCCGGGGATCCTCTAGAGTCGACCTGCAGGCATGCAAGCTTGGCACTG-3’), and the two complementary oligos were then annealed.

For the experiment in Figure S12, 10 nM DNA substrates were mixed with the indicated proteins in 10 µl of EMSA buffer at room temperature for 10 minutes. Reaction were then loaded in ‘Novex™ TBE Gels 4-12% 12 well’ (ThermoFisher, EC62352BOX) and separated at 150 V for 40 minutes. The Cy5-labeled DNA fragments were detected with a GelDoc imaging system (BIO-RAD).

## Isothermal Titration Calorimetry assay

Peptides corresponding to mouse TRAIP 313-329 (FGDEIDLNTTFDVN**<u>T</u>**PPTQ, where **<u>T</u>** corresponds to TRAIP-T325) were synthesised (Peptide & Elephants) with TRAIP-T325 either phosphorylated or unphosphorylated. Subsequently, 1 mg of each lyophilised peptide was dissolved in 120 µl of 200 mM Tris-HCl pH 7.5, corresponding to 3.7 mM of the phosphorylated peptide and 3.9 mM of the non-phosphorylated peptide, before storage at -70°C.

HisSUMO-TTF2 1-100 was dialysed with ITC buffer (50 mM Tris-HCl pH 7.5, 100 mM NaCl, 0.5 mM TCEP) for 16 hours, using a dialysis bag Slide-A-Lyzer MINI Dialysis unit (Thermo Scientific, 69553). Before each measurement, the peptides were further diluted to 200 µM with ITC buffer. All samples were then degassed at room temperature by centrifugation at 21,000 g for 10 minutes.

The isothermal titration calorimetry assay was then performed with a Micro-Cal PEAQ-ITC (Malvern, UK). In each case, 300 µl of 20 µM HisSUMO-TTF2 1-100 was loaded into the cell and 2 µl of 200 µM TRAIP peptides were injected at 19 consecutive timepoints with 130 second intervals. The data were analysed using MicroCal PEAQ-ITC analysis software (Malvern). Titration curves were fitted with baseline subtraction by the software.

## SDS-PAGE and Immunoblotting

Protein samples were resolved by SDS–polyacrylamide gel electrophoresis in NuPAGE Novex 4 - 12% Bis-Tris gels (ThermoFisher Scientific, NP0321 and WG1402A) with NuPAGE MOPS SDS buffer (ThermoFisher Scientific, NP0001), or in NuPAGE Novex 3 - 8% Tris-Acetate gels (ThermoFisher Scientific, EA0375BOX and WG1602BOX) using NuPAGE Tris-Acetate SDS buffer (ThermoFisher Scientific, LA0041). Resolved proteins were either stained with InstantBlue (Expedeon, ISB1L) or were transferred to a nitrocellulose membrane with the iBlot2 Dry Transfer System (Invitrogen, IB21001S). Antibodies used for protein detection in this study are described in Table S4. Conjugates to horseradish peroxidase of anti-sheep IgG from donkey (Sigma-Aldrich, A3415), or anti-mouse IgG from goat (Sigma-Aldrich, A4416) were used as secondary antibodies before the detection of chemiluminescent signals on Hyperfilm ECL (cytiva, 28906837, 28906839) or the AZURE 300 imaging system (Azure Biosystems), using ECL Western Blotting Detection Reagent (Cytiva, RPN2124).

## Phos-tag SDS-PAGE

TRAIP phosphorylation was monitored in 8% polyacrylamide gels containing 100 µM ZnCl_2_ and 50 µM Phos-tag (Kinoshita *et al*., 2006). The separation gel (10 ml in total) comprised 3.6 ml of MilliQ, 3.5 ml of 1 M Bis-Tris-HCl (pH 6.8), 2.67 ml of 30% Acrylamide-Bisacrylamide solution (37.5:1), 100 µl of 5 mM Phos-tag acrylamide (MRC PPU Reagents and Services), 1 µl of freshly prepared 1 M ZnCl_2_, 100 µl of 10 %(w/v) Ammonium persulfate and 25 µl of Tetramethyl ethylenediamine. Stacking gel (5 ml in total) contained 3.65 ml of MilliQ dH_2_0, 0.62 ml of 1 M Bis-Tris-HCl (pH 6.8), 0.65 ml of 2.67 ml of 30% Acrylamide-Bisacrylamide solution (37.5:1), 25 µl of 10 %(w/v) Ammonium persulfate and 5 µl of Tetramethyl ethylenediamine. Proteins were separated at 100 V for 4 hours at 4°C with NuPAGE MOPS SDS buffer (ThermoFisher Scientific, NP0001). Subsequently, Zinc ions were removed by washing the gel three times for 10 minutes each with 10 mM Tris-HCl (pH7.5), 10 mM EDTA and 0.05% SDS, followed by two washes of 5 minutes with MilliQ dH_2_0. Proteins were transferred to a nitrocellulose membrane with the iBlot2 Dry Transfer System (Invitrogen, IB21001S), before analysis by immunoblotting.

## Preparation of antibody-coated magnetic beads

Firstly, 300 mg of Dynabeads M-270 Epoxy (ThermoFisher Scientific, 14302D) were resuspended in 10 ml of dimethylformamide. Subsequently, 425 µl of the activated slurry of magnetic beads, corresponding to ∼ 1.4 x 10^9^ beads, were washed twice with 1 ml of 1 M NaPO_3_ (pH 7.4). The beads were then incubated with 300 µl of 3 M (NH4)_2_SO_4_, 300 µg of anti-FLAG M2 antibody (Sigma-Aldrich, F3165), and 1 M NaPO_3_ (pH 7.4) up to a total volume of 900 µl. Subsequently, the mixture was incubated at 4 °C for 2 days with rotation. Finally, the beads were treated as follows: four washes with 1 ml PBS, 10 minutes in 1 ml PBS containing 0.5 %(w/v) IGEPAL CA-630 with rotation at room temperature, five minutes in 1 ml PBS containing 5 mg / ml BSA with rotation at room temperature and one wash with 1 ml PBS containing 5 mg / ml BSA. The beads were then resuspended in 900 µl PBS containing 5 mg / ml BSA and stored at 4°C.

## Preparation of extracts of mouse ES cells and immunoprecipitation

Protein expression was monitored using denatured cell extracts that were prepared in the presence of trichloro acetic acid (TCA). Firstly, 1.0-3.0 x 10^6^ cells were released from the plate with 0.05 % Trypsin / EDTA as above, then harvested by centrifugation at 350 x g for 3 minutes. The cell pellet was then resuspended in 500 µl of 20 % (v/v) trichloro acetic acid (Sigma-Aldrich T0699), vortexed vigorously and then centrifuged at 500 x g for 10 minutes. The supernatant was discarded, and the pellets were then resuspended in 100 µl of ‘High pH SDS buffer’ before heating at 95 °C for 10 minutes.

To prepare native extracts for immunoprecipitation experiments, cells from two 15 cm dishes cells were released by incubation for 10 minutes with 10 ml of PBS containing 1 mM EGTA and 1 mM EDTA, before harvesting and centrifugation at 350 x g for 3 minutes. The cell pellets (∼0.3 g) were then snap-frozen in liquid nitrogen and stored at -70°C. Subsequently, the cell pellets were thawed and resuspended with an equal volume of ‘Mouse ES cell lysis buffer’, supplemented with 2 mM sodium fluoride, 2 mM sodium β-glycerophosphate pentahydrate, 10 mM sodium pyrophosphate, 1 mM sodium orthovanadate, 1 µg / ml LR-microcystin, 1 mM DTT, 1X Roche cOmplete EDTA-free protease inhibitor Cocktail and 5 μM Propargylated-Ubiquitin. Genomic DNA and RNA were digested for 30 minutes at 4 °C with 1600 U / ml of Pierce Universal Nuclease (ThermoFisher Scientific, 88702). The extracts were then centrifuged at 20,000 x g for 30 minutes at 4°C. A 10 μl aliquot of the supernatant was added to 90 μl of 1.5X LDS sample loading buffer (ThermoFisher Scientific, NP0007), which was heated at 95°C for 3 minutes and then stored the extract sample for the experiment. The rest of the extract was then incubated for 90 minutes as appropriate with either a 15 μl slurry of ChromoTek GFP-Trap magnet Particles M-270 (Proteintech, gtd-100), or a 15 μl slurry of M-270 epoxy beads (ThermoFisher Scientific, 14302D) coupled with anti-FLAG M2 antibody (Sigma Aldrich, F3165). The beads were then washed four times with 1 ml of Wash buffer and the bound proteins eluted at 95 °C for 3 minutes in 40 µl 1X LDS sample loading buffer.

For mass spectrometry analysis, the scale of cell culture was increased to produce a cell pellet of ∼2.0 g, which was then processed for immunoprecipitation as above, except that 150 μl slurry of M-270 epoxy beads coupled to anti-FLAG M2 antibody were used to immunoprecipitate TRAIP-5FLAG. The beads were resuspended after washing in 35 µl 1X LDS sample loading buffer, of which 5 µl was used for immunoblotting. The remaining 30 µl was resolved in a NuPAGE Novex 4-12 % Bis-Tris gel (ThermoFisher Scientific, NP0321 and WG1402A), before staining with SimplyBlue SafeStain (ThermoFisher Scientific, LC6060). Each sample lane was sliced into 40 bands before trypsin digestion and processing for mass spectrometry with Thermofisher Orbitrap platforms (MS Bioworks, https://www.msbioworks.com/). The data were analysed using Scaffold 5 software (Proteome Software Inc, USA). Mass spectrometry data have been deposited at the ProteomeExchange Consortium via the PRIDE partner repository database and will be made publicly available by the date of publication.

## Glycerol gradient ultracentrifugation

Glycerol gradients were manually assembled in an ultra-centrifuge tube (Beckman, P200915MGSG), by placing 45 µl of 30% glycerol buffer (25 mM Hepes-KOH pH 7.9, 100 mM KOAc, 0.02%(v/v), IGEPAL CA-630, 1 mM DTT and 10 mM Mg(OAc)_2_) at the bottom of the tube, followed by consecutive 45 µl aliquots of equivalent buffer containing 25%, 20%, 15% and 10% glycerol. Tubes were left on ice for 30 minutes, after which 5 µl of cell extract or purified protein was placed on the top of gradient. Samples were then ultracentrifuged for two hours at 249,000 g in a Beckman TLS55 rotor at 4°C. Finally, 18 fractions of 14 µl each were retrieved manually from top to the bottom of the gradient, before mixing of each fraction with 5 µl 4X LDS buffer and heating at 95°C for 10 minutes. Samples were then analysed by SDS-PAGE followed either by staining with Coomassie Brilliant Blue, or by immunoblotting.

## Imaging of mouse ES cells by spinning disk confocal microscopy

Cells were grown on chambered 4-well slides (µ-slide 4 well, 80426, Ibidi) and then fixed in 4 % (w/v) paraformaldehyde (ThermoFisher Scientific, J19943.K2) in PBS for 10 minutes at room temperature. The fixed cells were washed with PBS supplemented with 3 % (w/v) BSA, 1 mM MgCl_2_ and 1 mM CaCl_2_ for 30 minutes.

To stain nuclear DNA, cells were incubated at 4°C overnight in a mixture of 1µg / ml Hoechst 333423 (Sigma, B2261), 3 %(w/v) BSA, 1 mM MgCl_2_ and 1 mM CaCl_2_ in PBS. The chamber slides were then washed three times with PBS containing 3 % BSA, 1 mM MgCl_2_ and 1 mM CaCl_2_.

For immunofluorescence analysis of chromatin-bound RNA polymerase II, fixed and washed cells were first permeabilised by detergent extraction, by incubating for 10 minutes in PBS supplemented with 3 % (w/v) BSA, 1 mM MgCl_2_, 1 mM CaCl_2_ and 0.5 % (v/v) Triton X-100. Cells were then incubated for 30 minutes with a blocking mixture comprising PBS supplemented with 3 % (w/v) BSA, 1 mM MgCl_2_ and 1 mM CaCl_2_. Subsequently, cells were incubated at room temperature for 1 hour in a 1:1000 dilution of an antibody specific for phosphorylated Serine 5 in the C-terminal repeats of RPB1 (Abcam, ab5408), in PBS supplemented with 3 % (w/v) BSA, 1 mM MgCl_2_ and 1 mM CaCl_2_. The samples were then washed three times with 3 % (w/v) BSA, 1 mM MgCl_2_ and 1 mM CaCl_2_, before incubation in the dark for one hour at room temperature with a 1:2,000 dilution of secondary antibody (anti-mouse IgG coupled with Alexa Fluor^TM^ 594, Invitrogen, A-21203), in PBS supplemented with 3 % (w/v) BSA, 1 mM MgCl_2_, 1 mM CaCl_2_ and 1 µg / ml Hoechst 33342. After washing three times with PBS containing 1 mM MgCl_2_ and 1 mM CaCl_2_, cells on the chamber slides were mounted with 700 µl of PBS containing 1 mM MgCl_2_ and 1 mM CaCl_2_.

For time-lapse imaging of GFP-SLD5 and TTF2-mCherry in live mouse ES cells in Figure S5B, 2.0 x 10^5^ cells were seeded onto a 35 mm round dish (µ-dish 35 mm high, 81156, Ibidi) on the day before imaging, together with 2 ml of ‘no phenol red DMEM’ (ThermoFisher Scientific, 21063029) that was supplemented as described above (see ‘Culture of mouse ES cells’).

Confocal images of fixed cells were acquired with a Zeiss Cell Observer SD microscope with a Yokogawa CSU-X1 spinning disk, using a HAMAMATSU C13440 camera with a PECON incubator, a 60X 1.4-NA Plan-Apochromat oil immersion objective, and appropriate excitation and emission filter sets. Images were acquired using the ‘ZEN blue’ software (Zeiss) and processed with ImageJ software (National Institutes of Health) as previously described (Sonneville *et al*., 2019).

To monitor the association of CMG or RNA polymerase II with mitotic chromosomes, around 50-150 metaphase cells were monitored in each of three independent experiments. Similarly, MiDAS was assayed by examination of around 50-150 prometaphase cells in three independent experiments.

## Detection of Mitotic DNA Synthesis (MiDAS)

Mouse ES cells were treated with 0.4 μM aphidicolin for 16 hours. During the last 6 hours of aphidicolin treatment, 7 µM RO-3306 (CDK1 inhibitor) was added to block cells in G2-phase. Cells were then rinsed with pre-warmed PBS three times (within 5 minutes) and incubated with fresh medium containing 20 µM EdU and 200 ng / ml nocodazole for 45 minutes. The following steps were performed at room temperature. Mitotic cells were harvested by mitotic-shake off, pelleted by centrifugation at 233 g for 3 minutes, resuspended in 0.5 ml of medium and re-seeded onto microscope slides (Avantor, 631-1553), which was coated by Poly-L-Lysine (Sigma, P8920), heated at 300°C for 30 seconds and dried up before use.

Three minutes after re-seeding, excess medium was removed from the slides with a piece of tissue paper. The attached cells on the slide were then fixed with 100 µl of 4% PFA / PBS for 10 minutes. Subsequently, the slides were washed with PBS with 3% (w/v) BSA for 30 minutes and cells were then permeabilised with PBS containing 0.5% (v/v) Triton X-100 for 20 minutes, before washing twice with PBS with 3% (w/v) BSA.

The incorporation of EdU was detected with the Click-iT® Plus EdU Imaging Kit (Invitrogen, C10640). Firstly, 100 µl of ‘Click-iT® Plus reaction cocktail’ was placed on the slide, before covering loosely with a coverslip and incubating for one hour in the dark. The slides were then placed in a 50 ml tube and treated three times for 20 minutes each with 30 ml of PBS containing 3% (w/v) BSA and 0.5% (v/v) Triton X-100. Finally, the slides were stored at 4°C in PBS containing 3% (w/v) BSA, 0.5% (v/v) Triton X-100 and 1 µg / ml Hoechst.

For imaging, a coverslip was added to the stained cells on the slide and sealed with nail varnish. Confocal images of Hoechst and EdU-Alexa Fluor647 were acquired by spinning disk confocal microscopy as above. A z-stack of images with a 1 µm interval was acquired and ImageJ software was used to generate a 2-dimensional projection of the 3-dimensional data. The number of EdU foci on mitotic chromatin was counted manually.

## Yeast Two-Hybrid Assays

The budding yeast strain PJ69-4A (Table S4) was transformed with derivatives of pGADT7 (protein of interest fused to Gal4 activation domain; *LEU2* marker gene) and pGBKT7 (protein of interest fused to Gal4 DNA binding domain; *TRP1* marker gene). Five independent transformed colonies were resuspended together in water, diluted to 3.33 x 10^6^ cells / ml, and then further diluted to 3.33 x 10^5^, 3.33 x 10^4^, and 3.33 x 10^3^ cells / ml. For each dilution, 15 µl was plated in a series of spots that corresponded to 50,000, 5000, 500 and 50 cells. The plates contained 1.7%(w/v) agar, 2% (w/v) Glucose, ‘Complete supplement drop-out -Ade-His-Leu-Met-Trp-Ura’ (FORMEDIUM, DCS1511) and BD Difco™ Yeast Nitrogen Base without Amino Acids and Ammonium Sulfate (BD, 233520), supplemented as necessary with 0.04 mg / ml adenine and 0.04 mg / ml histidine. The plates were then incubated at 30°C for 2-3 days as indicated and imaged with a scanner.

## Protein Sequence alignments

Protein sequences were aligned using Clustal Omega (https://www.ebi.ac.uk/jdispatcher/msa/clustalo) and visualised with Mview (https://www.ebi.ac.uk/jdispatcher/msa/mview).

## Structure predictions with AlphaFold Multimer and AlphaFold 3

Structural predictions were performed using AlphaFold-Multimer (Evans *et al*., 2022) and AlphaFold 3 (Abramson *et al*., 2024), using ColabFold (Mirdita *et al*., 2022) and the AlphaFold 3 server (https://golgi.sandbox.google.com/about), before analysis with Chimera X (https://www.rbvi.ucsf.edu/chimerax/).

Predicted Alignment Error (PAE) plots of the AlphaFold 3 prediction data were generated using the AlphaFold 3 Analysis Tool (https://predictomes.org/tools/af3/).

## Acknowledgements

We thank Axel Knebel for propargylated ubiquitin, Emma Brannigan & Ron Hay for assistance with assistance with mass photometry, Anna Perez & Yogesh Kulathu for help with isothermal titration calorimetry, Matt Toman, Nicola Wiechens, Ramasubramanian Sundaramoorthy and Tom Owen-Hughes for assistance with ATPase assays and MRC PPU Reagents and Services (https://mrcppureagents.dundee.ac.uk) for antibody production.

## Author contributions

Conceptualization: KL, RF Methodology: RF, KL Investigation: RF Visualization: RF, KL Funding acquisition: KL Project administration: KL Supervision: KL Writing: KL, RF

## Data and materials availability

Materials generated in this study are listed in Appendix Table S4 and are available from MRC PPU Reagents and Services (https://mrcppureagents.dundee.ac.uk) or upon request.

## List of Supplementary Materials

Figs. S1 to S16

Tables S1 to S4.

## Legends to Supplementary Figures

**Figure S1.**
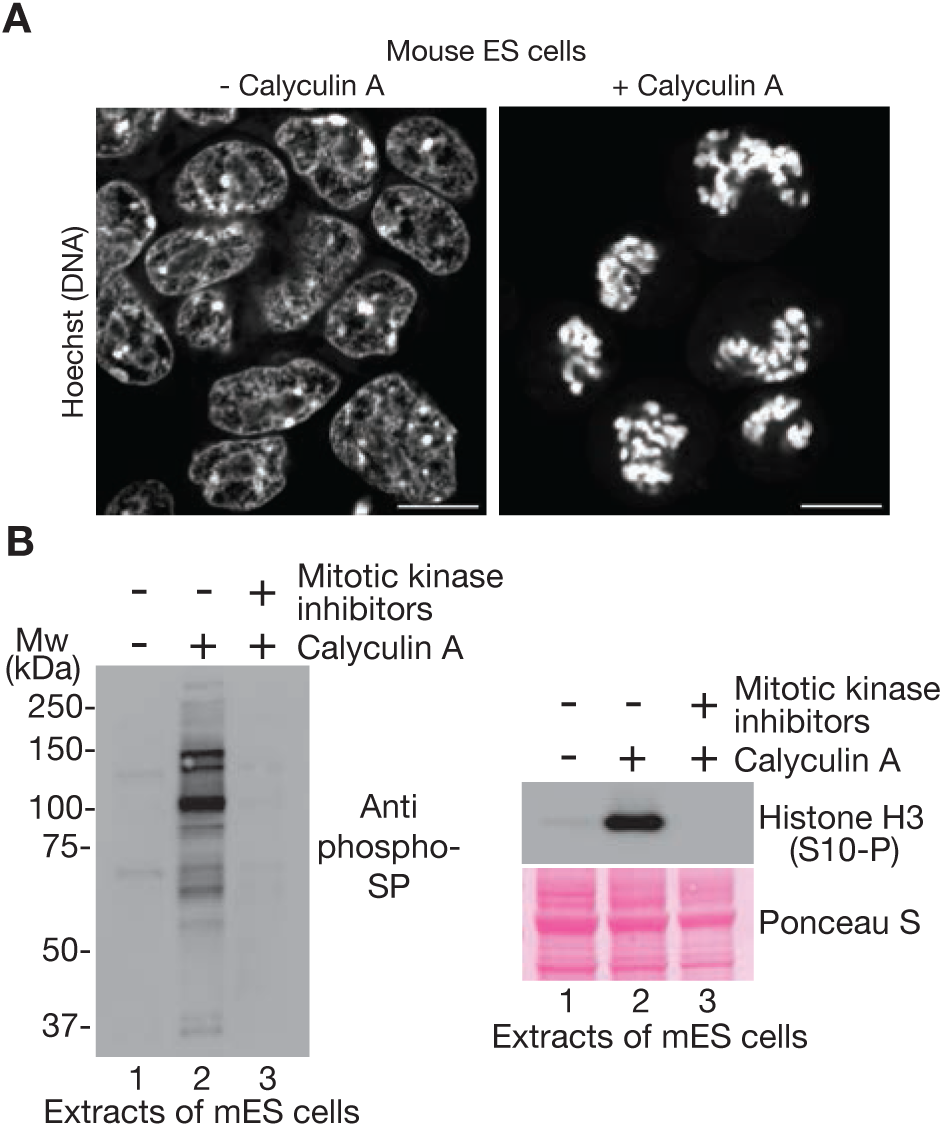
Calyculin A drives mouse ES cells into premature mitosis. **(A)** Spinning-disk confocal microscopy of asynchronous mouse ES cells, before (left) and after (right) treatment with 100 nM Calyculin A for one hour. Scale bars = 10 µm. (**B**) Mouse ES cells were treated as indicated with 100 nM Calyculin A and a cocktail of inhibitors of mitotic protein kinases (10 µM each of the CDK inhibitor RO-3306, the pan-Aurora inhibitor VX-680 and the PLK1 inhibitor BI2536). Global CDK phosphorylation was monitored by immunoblotting with an antibody specific to phosphoSerine-Proline (left panel, anti phospho-SP), whereas Aurora kinase activity was monitored by blotting with an antibody specific to phosphorylated Serine 10 of Histone H3 (right panel, Histone H3 (S10-P)).

**Figure S2.**
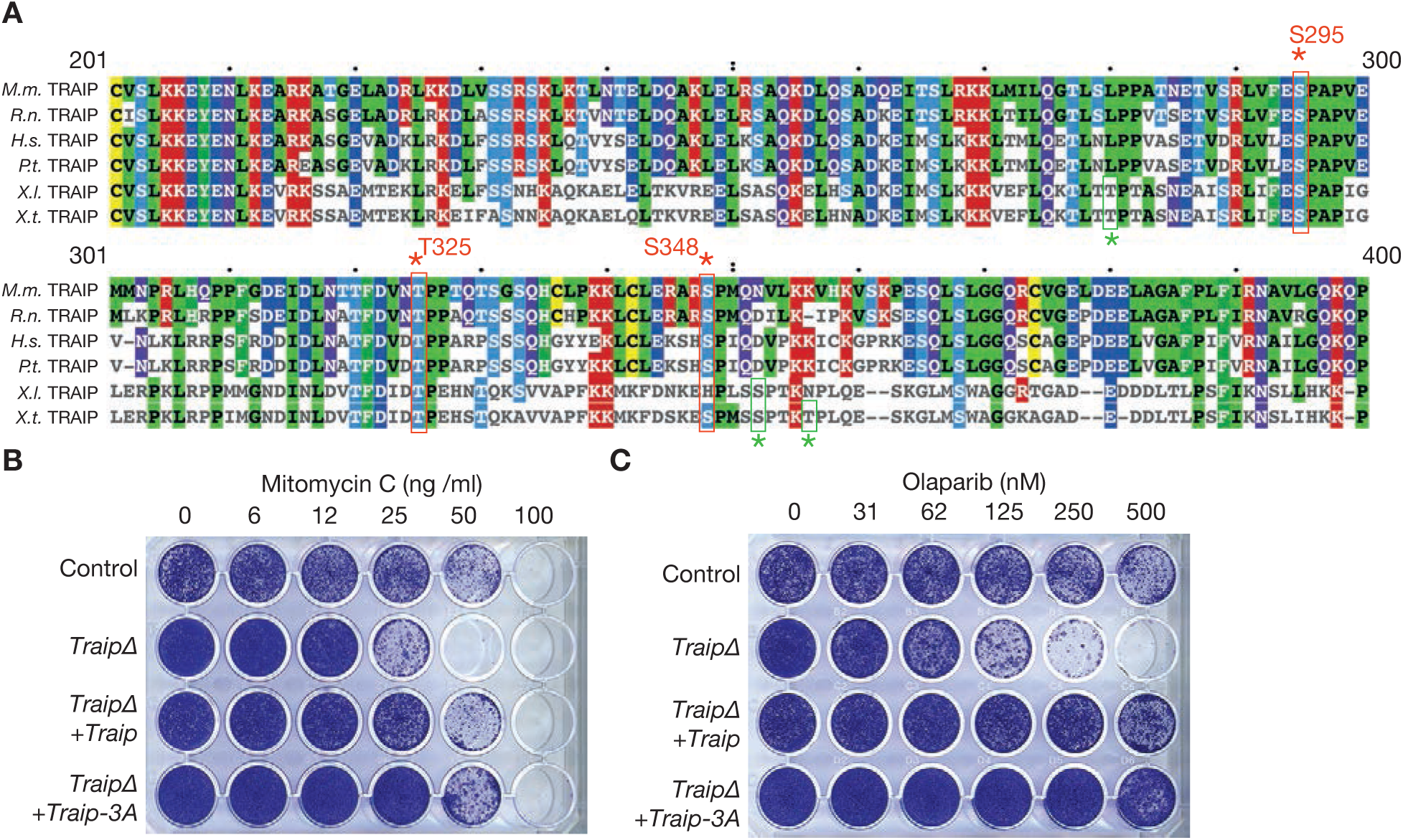
Consensus CDK phosphorylation sites in TRAIP are dispensable for growth in the presence of inter-strand DNA crosslinks or upon inhibition of PARP activity. **(A)** Multiple sequence alignment of the TRAIP region that spans the three consensus CDK sites. *M.m.* = *Mus musculus*, *H.s.* = *Homo sapiens*, *R.r.* = *Rattus rattus*, *P.t.* = *Pan troglodytes*, *X.I.* = *Xenopus laevis*, *X.t.* = *Xenopus tropicalis*. (**B**) Cells of the indicated genotypes were grown in the presence of varying concentrations of Mitomycin C or Olaparib for 24 hours. Subsequently, cells were grown in fresh medium another 4 days before fixation and staining with crystal violet.

**Figure S3.**
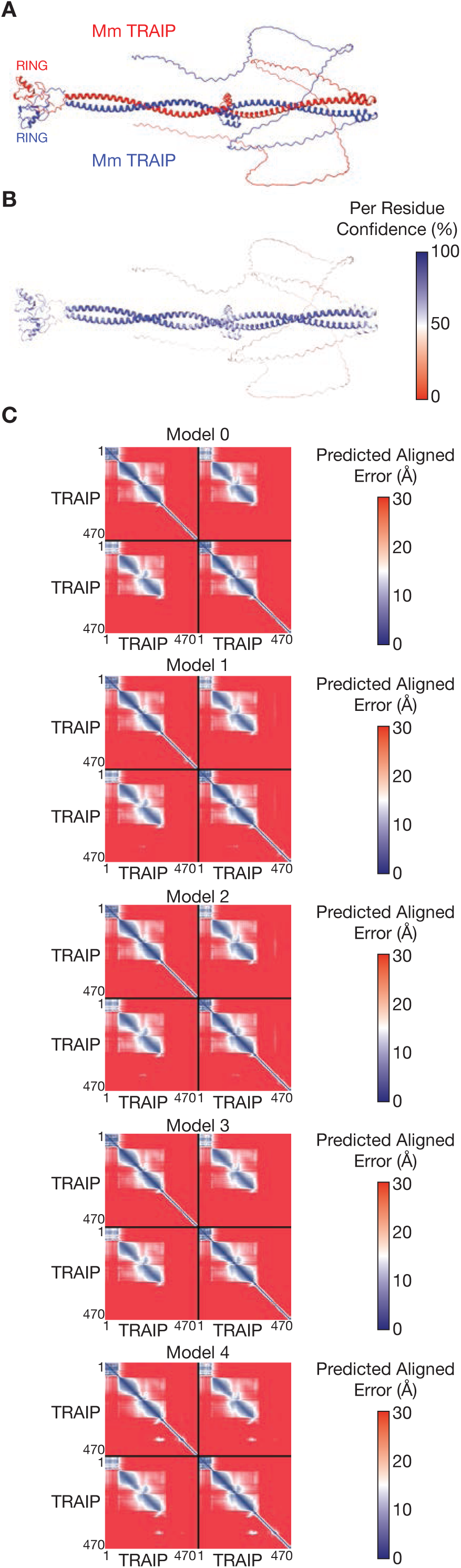
AlphaFold predicts that the TRAIP ubiquitin ligase is an elongated dimer. **(A)** AlphaFold 3 prediction of dimeric mouse TRAIP, showing the location of the RING domains at the amino termini of the two protomers (coloured red and blue). **(B)** The predicted model of TRAIP dimer from (A) is shown with colouring according to Per Residue Confidence. (**C**) Predicted Alignment Error (PAE) plots for the five predictions of the TRAIP dimer by Alphafold3.

**Figure S4.**
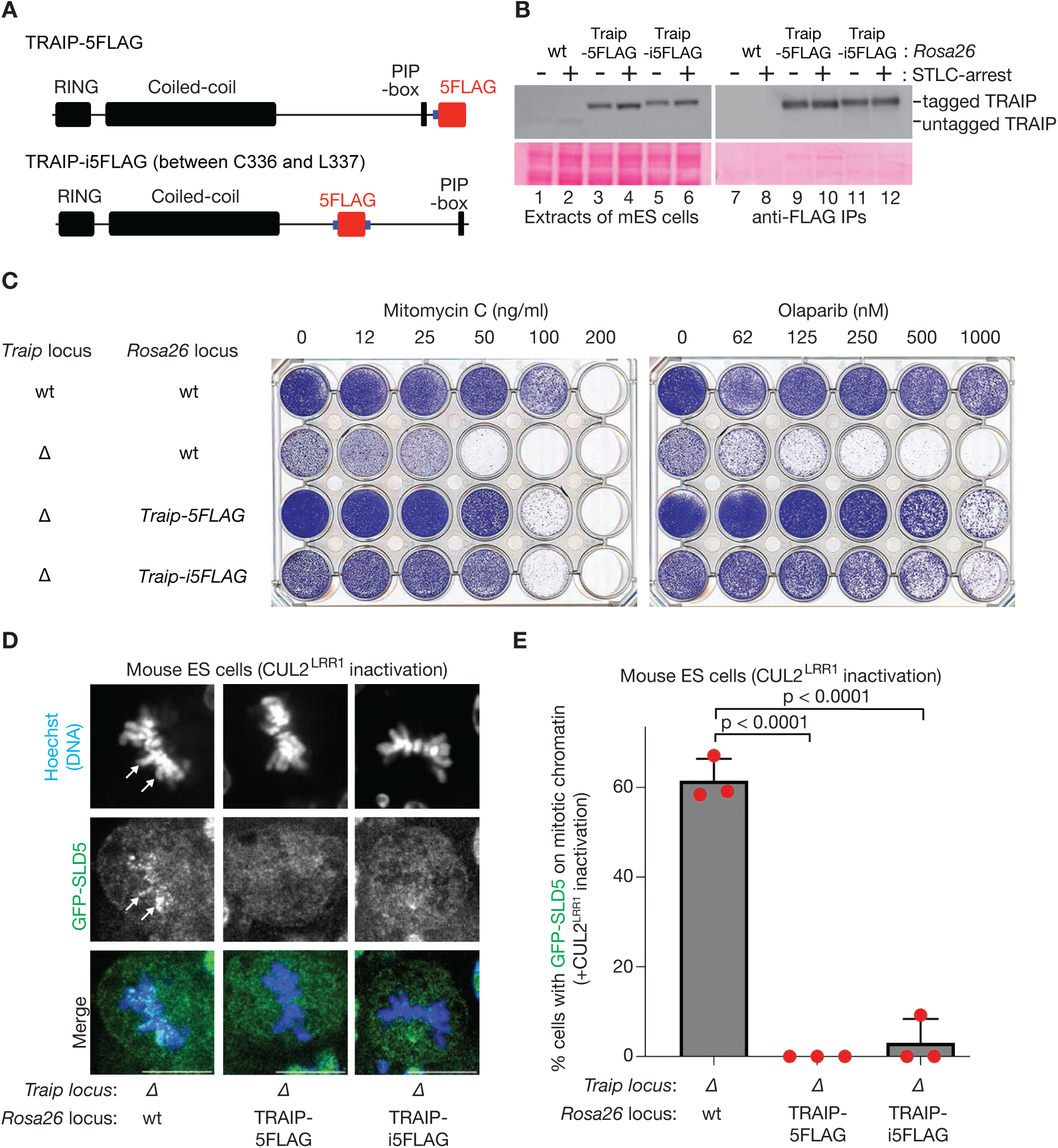
Tagged versions of TRAIP are functional in mouse ES cells. **(A)** Schematic representations of TRAIP protein with the 5FLAG tag (red) inserted at the carboxyl terminus (TRAIP-5FLAG), or internally between Cysteine 336 and Leucine 337 (TRAIP-i5FLAG). A 10-amino-acid linker (5X Gly-Ala, shown in blue) was included between TRAIP and 5FLAG (TRAIP-5FLAG) or on both sides of the 5FLAG-tag (TRAIP-i5FLAG). (**B**) Cells of the indicated genotype were arrested as shown with 5 µM STLC. Cell extracts were incubated with anti-FLAG antibodies before immunoblotting with antibodies specific for TRAIP (upper panels). Immunoblot membranes were stained with Ponceau S (lower panels) to monitor protein loading. (**C**) Cells of the indicated genotype were treated as shown with Mitomycin C or Olaparib for 24 hours. Cells were then grown in fresh medium for another 4 days, before fixation and staining with crystal violet. (**D**) Mitotic CMG disassembly was monitored as in Figure 1D. White arrows indicate the persistence of GFP-SLD5 on mitotic chromatin. Scale bars correspond to 10 µm. (**E**) Quantification of the data from (D). The mean values for three independent experiments are shown as red spots, whereas the histograms show the average of the mean values, with associated standard deviations. P-values were calculated using a one-way ANOVA followed by Dunnett’s multiple-comparison test.

**Figure S5.**
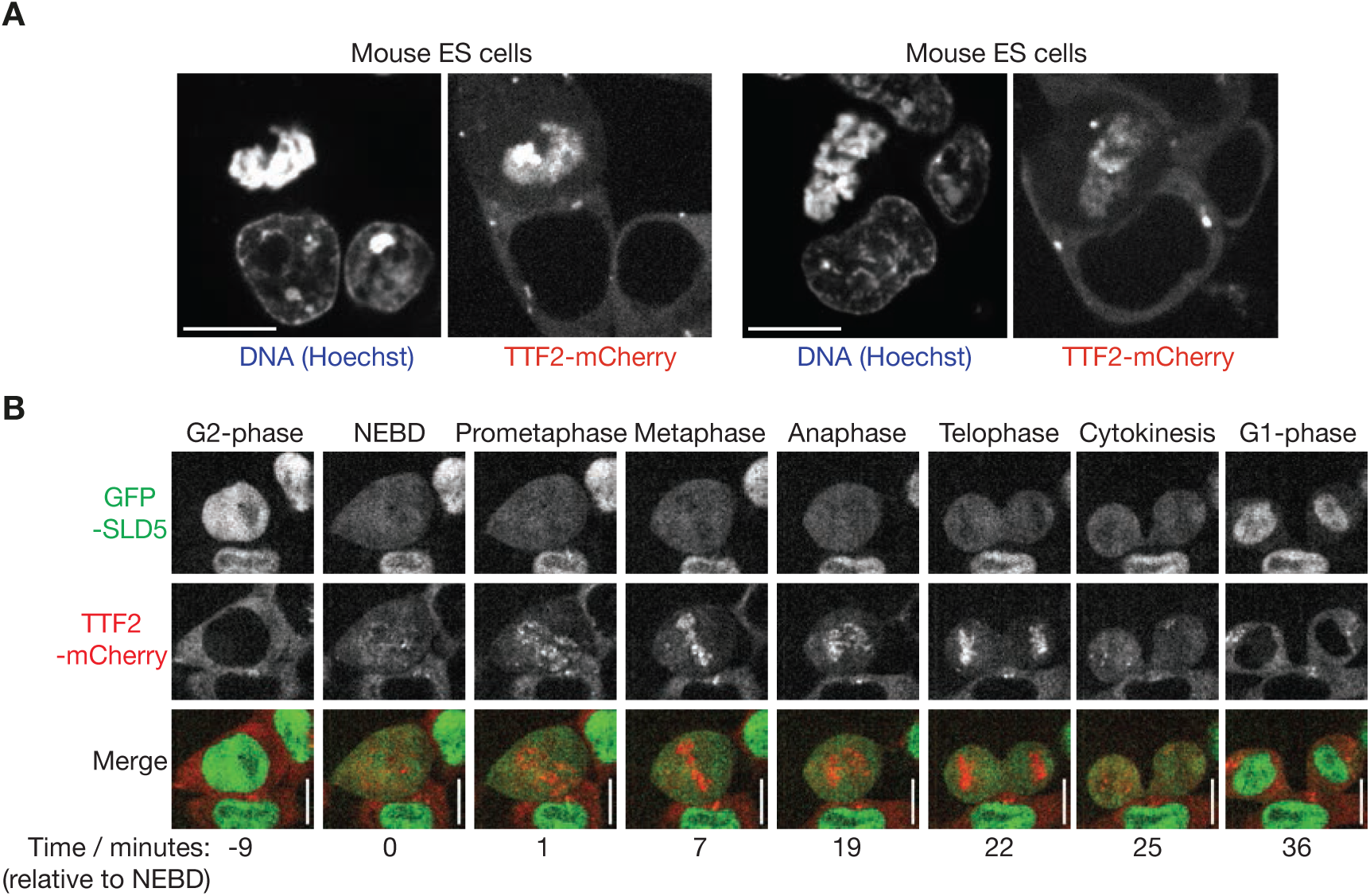
Mouse TTF2 is a cytoplasmic protein that associates with chromatin during mitosis. **(A)** Spinning disk confocal microscopy of live mouse cells expressing TTF2-mCherry from the *Ttf2* locus. (**B**) Time-lapse video microscopy of mouse ES cells expressing GFP-SLD5 and TTF2-mCherry. Cells were imaged at the indicated times, beginning nine minutes before nuclear envelope breakdown (NEBD). Scale bars correspond to 10 µm.

**Figure S6.**
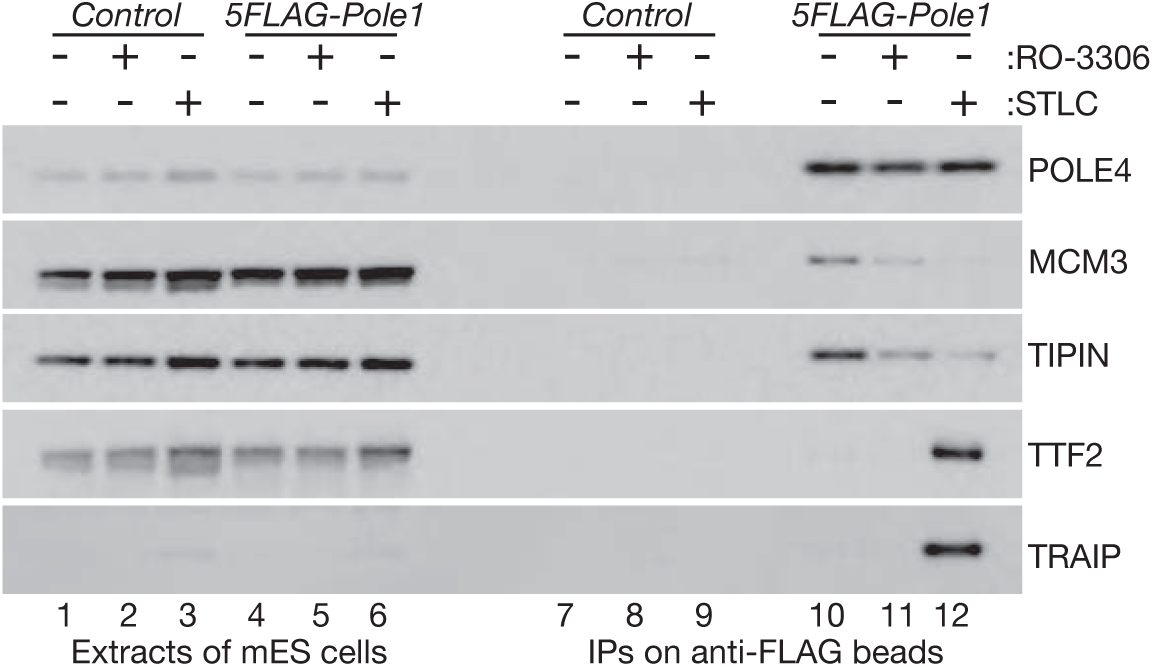
DNA polymerase epsilon associates with TTF2 and TRAIP during mitosis. Control or *5FLAG-Pole1* mouse ES cells were treated as shown with 5 µM STLC for 8 hours, or 7 µM RO-3306 (CDK1 inhibitor) for 6 hours. Extracts were then incubated with anti-FLAG beads, before immunoblotting of the indicated proteins.

**Figure S7.**
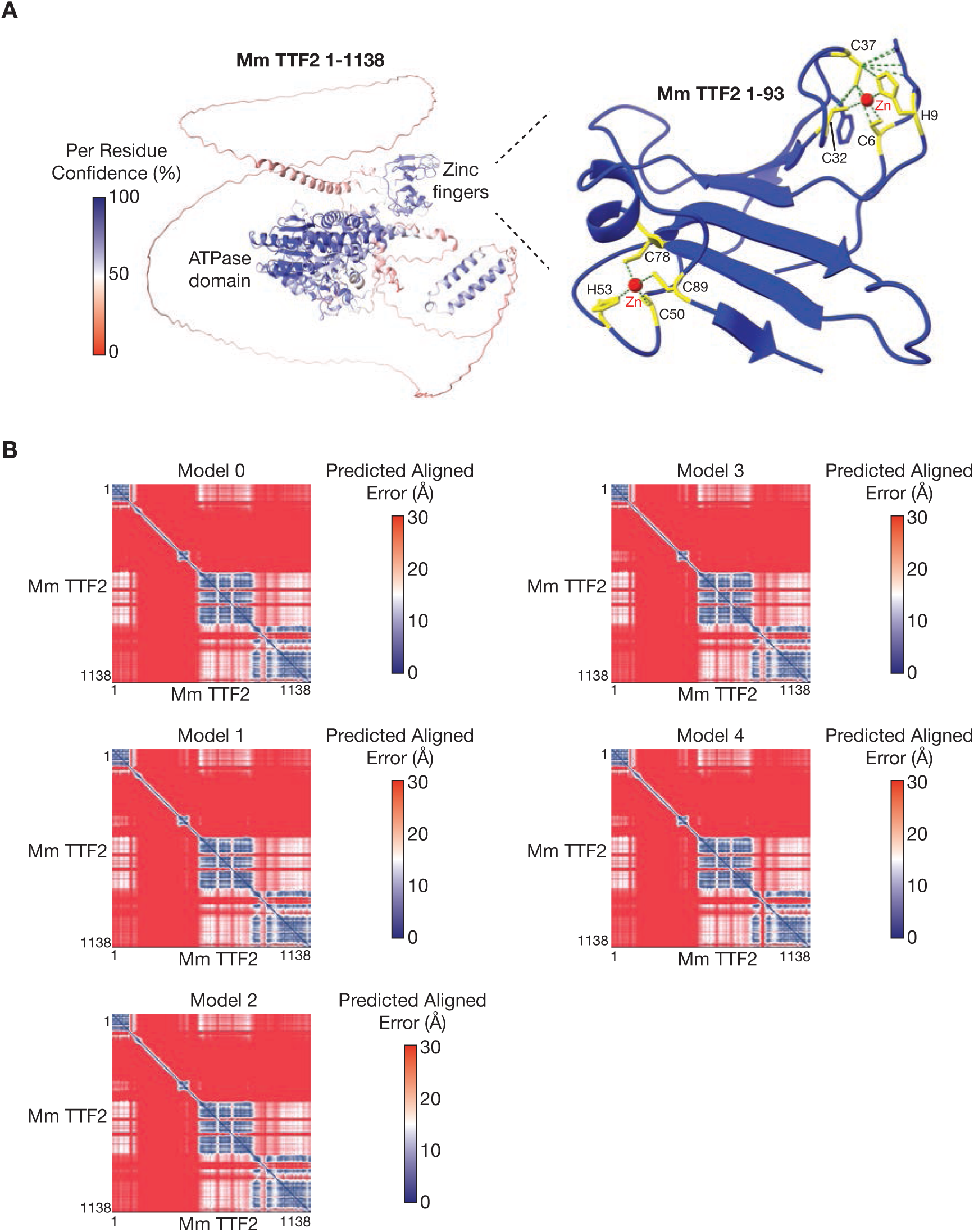
The TTF2 ATPase has a pair of Zinc finger domains at the amino terminus. **(A)** AlphaFold 3 prediction of mouse TTF2 (left panel), showing the location of the pair of Zinc fingers at the amino terminus of the protein, separated by a long-disordered region from the ATPase domain. The prediction is coloured by ‘Per Residue Confidence’. The right panel shows the pair of Zinc finger domains in more detail (‘Zn’ = Zinc ion). (**B**) Predicted Alignment Error (PAE) plots for the five predictions of the TTF2 structure by Alphafold3.

**Figure S8.**
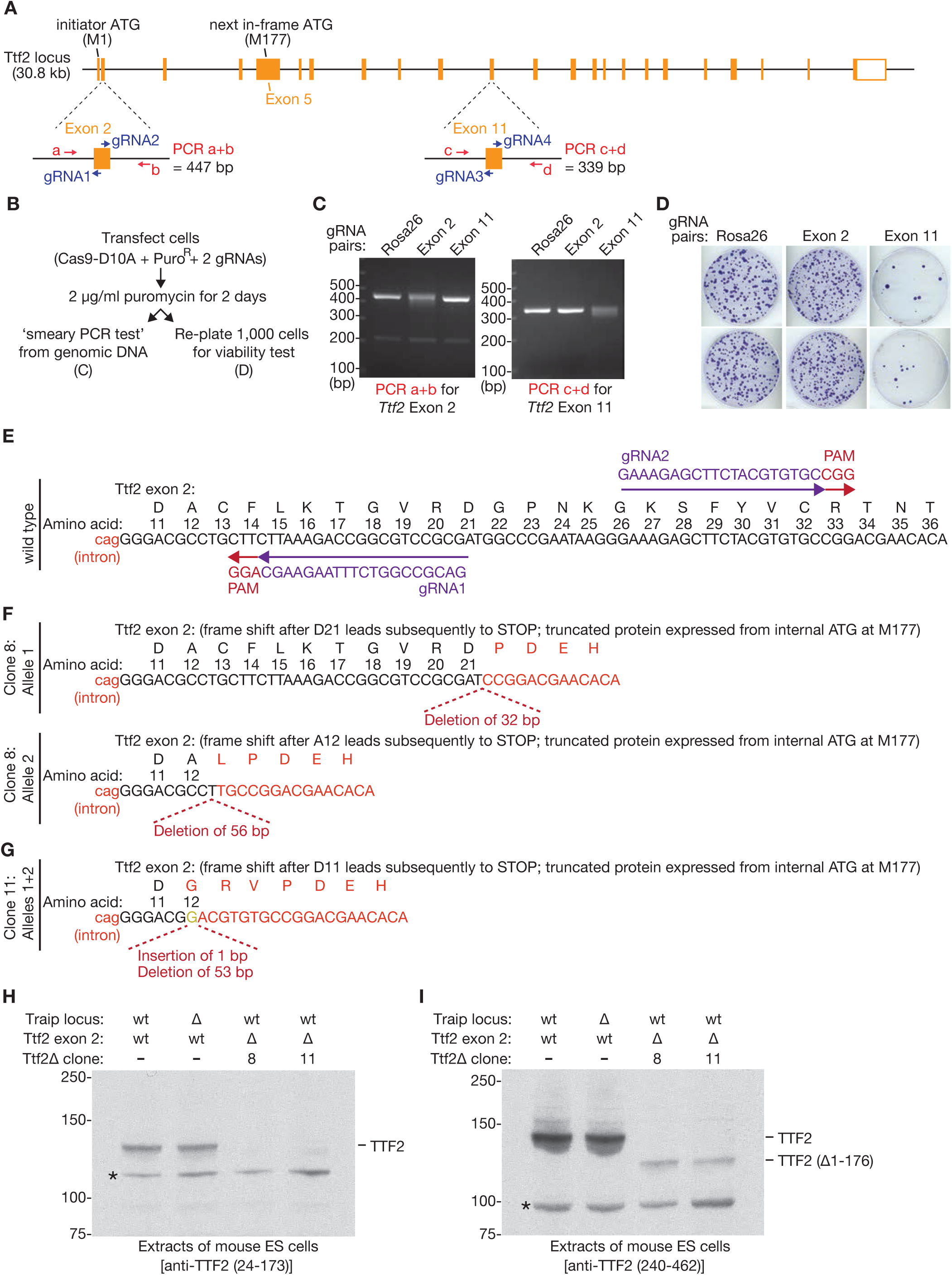
CRISPR deletions leading to expression of truncated TTF2 without the amino terminal Zn fingers are viable in mouse ES cells. **(A)** Outline of the mouse *Ttf2* gene, illustrating the location of gRNA pairs that were used to target the Cas9-D10A nickase to exons 2 and 11. PCR pairs are shown that were used to monitor repair of nicks by Cas9-D10A in *Ttf2* Exon2 (a+b) or *Ttf2* Exon 11 (c+d). (**B**) Summary of experimental procedure. (**C**) PCR tests after targeting Cas9-D10A via pairs of gRNAs to Exon 2 or Exon 11 of *Ttf2*. Nicking and repair leads to a smeary PCR product. (**D**) Cells were processed as in (B) and then grown for 10 days. Targeting Cas9-D10A to Exon 11 reduced viability dramatically, whereas colony formation was not impaired by targeting of Cas9-D10A to Exon **(A)** 2. (**E**) DNA sequence of the region in Exon 2 of *Ttf2* that is targeted by gRNAs 1-2. PAM = Protospacer Adjacent Motif (required for cutting by Cas9). (**F**-**G**) DNA sequence of the equivalent region of Exon 2 of *Ttf2*, for two representative clones from (D) after targeting of Cas9-D10A and subsequent repair. (**H**-**I**) Immunoblotting of cell extracts from *Ttf2* Exon 2Δ clones from (F-G), confirming expression of a truncated form of TTF2 that lacks the amino terminus and is predicted to commence from Methionine 177 of the full-length TTF2 protein.

**Figure S9.**
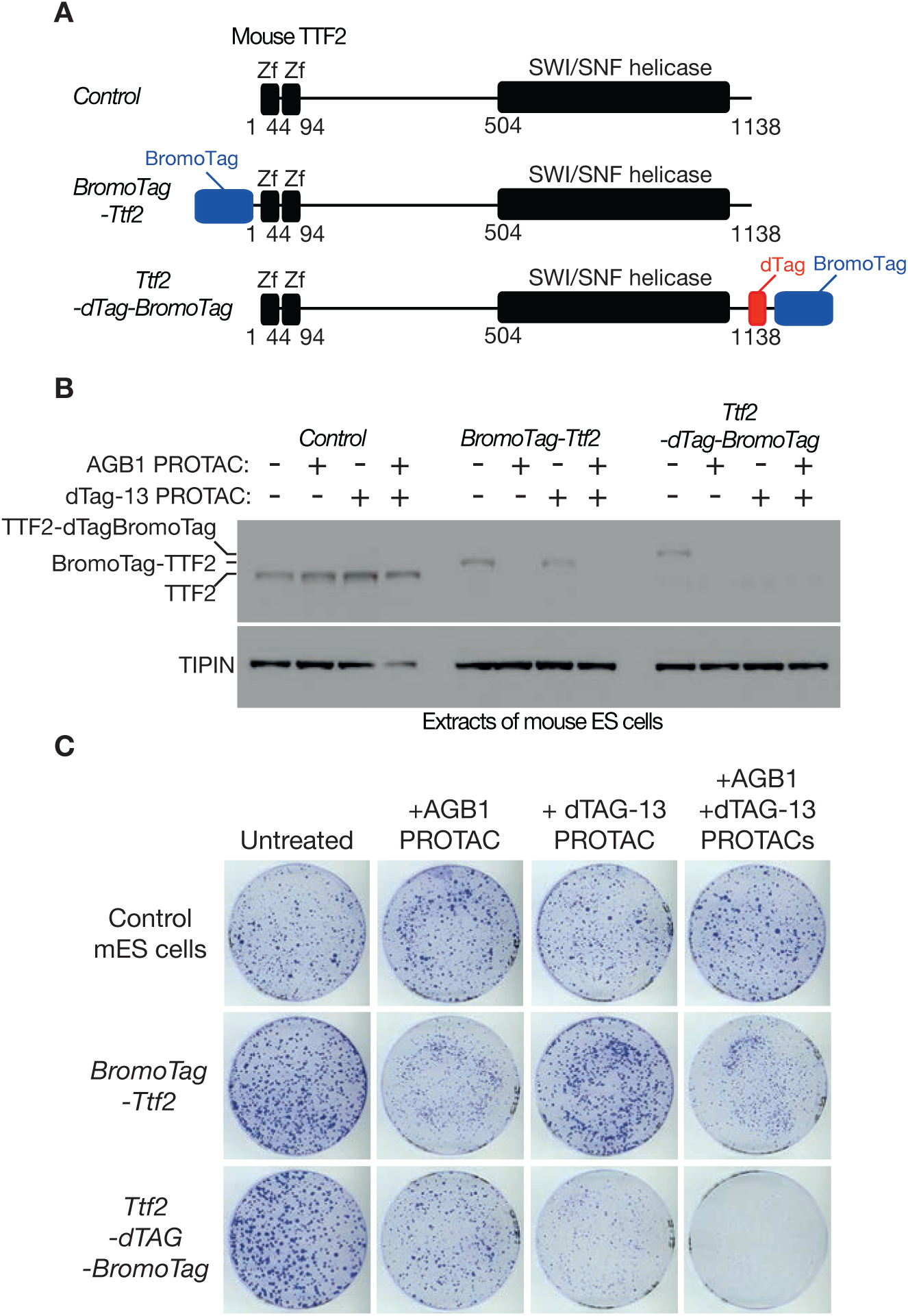
Efficient depletion of degron-tagged TTF2 is lethal in mouse ES cells. **(A)** BromoTag and dTAG degron cassettes were added to TTF2 as shown, at the endogenous *Ttf2* locus in mouse ES cells. ‘Zf’ = Zinc finger domain. (**B**) Cells of the indicated genotype were treated as shown with AGB1 and dTag-13 PROTACs for two hours. The indicated factors were monitored by SDS-PAGE and immunoblotting of cell extracts. (**C**) Approximately 1,000 cells for each of the indicated genotypes were plated and grown on 10 cm plates for eight days, in the presence or absence of PROTACs as shown. Surviving colonies were fixed and stained with crystal violet.

**Figure S10.**
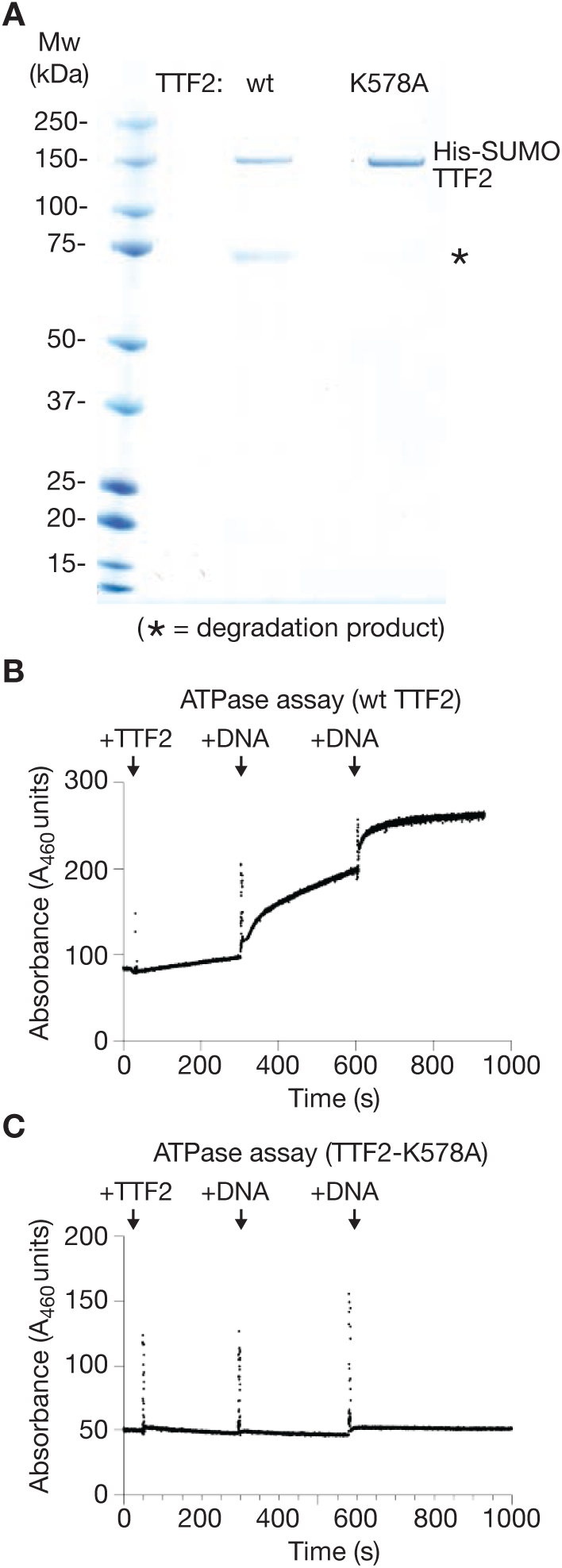
The TTF2-K578A mutant is defective in DNA-dependent ATPase activity. **(A)** Recombinant mouse TTF2 was purified from *E. coli* cells, together with the TTF2-K578A Walker A mutant, via an amino-terminal His-SUMO tag. Purified proteins were analysed by SDS-PAGE and Coomassie staining (the asterisk indicates a truncated form of TTF2). (**B**) ATPase activity of wild type recombinant TTF2 was monitored *in vitro* by a fluorescent assay (see Methods), upon addition of plasmid DNA at the indicated times. (**C**) Analogous assay with TTF2-K578A.

**Figure S11.**
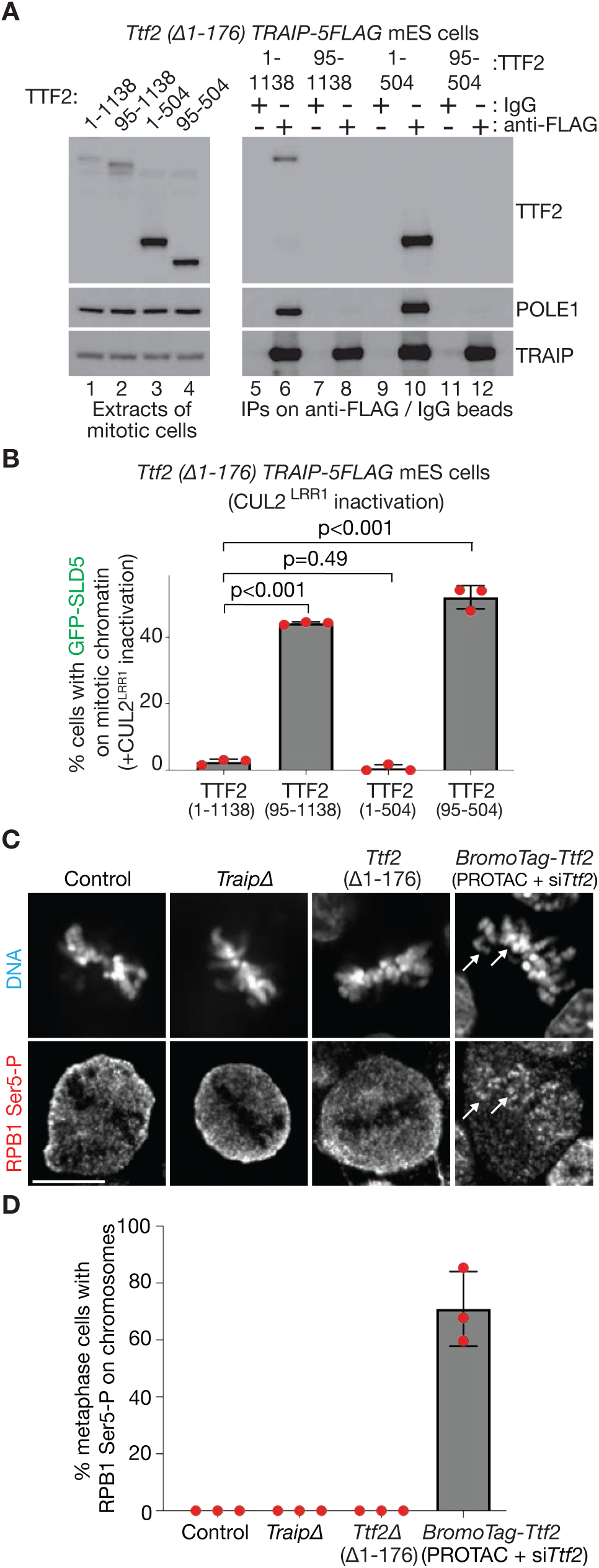
The TTF2 Zinc fingers are required for the mitotic TRAIP-TTF2-Polε complex & for mitotic CMG disassembly but are dispensable for unloading of RNA polymerase II from mitotic chromosomes. **(A)** *Ttf2(Δ1-176) Traip-5FLAG* cells, expressing the indicated versions of TTF2 as a second copy, were arrested in mitosis with 5 µM STLC. Extracts were then incubated with magnetic beads coated with anti-FLAG antibodies or control IgG as shown, before immunoblotting of the indicated factors. (**B**) Mitotic CMG disassembly was assayed as in Figure 1D, for *Ttf2 (Δ1-176)* mouse ES cells expressing the indicated versions of TTF2. The mean values for three independent experiments are shown as red spots, whereas the histograms show the average of the mean values, with associated standard deviations. P-values were calculated using a one-way ANOVA followed by Dunnett’s multiple-comparison test. (**C**) The association of RNA Polymerase II with mitotic chromatin (white arrows) was assayed as in Figure 3I, for mouse ES cells of the indicated genotypes. ‘PROTAC’ = 250 nM AGB1 and ‘*siTtf2*’ = siRNA for *Ttf2*. The scale bar corresponds to 10 µm. (**D**) Quantification of the data from (C), as in Figure 3J.

**Figure S12.**
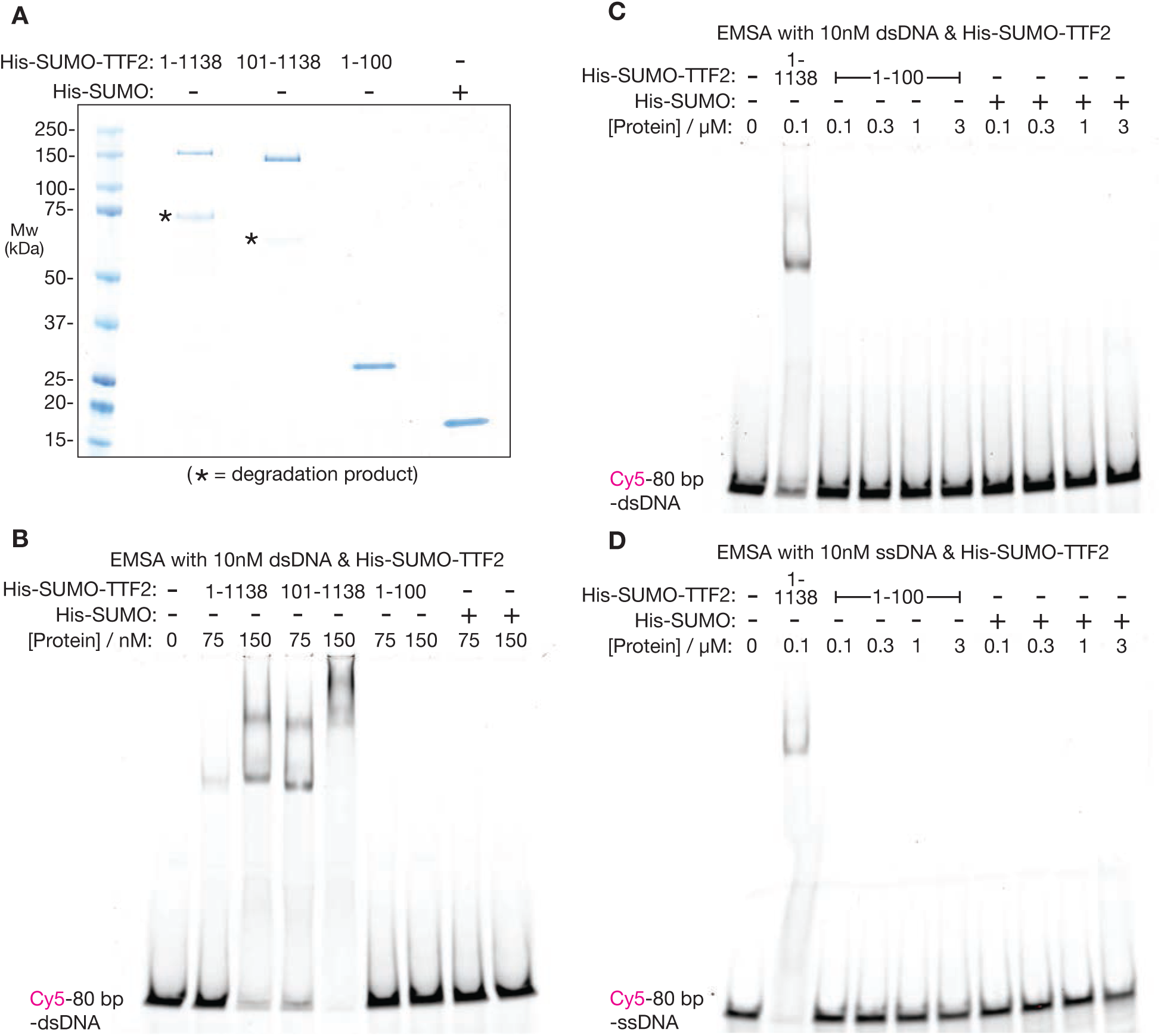
The TTF2 Zinc fingers do not bind DNA in vitro, in contrast to the TTF2 ATPase domain. **(A)** The indicated versions of TTF2 were expressed as His-SUMO fusions in E. coli as in Figure S10A, together with His-SUMO as a control protein. The purified proteins were resolved by SDS-PAGE and monitored by Coomassie staining (asterisks indicate truncated forms). (**B**-**D**) The ability of the indicated recombinant proteins to bind to double-stranded DNA (dsDNA) or single-stranded DNA (ssDNA) was monitored in Electrophoretic Mobility Shift Assays (EMSAs), with DNA substrates coupled to the far-red fluorescent dye Cyanine5 (Cy5). Reactions were resolved by PAGE before detection of Cy5 fluorescence.

**Figure S13.**
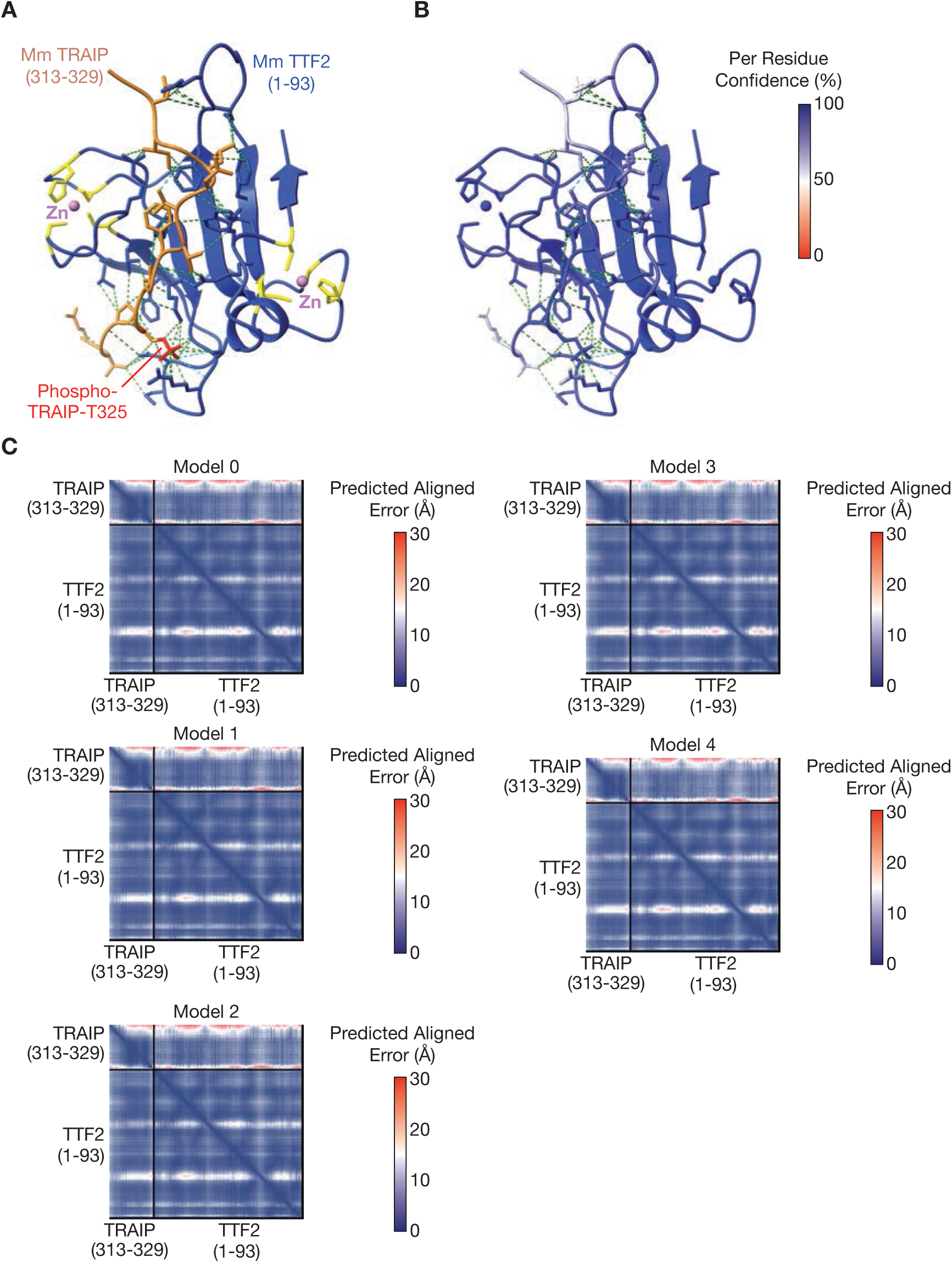
The TTF2 tandem Zinc finger domains are a phospho-receptor for TRAIP-T325. **(A)** AlphaFold 3 prediction of the TTF2 Zinc finger domains (blue) bound to TRAIP 313-329 (gold) containing phosphorylated T325 (red). The residues coordinating the two Zinc ions (magenta, ‘Zn’) are shown in yellow – see Figure S7A for more details. **(B)** The same prediction coloured by ‘Per Residue Confidence’. (**C**) Predicted Alignment Error (PAE) plots for the five predictions of TRAIP 313-329 bound to TTF2 1-93.

**Figure S14.**
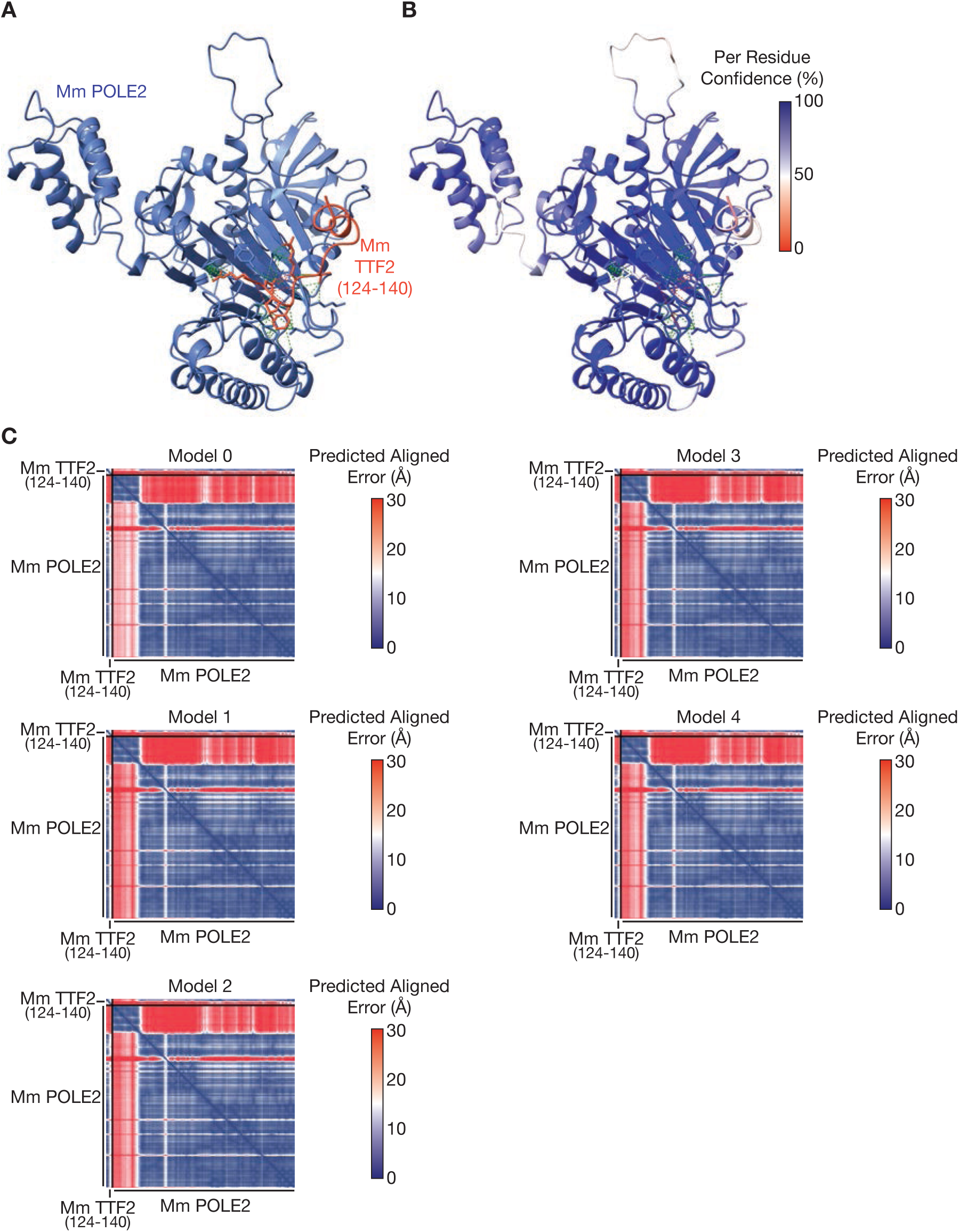
A short linear motif in TTF2 binds to the POLE2 subunit of DNA polymerase ε. **(A)** AlphaFold 3 prediction of mouse POLE2 (blue) bound to TTF2 124-140 (red). **(B)** The same prediction coloured by ‘Per Residue Confidence’. (**C**) Predicted Alignment Error (PAE) plots for the five predictions of mouse POLE2 bound to mouse TTF2 124-140.

**Figure S15.**
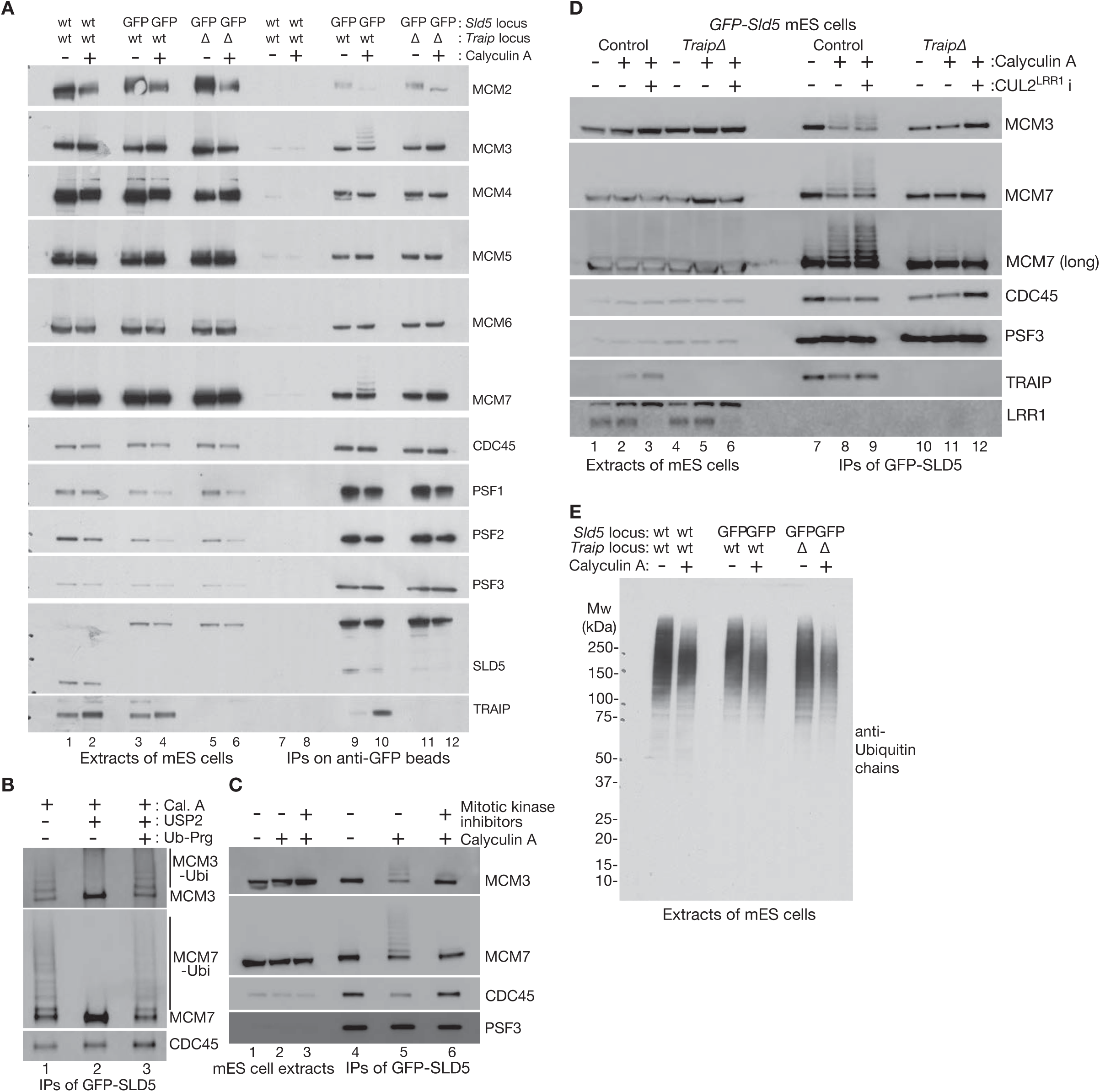
TRAIP-dependent mitotic ubiquitylation of the MCM3 and MCM7 subunits of CMG. **(A)** Cells of the indicated genotypes were treated as shown with 100 nM Calyculin A. Cell extracts were incubated with anti-GFP beads, before immunoblotting of the indicated factors. (**B**) The CMG helicase was isolated by immunoprecipitation on anti-GFP beads, from extracts of mouse ES cells treated with 100 nM Calyculin A. The beads were then incubated as shown with 1 µM USP2 deubiquitylase, and 16 µM propargylated ubiquitin (Ub-Prg) as a deubiquitylase inhibitor, for 20 minutes at 37°C. The indicated factors were detected by immunoblotting. (**C**) Mouse ES cells were treated with 10 µM each of RO-3306 (CDK1 inhibitor), VX-680 (AURORA kinase inhibitor) and BI-2536 (PLK1 inhibitor) for one hour, before addition of 100 nM Calyculin A for one hour as shown. The CMG helicase and associated factors were isolated from extracts as above and monitored by immunoblotting. (**D**) Mouse ES cells were treated as shown with inhibitors of CUL2^LRR1^ (see Methods) before treatment with 100 nM Calyculin A. CMG was isolated as above and monitored by immunoblotting. (**E**) Cells of the indicated genotype were treated as shown with 100 nM Calyculin A. Cell extracts were then analysed by SDS-PAGE and immunoblotting with an antibody to poly-ubiquitin chains (FK2 / UBCJ2).

**Figure S16.**
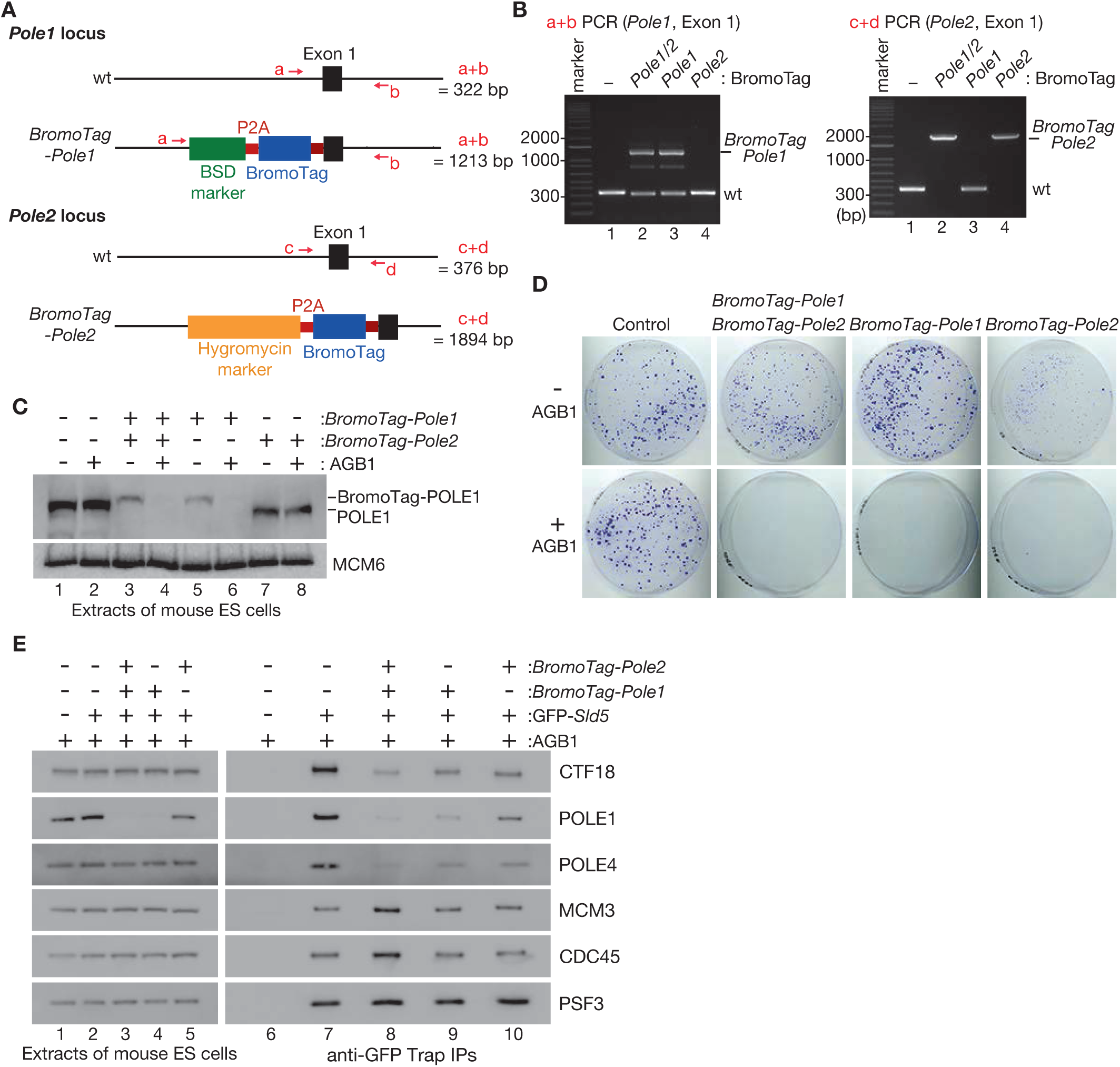
Inactivation of DNA polymerase ε by fusion of BromoTag to POLE1 and POLE2. **(A)** The BromoTag was inserted as shown into the *Pole1* and *Pole2* loci, preceded by marker genes encoding resistance to Blasticidin (BSD) or Hygromycin, and a ribosome skipping sequence (P2A). The location is shown of PCR primers that were used to check the loci, together with predicted size of the corresponding PCR products. (**B**) PCR analysis of mouse ES clones with BromoTag inserted at the *Pole1* locus, the *Pole2* locus, or both. For *Pole1*, one allele had insertion of the BromoTag cassette, whereas the other contained a small deletion that prevented expression of wild type protein. (**C**) Cells of the indicated genotypes were treated as shown with AGB1 for two hours. Cell extracts were resolved by SDS-PAGE and monitored by immunoblotting. (**D**) For each of the indicated genotypes, 500 cells were plated on 10 cm plates and grown for eight days in the presence or absence of 250 nM AGB1. Surviving colonies were fixed and stained with crystal violet. (**E**) Cells of the indicated genotypes were treated as shown with AGB1 for five hours. Cell extracts were incubated with anti-GFP antibodies before immunoblotting of the indicated factors. Displacement of the Pol ε subunit POLE4 and the Pol ε partner protein CTF18 from CMG was enhanced upon simultaneous degradation of both BromoTag-POLE1 and BromoTag-POLE2.

**Table S1.** Mass spectrometry data showing that TRAIP co-purifies with TTF2 and Pole from extracts of mitotic cells. Extracts of mouse ES cells were incubated with anti-FLAG beads before mass spectrometry analysis. The table shows selected factors that were enriched in the immunoprecipitates of TRAIP-5FLAG. This is the first of three independent experiments (the others are in Tables S2-S3).

| **Protein** | **Complex** | **Spectral counts**<br>**(Asyn. control**<br>**mES cells)** | **Spectral counts**<br>**(Asyn. TRAIP-**<br>**i5FLAG)** | **Spectral counts**<br>**(Mitotic control**<br>**mES cells)** | **Spectral counts**<br>**(Mitotic TRAIP-**<br>**i5FLAG)** |
| --- | --- | --- | --- | --- | --- |
| TRAIP<br>(53 kDa) | - | 136 | 2085 | 167 | 1945 |
| RAP80<br>(81) | BRCA1-A | 0 | 149 | 0 | 127 |
| ABRAXAS1<br>(46 kDa) | BRCA1-A | 0 | 115 | 0 | 106 |
| BABAM2<br>(44 kDa) | BRCA1-A | 0 | 74 | 0 | 65 |
| BABAM1<br>(37 kDa) | BRCA1-A | 0 | 49 | 0 | 34 |
| BRCC36<br>(33 kDa) | BRCA1-A | 0 | 88 | 0 | 53 |
| TTF2<br>(126 kDa) | - | 0 | 29 | 0 | 541 |
| POLE1<br>(262 kDa) | Pol e | 0 | 271 | 0 | 931 |
| POLE2<br>(59 kDa) | Pol e | 0 | 64 | 0 | 132 |
| POLE3<br>(17 kDa) | Pol e | 0 | 7 | 0 | 16 |
| POLE4<br>(12 kDa) | Pol e | 0 | 5 | 0 | 36 |

**Table S2.** Mass spectrometry data showing that TRAIP co-purifies with TTF2 and Pole from extracts of mitotic cells. Extracts of mouse ES cells were incubated with anti-FLAG beads before mass spectrometry analysis. The table shows selected factors that were enriched in the immunoprecipitates of TRAIP-i5FLAG. This is the second of three independent experiments (the others are in Tables S1 and S3).

| **Protein** | **Complex** | **Spectral counts**<br>**(Asyn. control**<br>**mES cells)** | **Spectral counts**<br>**(Asyn. TRAIP-**<br>**i5FLAG)** | **Spectral counts**<br>**(Mitotic control**<br>**mES cells)** | **Spectral counts**<br>**(Mitotic TRAIP-**<br>**i5FLAG)** |
| --- | --- | --- | --- | --- | --- |
| TRAIP<br>(53 kDa) | - | 0 | 1307 | 0 | 1910 |
| RAP80<br>(81) | BRCA1-A | 0 | 49 | 0 | 95 |
| ABRAXAS1<br>(46 kDa) | BRCA1-A | 0 | 52 | 0 | 67 |
| BABAM2<br>(44 kDa) | BRCA1-A | 0 | 36 | 4 | 46 |
| BABAM1<br>(37 kDa) | BRCA1-A | 0 | 14 | 0 | 29 |
| BRCC36<br>(33 kDa) | BRCA1-A | 0 | 15 | 0 | 53 |
| TTF2<br>(126 kDa) | - | 6 | 0 | 15 | 230 |
| POLE1<br>(262 kDa) | Pol e | 0 | 52 | 0 | 452 |
| POLE2<br>(59 kDa) | Pol e | 4 | 12 | 7 | 80 |
| POLE3<br>(17 kDa) | Pol e | 0 | 2 | 3 | 8 |
| POLE4<br>(12 kDa) | Pol e | 0 | 0 | 0 | 15 |

**Table S3.** Mass spectrometry data showing that TRAIP co-purifies with TTF2 and Pole from extracts of mitotic cells. Extracts of mouse ES cells were incubated with anti-FLAG beads before mass spectrometry analysis. The table shows selected factors that were enriched in the immunoprecipitates of TRAIP-i5FLAG. This is the third of three independent experiments (the others are in Tables S1-S2).

| **Protein** | **Complex** | **Spectral counts**<br>**(Asyn. control**<br>**mES cells)** | **Spectral counts**<br>**(Asyn. TRAIP-**<br>**i5FLAG)** | **Spectral counts**<br>**(Mitotic control**<br>**mES cells)** | **Spectral counts**<br>**(Mitotic TRAIP-**<br>**i5FLAG)** |
| --- | --- | --- | --- | --- | --- |
| TRAIP<br>(53 kDa) | - | 4 | 2126 | 0 | 1960 |
| RAP80<br>(81) | BRCA1-A | 0 | 75 | 0 | 98 |
| ABRAXAS1<br>(46 kDa) | BRCA1-A | 0 | 39 | 0 | 63 |
| BABAM2<br>(44 kDa) | BRCA1-A | 3 | 27 | 0 | 21 |
| BABAM1<br>(37 kDa) | BRCA1-A | 0 | 19 | 0 | 21 |
| BRCC36<br>(33 kDa) | BRCA1-A | 0 | 30 | 0 | 37 |
| TTF2<br>(126 kDa) | - | 0 | 8 | 7 | 201 |
| POLE1<br>(262 kDa) | Pol e | 0 | 135 | 0 | 464 |
| POLE2<br>(59 kDa) | Pol e | 0 | 30 | 0 | 76 |
| POLE3<br>(17 kDa) | Pol e | 4 | 6 | 0 | 11 |
| POLE4<br>(12 kDa) | Pol e | 0 | 6 | 0 | 15 |

**Supplementary Table S4.**
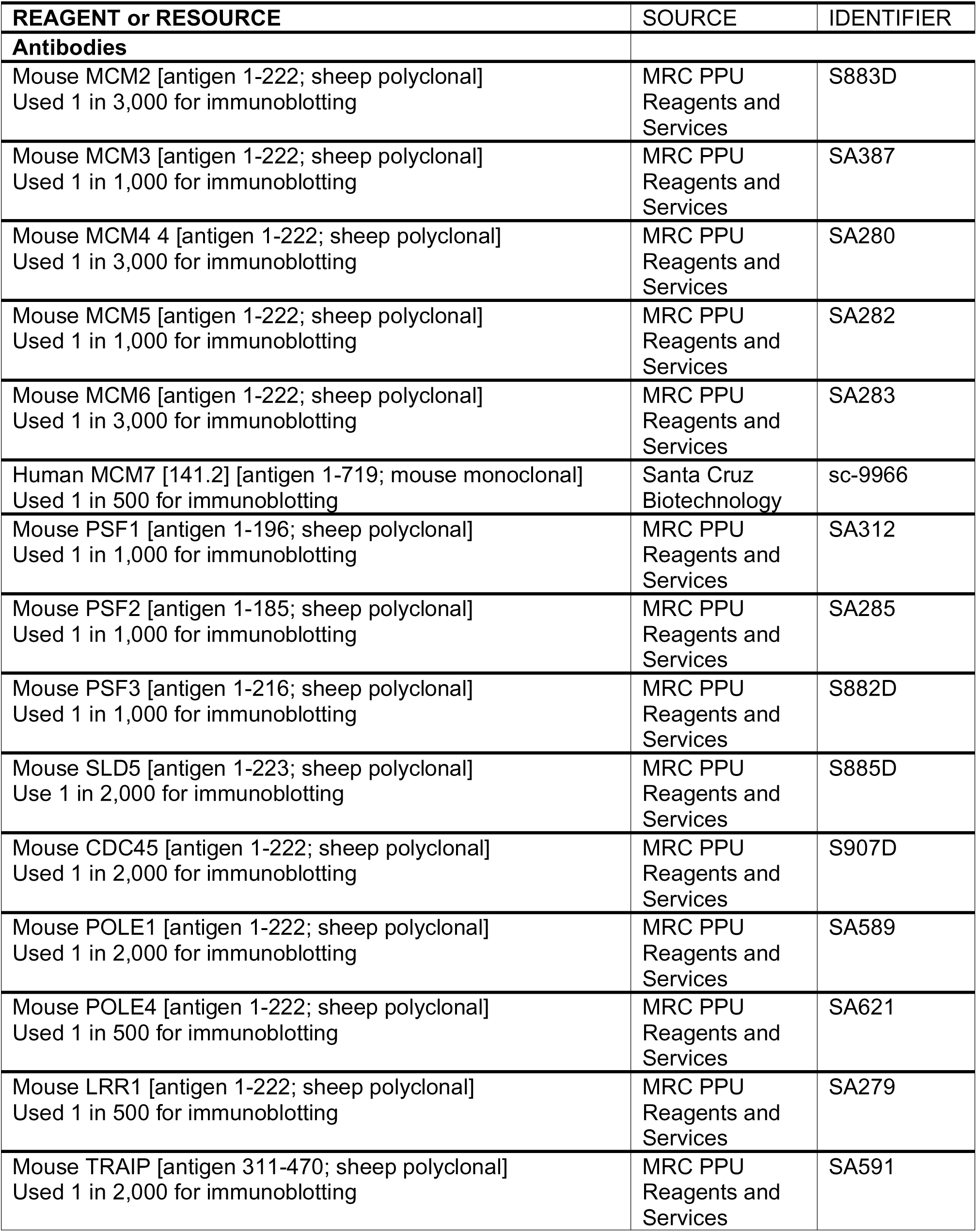
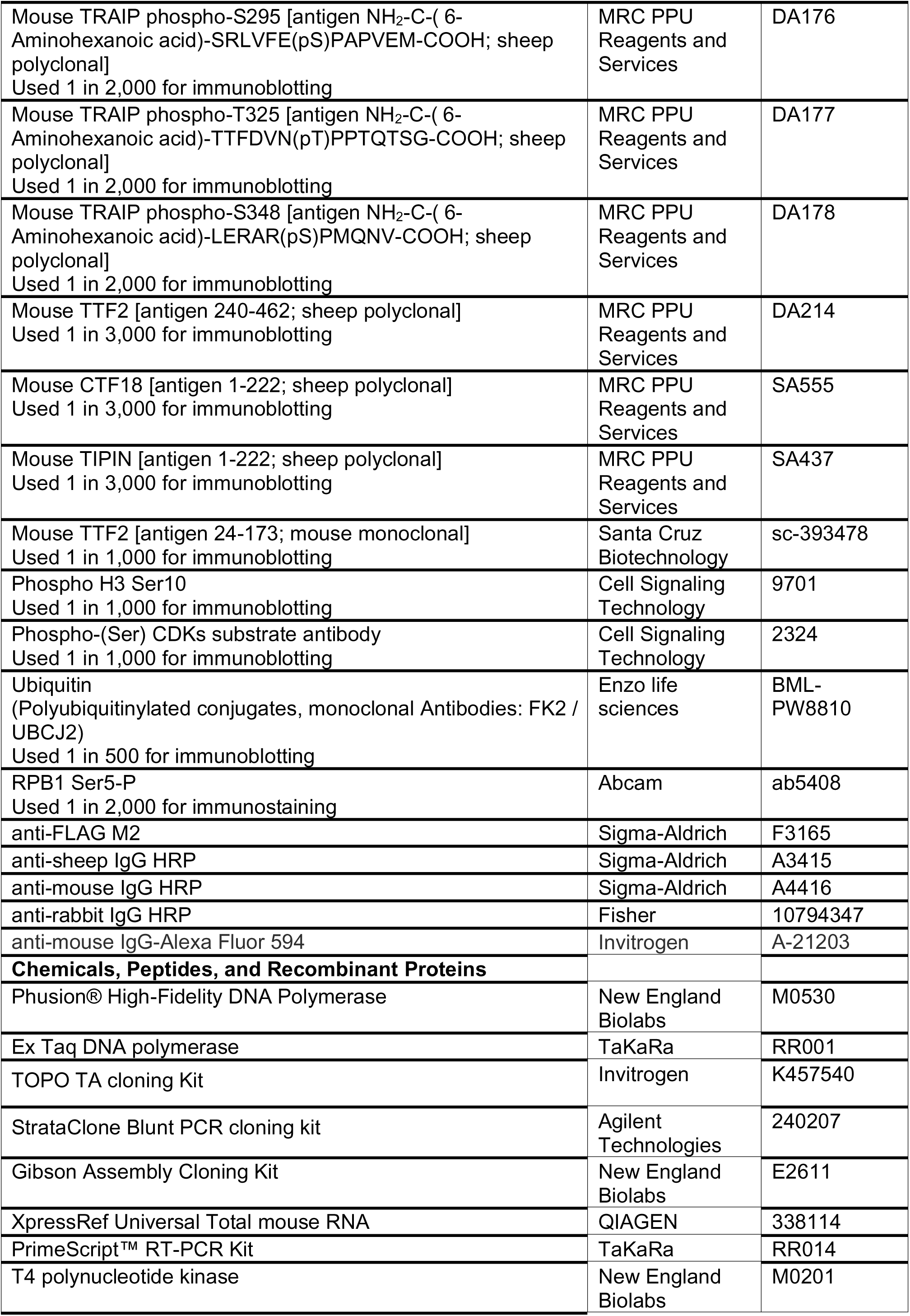
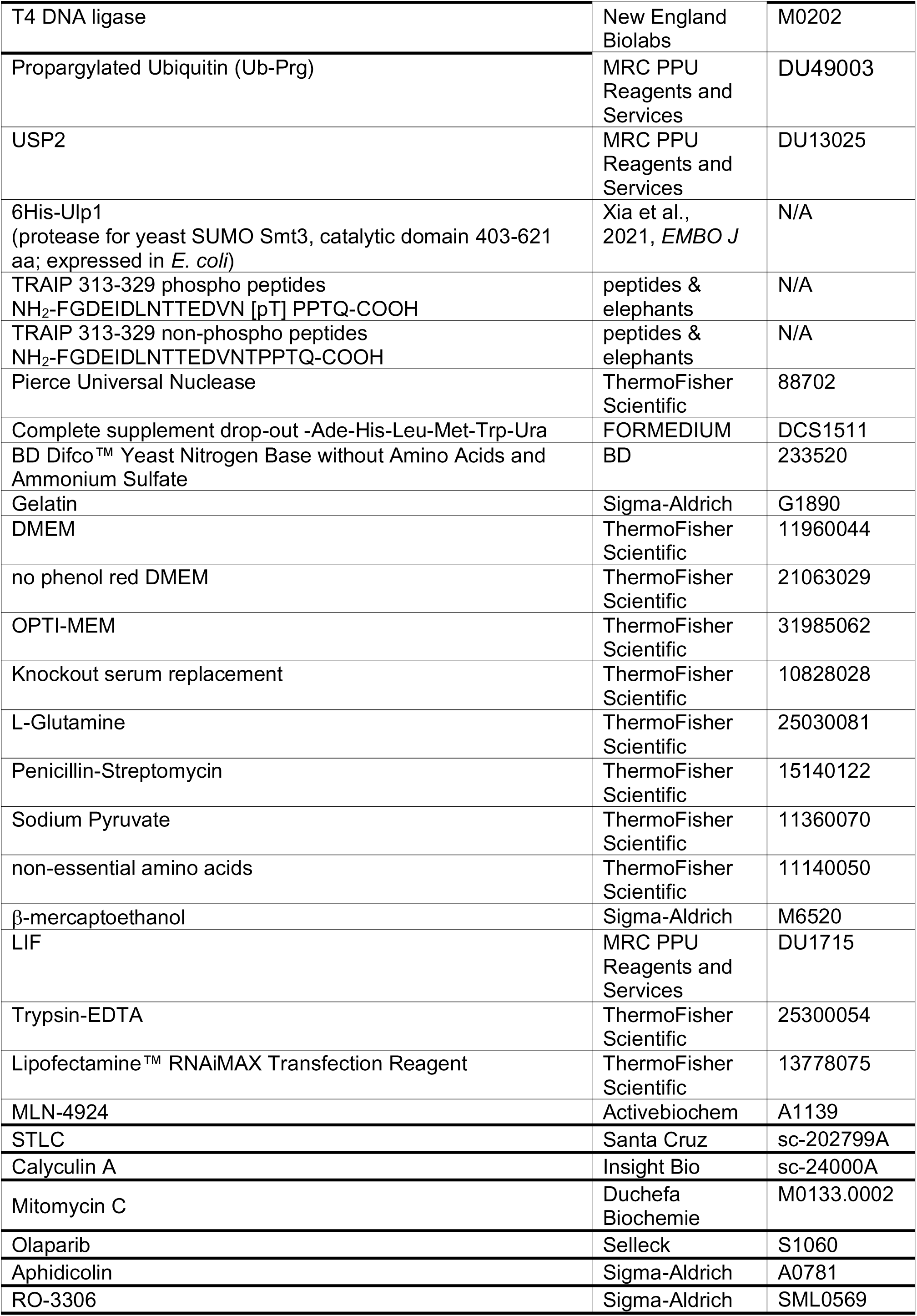
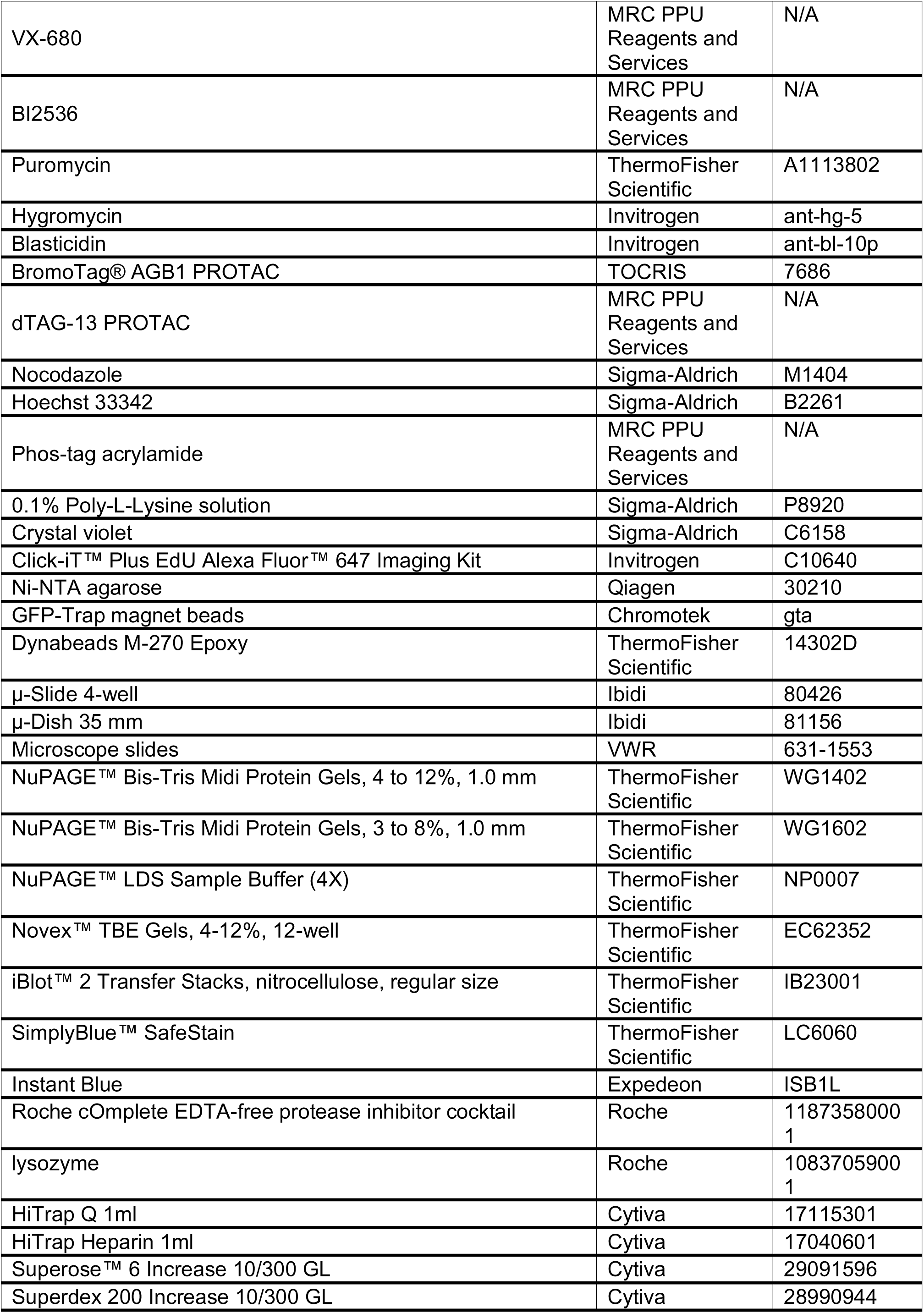
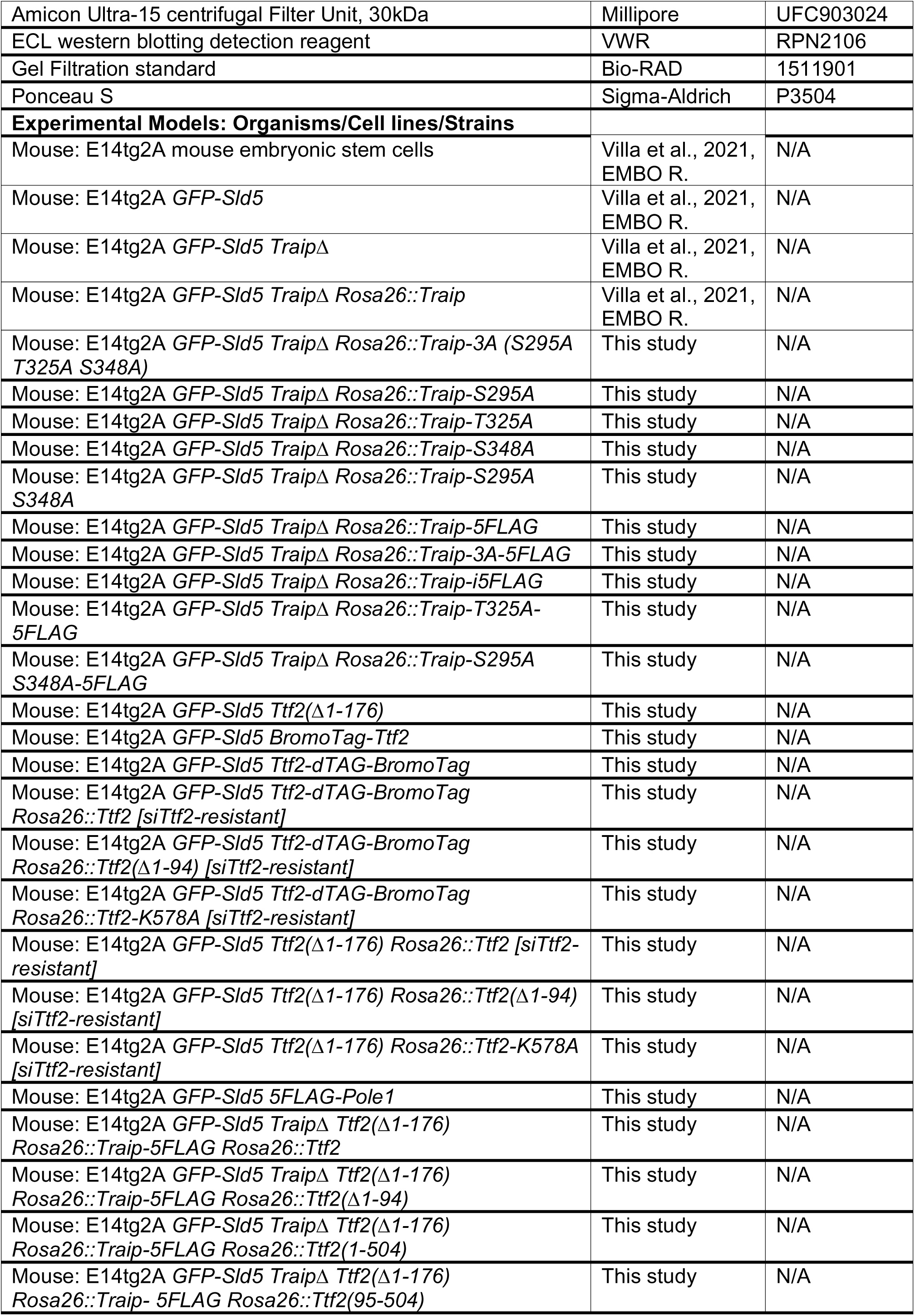
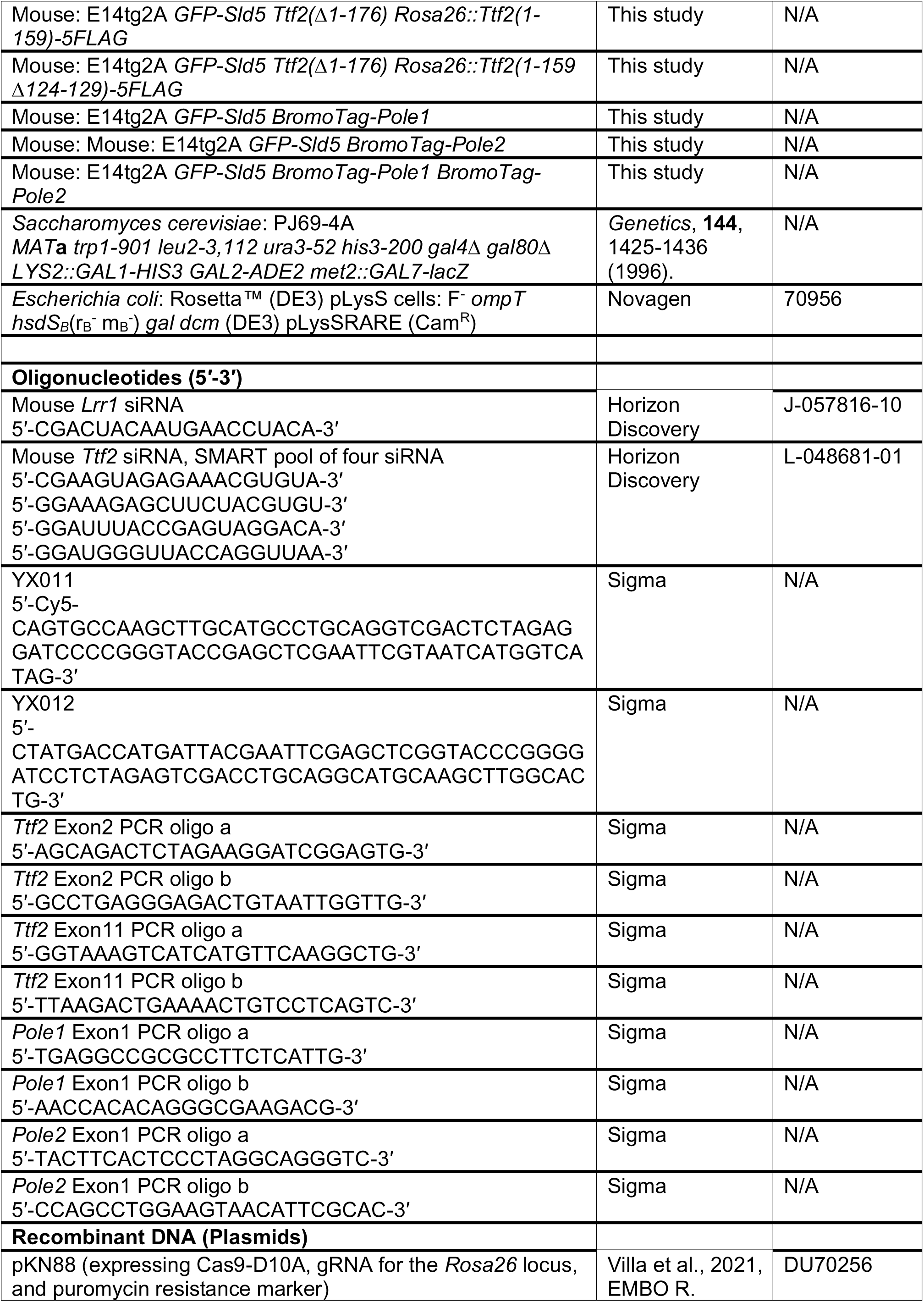
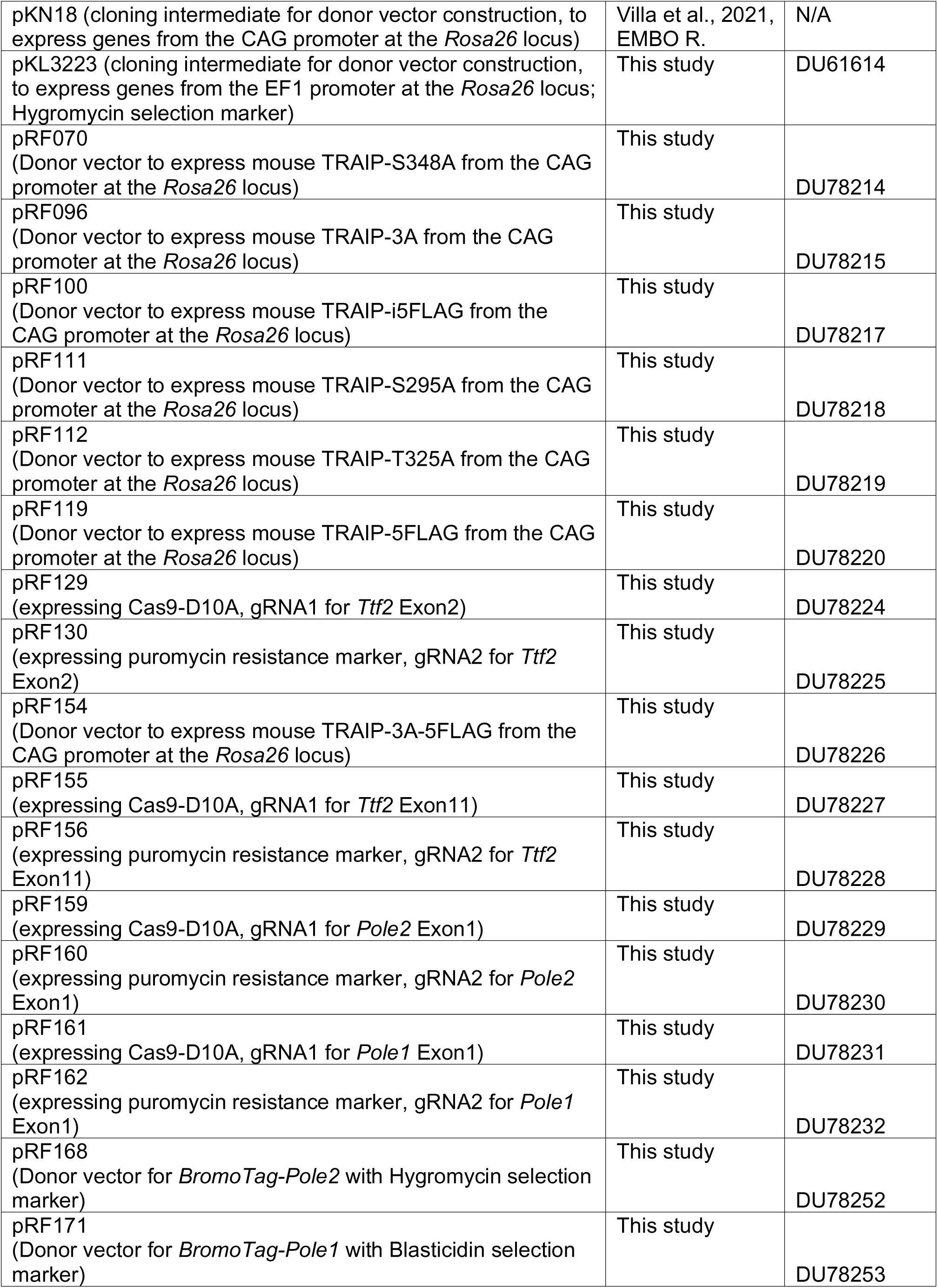
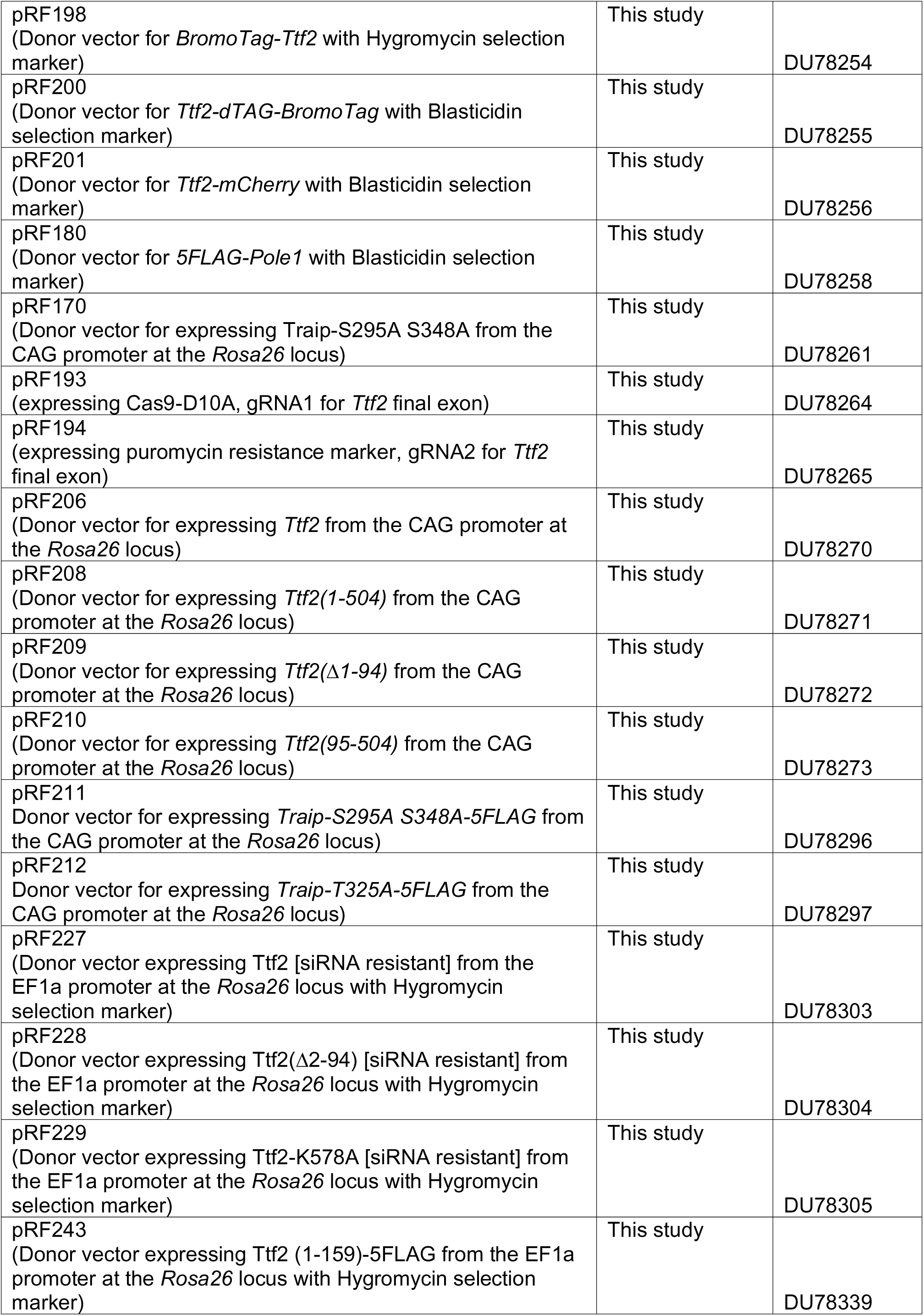
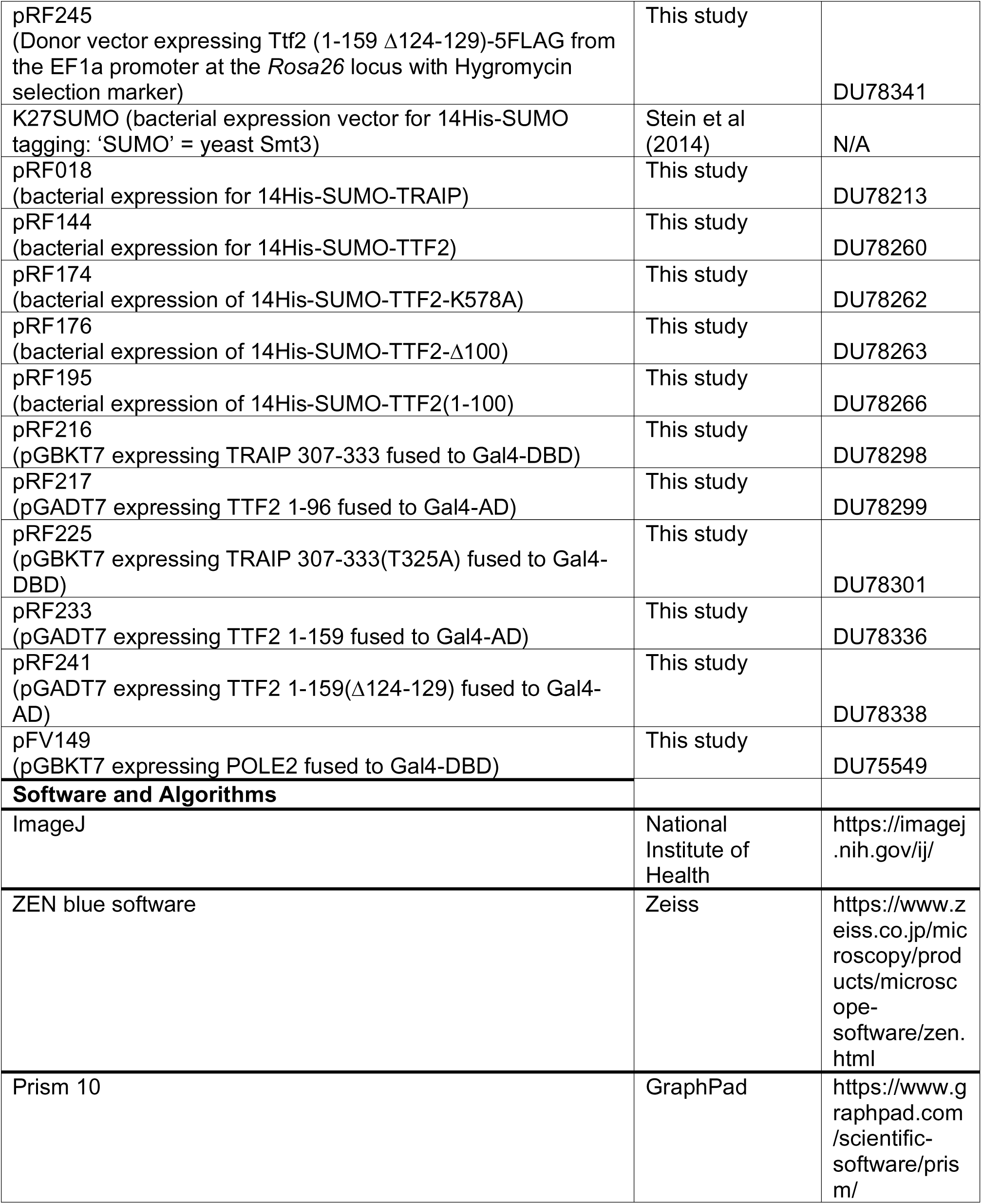
Reagents and resources used in this study.

